# The Genomic Formation of Human Populations in East Asia

**DOI:** 10.1101/2020.03.25.004606

**Authors:** Chuan-Chao Wang, Hui-Yuan Yeh, Alexander N Popov, Hu-Qin Zhang, Hirofumi Matsumura, Kendra Sirak, Olivia Cheronet, Alexey Kovalev, Nadin Rohland, Alexander M. Kim, Rebecca Bernardos, Dashtseveg Tumen, Jing Zhao, Yi-Chang Liu, Jiun-Yu Liu, Matthew Mah, Swapan Mallick, Ke Wang, Zhao Zhang, Nicole Adamski, Nasreen Broomandkhoshbacht, Kimberly Callan, Brendan J. Culleton, Laurie Eccles, Ann Marie Lawson, Megan Michel, Jonas Oppenheimer, Kristin Stewardson, Shaoqing Wen, Shi Yan, Fatma Zalzala, Richard Chuang, Ching-Jung Huang, Chung-Ching Shiung, Yuri G. Nikitin, Andrei V. Tabarev, Alexey A. Tishkin, Song Lin, Zhou-Yong Sun, Xiao-Ming Wu, Tie-Lin Yang, Xi Hu, Liang Chen, Hua Du, Jamsranjav Bayarsaikhan, Enkhbayar Mijiddorj, Diimaajav Erdenebaatar, Tumur-Ochir Iderkhangai, Erdene Myagmar, Hideaki Kanzawa-Kiriyama, Msato Nishino, Ken-ichi Shinoda, Olga A. Shubina, Jianxin Guo, Qiongying Deng, Longli Kang, Dawei Li, Dongna Li, Rong Lin, Wangwei Cai, Rukesh Shrestha, Ling-Xiang Wang, Lanhai Wei, Guangmao Xie, Hongbing Yao, Manfei Zhang, Guanglin He, Xiaomin Yang, Rong Hu, Martine Robbeets, Stephan Schiffels, Douglas J. Kennett, Li Jin, Hui Li, Johannes Krause, Ron Pinhasi, David Reich

**Affiliations:** Department of Anthropology and Ethnology, Institute of Anthropology, School of Sociology and Anthropology and State Key Laboratory of Cellular Stress Biology, School of Life Sciences, Xiamen University, Xiamen 361005, China; Department of Genetics, Harvard Medical School, Boston, Massachusetts 02115, USA; Max Planck Institute for the Science of Human History, 07745 Jena, Germany; MOE Key Laboratory of Contemporary Anthropology, Department of Anthropology and Human Genetics, School of Life Sciences, Fudan University, Shanghai 200438, China; School of Humanities, Nanyang Technological University, Nanyang 639798, Singapore; Scientific Museum, Far Eastern Federal University, 690950 Vladivostok, Russia; Key Laboratory of Biomedical Information Engineering of Ministry of Education, School of Life Science and Technology, Xi’an Jiaotong University, Xi’an 710049, China; School of Health Science, Sapporo Medical University, S1 W17, Chuo-ku, Sapporo, 060-8556, Japan; Department of Human Evolutionary Biology, Harvard Unviersity, Cambridge, MA 02138, USA; Department of Evolutionary Anthropology, University of Vienna, 1090 Vienna, Austria; Institute of Archaeology, Russian Academy of Sciences, Moscow, Russia; Department of Anthropology, Harvard University, Cambridge, Massachusetts 02138, USA; Department of Anthropology and Archaeology, National University of Mongolia, Ulaanbaatar 46, Mongolia; Institute of Archaeology, National Cheng Kung University, Tainan 701, Taiwan; Department of Anthropology, University of Washington, 314 Denny Hall, Seattle, USA; Broad Institute of Harvard and MIT, Cambridge, MA, 02142, USA; Howard Hughes Medical Institute, Harvard Medical School, Boston, MA 02115, USA; Institutes of Energy and the Environment, The Pennsylvania State University, University Park, PA 16802, USA; Department of Anthropology, Pennsylvania State University, University Park, PA 16802, USA; Institute of Archaeological Science, Fudan University, Shanghai 200433, China; School of Ethnology and Sociology, Minzu University of China, Beijing 100081, China; Museum of Archaeology and Ethnology of Institute of History, Far Eastern Branch of Russian Academic of Sciences, Vladivostok 690001, Russia; Institute of Archaeology and Ethnography, Siberian Branch of Russian Academy of Sciences, Novosibirsk 630090, Russia; Department of Archeology, Ethnography and Museology, Altai State University, Barnaul, Altaisky Kray 656049, Russia; Shaanxi Provincial Institute of Archaeology, Xi’an 710054, China; College of Cultural Heritage, Northwest University, Xi’an 710069, China; Xi’an AMS Center, Institute of Earth Environment, Chinese Academy of Sciences, Xi’an 710061, China; Research Center at the National Museum of Mongolia, Ulaanbaatar, Region of Sukhbaatar 14201, Mongolia; Department of Archaeology, Ulaanbaatar State University, Ulaanbaatar, Region of Bayanzurkh 13343, Mongolia; Department of Anthropology, National Museum of Nature and Science, Tsukuba City, Ibaraki Prefecture 305-0005, Japan; Archeological Center of Chiba City, Chiba 260-0814 Japan; Department of Archeology, Sakhalin Regional Museum, Yuzhno-Sakhalinsk, Russia; Department of Human Anatomy and Center for Genomics and Personalized Medicine, Guangxi Medical University, Nanning 530021, China; Key Laboratory for Molecular Genetic Mechanisms and Intervention Research on High Altitude Disease of Tibet Autonomous Region, Key Laboratory of High Altitude Environment and Gene Related to Disease of Tibet, Ministry of Education, School of Medicine, Xizang Minzu University, Xianyang 712082, Shaanxi, China; Guangxi Museum of Nationalities, Nanning 530028, Guangxi, China; Department of Biology, Hainan Medical University, Haikou 571199, Hainan, China; Department of Biochemistry and Molecular Biology, Hainan Medical University, Haikou 571199, Hainan, China; College of History, Culture and Tourism, Guangxi Normal University, Guilin 541001, China; Guangxi Institute of Cultural Relics Protection and Archaeology, Nanning 530003, Guangxi, China; Belt and Road Research Center for Forensic Molecular Anthropology, Key Laboratory of Evidence Science of Gansu Province, Gansu Institute of Political Science and Law, Lanzhou 730070, China; Department of Anthropology, University of California, Santa Barbara, CA 93106, USA

## Abstract

The deep population history of East Asia remains poorly understood due to a lack of ancient DNA data and sparse sampling of present-day people. We report genome-wide data from 191 individuals from Mongolia, northern China, Taiwan, the Amur River Basin and Japan dating to 6000 BCE – 1000 CE, many from contexts never previously analyzed with ancient DNA. We also report 383 present-day individuals from 46 groups mostly from the Tibetan Plateau and southern China. We document how 6000-3600 BCE people of Mongolia and the Amur River Basin were from populations that expanded over Northeast Asia, likely dispersing the ancestors of Mongolic and Tungusic languages. In a time transect of 89 Mongolians, we reveal how Yamnaya steppe pastoralist spread from the west by 3300-2900 BCE in association with the Afanasievo culture, although we also document a boy buried in an Afanasievo barrow with ancestry entirely from local Mongolian hunter-gatherers, representing a unique case of someone of entirely non-Yamnaya ancestry interred in this way. The second spread of Yamnaya-derived ancestry came via groups that harbored about a third of their ancestry from European farmers, which nearly completely displaced unmixed Yamnaya-related lineages in Mongolia in the second millennium BCE, but did not replace Afanasievo lineages in western China where Afanasievo ancestry persisted, plausibly acting as the source of the early-splitting Tocharian branch of Indo-European languages. Analyzing 20 Yellow River Basin farmers dating to ∼3000 BCE, we document a population that was a plausible vector for the spread of Sino-Tibetan languages both to the Tibetan Plateau and to the central plain where they mixed with southern agriculturalists to form the ancestors of Han Chinese. We show that the individuals in a time transect of 52 ancient Taiwan individuals spanning at least 1400 BCE to 600 CE were consistent with being nearly direct descendants of Yangtze Valley first farmers who likely spread Austronesian, Tai-Kadai and Austroasiatic languages across Southeast and South Asia and mixing with the people they encountered, contributing to a four-fold reduction of genetic differentiation during the emergence of complex societies. We finally report data from Jomon hunter-gatherers from Japan who harbored one of the earliest splitting branches of East Eurasian variation, and show an affinity among Jomon, Amur River Basin, ancient Taiwan, and Austronesian-speakers, as expected for ancestry if they all had contributions from a Late Pleistocene coastal route migration to East Asia.

---

East Asia, one of the oldest centers of animal and plant domestication, today harbors more than a fifth of the world’s human population, with present-day groups speaking languages representing eleven major families: Sino-Tibetan, Tai-Kadai, Austronesian, Austroasiatic, Hmong-Mien, Indo-European, Altaic (Mongolic, Turkic, and Tungusic), Koreanic, Japonic, Yukgahiric, and Chukotko-Kanchatkan^1^. The past 10,000 years have been a period of profound economic and cultural change in East Asia, but our current understanding of the genetic diversity, major mixture events, and population movements and turnovers during the transition from foraging to agriculture remains poor due to minimal sampling of the diversity of present-day people on the Tibetan Plateau and southern China^2^. A particular limitation has been a deficiency in ancient DNA data, which has been a powerful tool for discerning the deep history of populations in Western and Central Eurasia^3-8^.

We genotyped 383 present-day individuals from 46 populations indigenous to China (n=337) and Nepal (n=46) using the Affymetrix Human Origins array (Table S1 and Supplementary Information section 1). We also report genome-wide data from 191 ancient East Asians, many from cultural contexts for which there is no published ancient DNA data. From Mongolia we report 89 individuals from 52 sites dating between ∼6000 BCE to ∼1000 CE. From China we report 20 individuals from the ∼3000 BCE Neolithic site of Wuzhuangguoliang. From Japan we report 7 Jomon hunter-gatherers from 3500-1500 BCE. From the Russian Far East we report 23 individuals: 18 from the Neolithic Boisman-2 cemetery at ∼5000 BCE, 1 from the Iron Age Yankovsky culture at ∼1000 BCE, 3 from the Medieval Heishui Mohe and Bohai Mohe culture at ∼1000 CE; and 1 historic period hunter-gatherer from Sakhalin Island. From archaeological sites in Eastern Taiwan—the Bilhun site at Hanben on the main island and the Gongguan site on Green Island—we report 52 individuals from the Late Neolithic through the Iron Age spanning at least 1400 BCE – 600 CE.

For all but the Chinese samples we enriched the ancient DNA for a targeted set of about 1.2 million single nucleotide polymorphisms (SNPs)^4,9^, while for the Wuzhuangguoliang samples from China we used exome capture (18 individuals) or shotgun sequencing (2 individuals) (Figure 1, Supplementary Data files 1 and 2 and Supplementary Information section 1). We performed quality control to test for contamination by other human sequences, assessed by the rate of cytosine to thymine substitution in the terminal nucleotide and polymorphism in mitochondrial DNA sequences^10^ as well as X chromosome sequences in males, and restricted analysis to individuals with minimal contamination^11^ (Online Table 1). We detected close kinship between individuals at the same site, including a Boisman nuclear family with 2 parents and 4 children (Table S2). We merged the new data with previously reported data: 4 Jomon individuals, 8 Amur River Basin Neolithic individuals from the Devil’s Gate site, 72 individuals from the Neolithic to the Iron Age in Southeast Asia, and 8 from Nepal^7,12-20^. We assembled 123 radiocarbon dates using bone from the individuals, of which 94 are newly reported (Online Table 3), and clustered individuals based on time period and cultural associations, then further by genetic cluster which in the Mongolian samples we designated by number (our group names thus have the format “<Country>_<Time Period>_<Genetic Cluster>_<Cultural Association If Any>“) (Supplementary Note, Table S1 and Online Table 1). We merged the data with previously reported data (Online Table 4).

**Figure 1:**
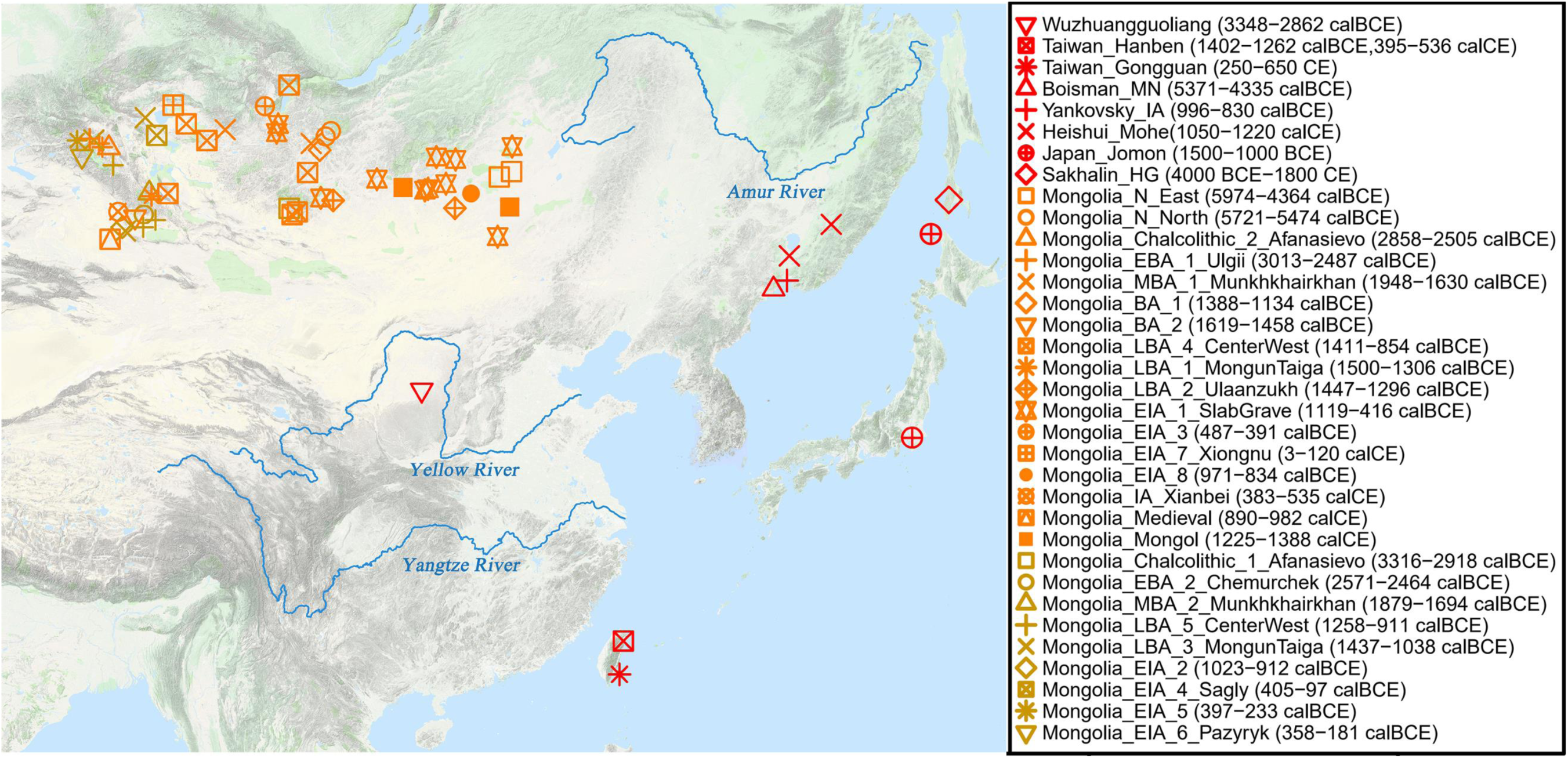
Geographical locations of newly reported ancient individuals. We use different colors for the two ancient Mongolia clusters. Detailed information are given in Table S1, Online Table 1 and Supplemental Experimental Procedures.

We carried out Principal Component Analysis (PCA) using smartpca^21^, projecting the ancient samples onto axes computed using present-day people. The analysis shows that population structure in East Asia is correlated with geographic and linguistic categories, albeit with important exceptions. Groups in Northwest China, Nepal, and Siberia deviate towards West Eurasians in the PCA (Supplementary Information section 2, Figure 2), reflecting multiple episodes of West Eurasian-related admixture that we estimate occurred 5 to 70 generations ago based on the decay of linkage disequilibrium^22^ (Table S3 and Table S4). East Asians with minimal proportions of West Eurasian-related ancestry fall along a gradient with three clusters at their poles. The “Amur Basin Cluster” correlates geographically with ancient and present-day populations living in the Amur River Basin, and linguistically with present-day indigenous people speaking Tungusic languages and the Nivkh. The “Tibetan Plateau Cluster” is most strongly represented in ancient Chokhopani, Mebrak, and Samzdong individuals from Nepal^15^ and in present-day people speaking Tibetan-Burman languages and living on the Tibetan Plateau. The “Southeast Asian Cluster” is maximized in ancient Taiwan groups and present-day people in Southeast Asia and southern parts of China speaking Austroasiatic, Tai-Kadai and Austronesian languages (Figure S1, Figure S2). Han are intermediate among these clusters, with northern Han projecting close to the Neolithic Wuzhuangguoliang individuals from northern China (Figure 2). We observe two genetic clusters within Mongolia: one falls closer to ancient individuals from the Amur Basin Cluster (‘East’ based on their geography), and the second clusters toward ancient individuals of the Afanasievo culture (‘West’), while a few individuals take intermediate positions between the two (Supplementary Information section 2).

**Figure 2:**
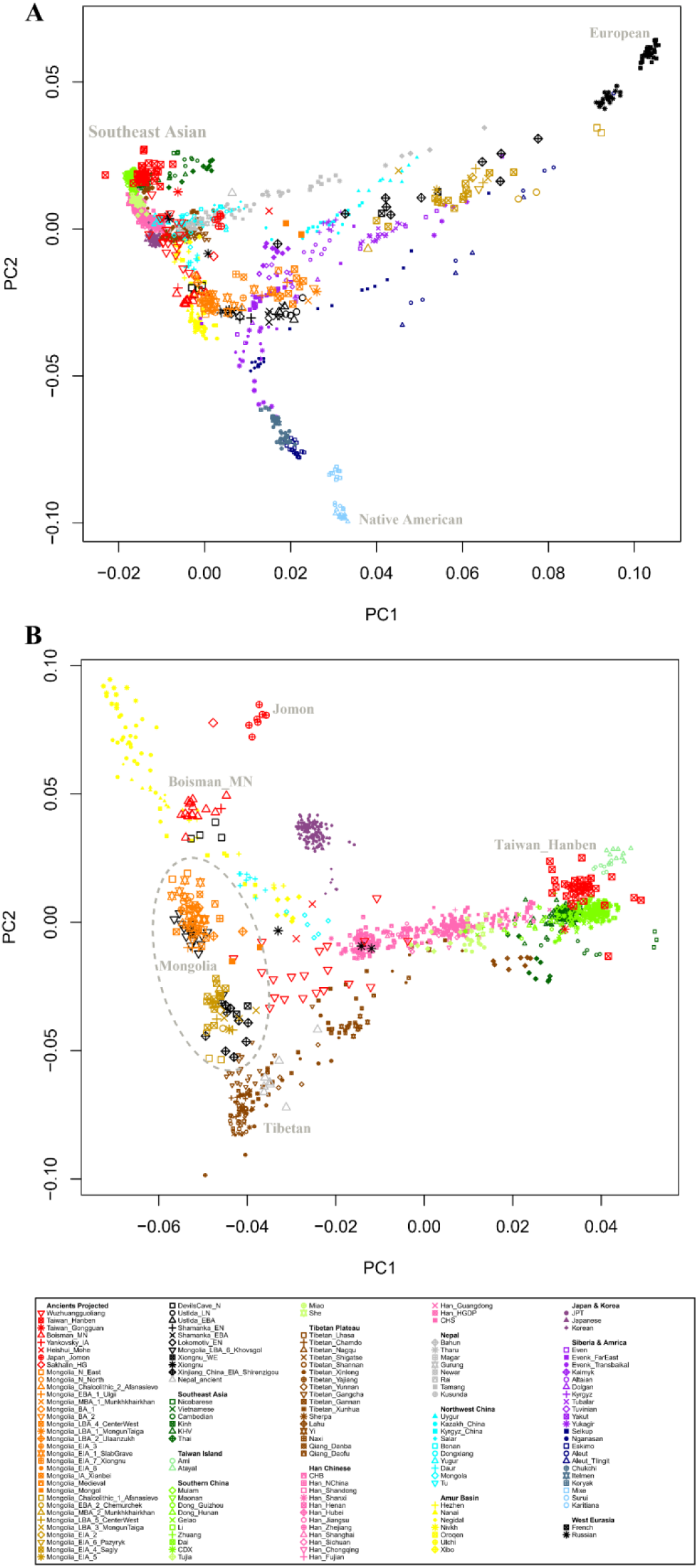
Principal Component Analysis (PCA). **(A)** Projection of ancient samples onto PCA dimensions 1 and 2 defined by East Asians, Europeans, Siberians and Native Americans. **(B)** Projection onto groups with the little West Eurasian mixture.

The three most ancient individuals of the Mongolia ‘East’ cluster are from the Kherlen River region of eastern Mongolia (Tamsag-Bulag culture) and date to 6000-4300 BCE (this places them in the Early Neolithic period, which in Northeast Asia is defined by the use of pottery and not by agriculture^23^). These individuals are genetically similar to previously reported Neolithic individuals from the cis-Baikal region and have minimal evidence of West Eurasian-related admixture as shown in PCA (Figure 2), *f*_*4*_-statistics and *qpAdm* (Table S5, Online Table 5, labeled as Mongolia_East_N). The other seven Neolithic hunter-gatherers from northern Mongolia (labeled as Mongolia_North_N) can be modeled as having 5.4% ± 1.1% ancestry from a source related to previously reported West Siberian Hunter-gatherers (WSHG)^8^ (Online Table 5), consistent with the PCA where they are part of an east-west Neolithic admixture cline in Eurasia with increasing proximity to West Eurasians in groups further west. Because of this ancestry complexity, we use the Mongolia_East_N individuals without significant evidence of West Eurasian-related admixture as reference points for modeling the East Asian-related ancestry in later groups (Online Table 5). The two oldest individuals from the Mongolia ‘West’ cluster have very different ancestry: they are from the Shatar Chuluu kurgan site associated with the Afanasievo culture, with one directly dated to 3316-2918 calBCE (we quote a 95% confidence interval here and in what follows whenever we mention a direct date), and are indistinguishable in ancestry from previously published ancient Afanasievo individuals from the Altai region of present-day Russia, who in turn are similar to previously reported Yamnaya culture individuals supporting findings that eastward Yamnaya migration had a major impact on people of the Afansievo culture^5,8^. All the later Mongolian individuals in our time transect were mixtures of Mongolian Neolithic groups and more western steppe-related sources, as reflected by statistics of the form *f*_*3*_ (X, Y; Later Mongolian Groups), which resulted in significantly negative Z scores (Z<−3) when Mongolia_East_N was used as X, and when Yamnaya-related Steppe populations, AfontovaGora3, WSHG, or European Middle/Late Neolithic or Bronze Age populations were used as Y (Table S6).

To quantify the admixture history of the later Mongolians, we again used *qpAdm*. A large number of groups could be modeled as simple two-way admixtures of Mongolia_East_N as one source (in proportions of 65-100%) and WSHG as the other source (in proportions of 0-35%), with negligible contribution from Yamnaya-related sources as confirmed by including Russia_Afanasievo and Russia_Sintashta groups in the outgroup set (Figure 3). The groups that fit this model were not only the two Neolithic groups (0-5% WSHG), but also the Early Bronze Age people from the Afanasievo Kurgak govi site (15%), the Ulgii group (28%), the main grouping of individuals from the Middle Bronze Age Munkhkhairkhan culture (33%), Late Bronze Age burials of the Ulaanzuukh type (6%), a combined group from the Center-West region (27%), the Mongun Taiga type from Khukh tolgoi (35%), and people of the Iron Age Slab Grave culture (9%). A striking finding in light of previous archaeological and genetic data is that the male child from Kurgak govi (individual I13957, skeletal code AT_629) has no evidence of Yamnaya-related ancestry despite his association with Afanasievo material culture (for example, he was buried in a barrow in the form of circular platform edged by vertical stone slabs, in stretched position on the back on the bottom of deep rectangular pit and with a typical Afanasievo egg-shaped vessel (Supplementary Note); his late Afanasievo chronology is confirmed by a direct radiocarbon date of 2858-2505 BCE^24^). This is the first known case of an individual buried with Afanasievo cultural traditions who is not overwhelmingly Yamnaya-related, and he also shows genetic continuity with an individual buried at the same site Kurgak govi 2 in a square barrow (individual I6361, skeletal code AT_635, direct radiocarbon date 2618-2487 BCE). We label this second individuals as having an Ulgii cultural association, although a different archaeological assessment associates this individual to the Afanasievo or Chemurchek cultures^25^, so it is possible that this provides a second example of Afanasievo material culture being adopted by individuals without any Yamnaya ancestry. The legacy of the Yamnaya-era spread into Mongolia continued in two individuals from the Chemurchek culture whose ancestry can be only modeled by using Afanasievo as one of the sources (49.0%±2.6%, Online Table 5). This model fits even when ancient European farmers are included in the outgroups, showning that if the long-distance transfer of West European megalithic cultural traditions to people of the Chemurchek culture that has been suggested in the archaeological literature occurred,^26^ it must have been through spread of ideas rather than through movement of people.

**Figure 3:**
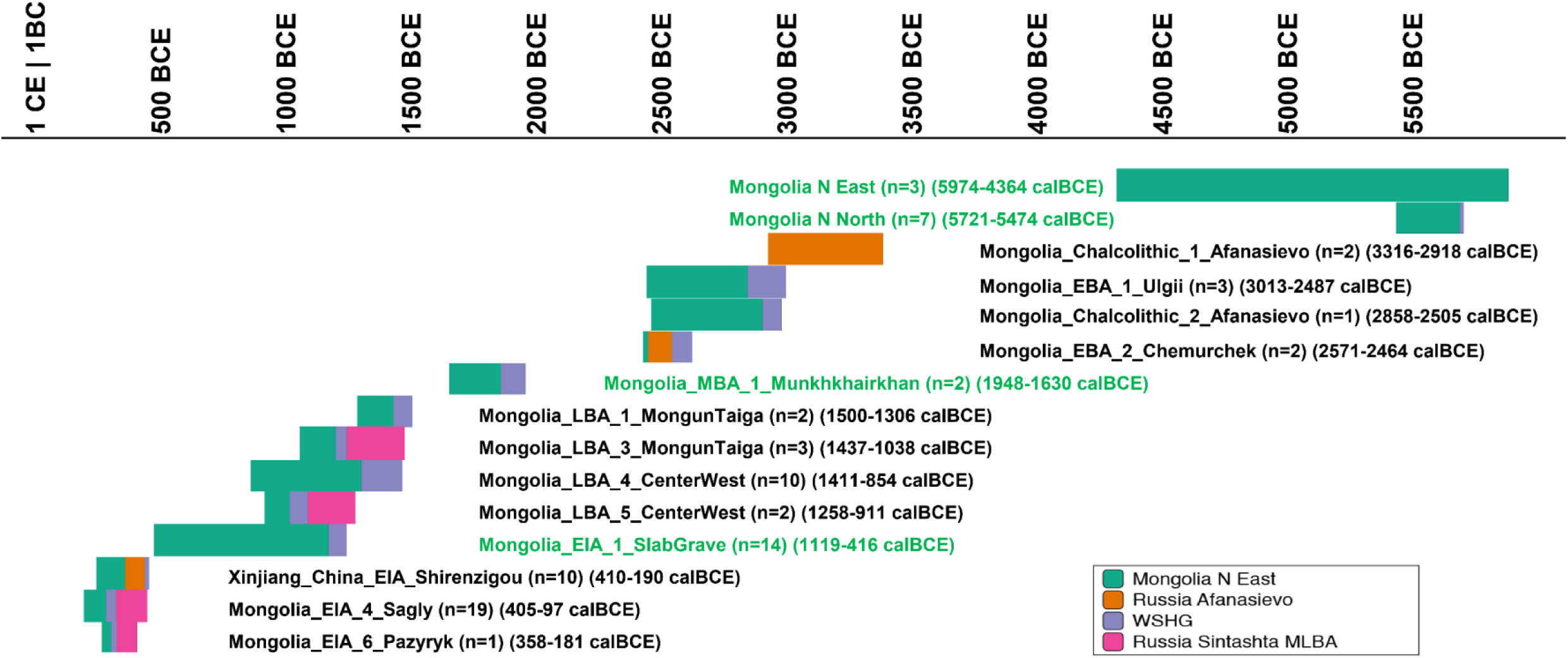
*qpAdm* modeling of ancestry change over time in Mongolia. We use Mongolia_East_N, Afanasievo, WSHG, and Sintashta_MLBA as sources, and for each combined archaeological and genetic grouping identify maximally parsimonious models (fewest numbers of sources) that fit with P>0.05 (Online Table 5). We plot results for groupings that give a unique parsimonious model, and include at least one individual with data that “PASS” at high quality and with a confident chronological assignment (Online Table 1). The bars show proportions of each ancestry source, and we also include time spans for the individuals in the cluster. Groupings that include more eastern individuals (longitude >102.7 degrees) are indicated in green and typically have very little Yamnaya-related admixture even at late dates.

Beginning in the Middle Bronze Age in Mongolia, there is no compelling evidence for a persistence of the Yamnaya-derivd lineages originally spread into the region with Afanasievo. Instead in the Late Bronze Age and Iron Age and afterward we have data from multiple Mongolian groups whose Yamnaya-related ancestry can only be modeled as deriving not from the initial Afanasievo migration but instead from a later eastward spread into Mongolia related to people of the Middle to Late Bronze Age Sintashta and Andronovo horizons who were themselves a mixture of ∼2/3 Yamnaya-related and 1/3 European farmer-related ancestry^5,7,8^. The Sintashta-related ancestry is detected in proportions of 5% to 57% in individuals from the Mongolia_LBA_6_Khovsgol (a culturally mixed group from the literature^14^), Mongolia_LBA_3_MongunTaiga, Mongolia_LBA_5_CenterWest, Mongolia_EIA_4_Sagly, Mongolia_EIA_6_Pazyryk, and Mongolia_Mongol groups, with the most substantial proportions of Sintashta-related ancestry always coming from western Mongolia (Figure 3, Online Table 5). For all these groups, the *qpAdm* ancestry models pass when Afanasievo is included in the outgroups while models with Afanasievo treated as the source with Sintashta more distantly related outgroups are all rejected (Figure 3, Online Table 5). Starting from the Early Iron Age, we finally detect evidence of gene flow in Mongolia from groups related to Han Chinese. Specifically, when Han are included in the outgroups, our models of mixtures in different proportions of Mongolia_East_N, Russia_Afanasievo, Russia_Sintashta, and WSHG continue to work for all Bronze Age and Neolithic groups, but fail for an Early Iron Age individual from Tsengel sum (Mongolia_EIA_5), and for Xiongnu and Mongols. When we include Han Chinese as a possible source, we estimate ancestry proportions of 20-40% in Xiongnu and Mongols (Online Table 5).

While the Afanasievo-derived lineages are consistent with having largely disappeared in Mongolia by the Late Bronze Age when our data showed that later groups with Steppe pastoralist ancestry made an impact, we confirm and strengthen previous ancient DNA analysis suggesting that the legacy of this expansion persisted in western China into the Iron Age Shirenzigou culture (410-190 BCE)^27^. The only parsimonious model for this group that fits according to our criteria is a 3-way mixture of groups related to Mongolia_N_East, Russia_Afanasievo, WSHG. The only other remotely plausible model (although not formally a good fit) also requires Russia_Afanasievo as a source (Figure 3, Online Table 5). The findings of the original study that reported evidence that the Afanasievo spread was the source of Steppe ancestry in the Iron Age Shirenzigou have been questioned with the proposal of alternative models that use ancient Kazakh Steppe Herders from the site of Botai, Wusun, Saka and ancient Tibetans from the site of Mebrak^15^ in present-day Nepal as major sources for Steppe and East Asian-related ancestry^28^. However, when we fit these models with Russia_Afanasievo and Mongolian_East_N added to the outgroups, the proposed models are rejected (P-values between 10^−7^ and 10^−2^), except in a model involving a single low coverage Saka individual from Kazakhstan as a source (P=0.17, likely reflecting the limited power to reject models with this low coverage). Repeating the modeling using other ancient Nepalese with very similar genetic ancestry to that in Mebrak results in uniformly poor fits (Online Table 5). Thus, ancestry typical of the Afanasievo culture and Mongolian Neolithic contributed to the Shirenzigou individuals, supporting the theory that the Tocharian languages of the Tarim Basin—from the second-oldest-known branch of the Indo-European language family—spread eastward through the migration of Yamnaya steppe pastoralists to the Altai Mountains and Mongolia in the guise of the Afansievo culture, from where they spread further to Xinjiang^5,7,8,27,29,30^. These results are significant for theories of Indo-European language diversification, as they increase the evidence in favor of the hypothesis the branch time of the second-oldest branch in the Indo-European language tree occurred at the end of the fourth millennium BCE^27,29,30^.

The individuals from the ∼5000 BCE Neolithic Boisman culture and the ∼1000 BCE Iron Age Yankovsky culture together with the previously published ∼6000 BCE data from Devil’s Gate cave^19^ are genetically very similar, documenting a continuous presence of this ancestry profile in the Amur River Basin stretching back at least to eight thousand years ago (Figure 2 and Figure S2). The genetic continuity is also evident in the prevailing Y chromosomal haplogroup C2b-F1396 and mitochondrial haplogroups D4 and C5 of the Boisman individuals, which are predominant lineages in present-day Tungusic, Mongolic, and some Turkic-speakers. The Neolithic Boisman individuals shared an affinity with Jomon as suggested by their intermediate positions between Mongolia_East_N and Jomon in the PCA and confirmed by the significantly positive statistic *f*_*4*_ (Mongolia_East_N, Boisman; Mbuti, Jomon). Statistics such as *f*_4_ (Native American, Mbuti; Test East Asian, Boisman/Mongolia_East_N) show that Native Americans share more alleles with Boisman and Mongolia_East_N than they do with the great majority of other East Asians in our dataset (Table S5). It is unlikely that these statistics are explained by back-flow from Native Americans since Boisman and other East Asians share alleles at an equal rate with the ∼24,000-year-old Ancient North Eurasian MA1 who was from a population that contributed about 1/3 of all Native American ancestry^31^. A plausible explanation for this observation is that the Boisman/Mongolia Neolithic ancestry was linked (deeply) to the source of the East Asian-related ancestry in Native Americans^3,31^. We can also model published data from Neolithic and Early Bronze Age individuals around Lake Baikal^7^ as sharing substantial ancestry (77-94%) with the lineage represented by Mongolia_East_N, revealing that this type of ancestry was once spread over a wide region spanning across Lake Baikal, eastern Mongolia, and the Amur River Basin (Table S7). Some present-day populations around the Amur River Basin harbor large fractions of ancestry consistent with deriving from more southern East Asian populations related to Han Chinese (but not necessarily Han themselves) in proportions of 13-50%. We can show that this admixture occurred at least by the Early Medieval period because one Heishui_Mohe individual (I3358, directly dated to 1050-1220 CE) is estimated to have harbored more than 50% ancestry from Han or related groups (Table S8).

The Tibetan Plateau, with an average elevation of more than 4,000 meters, is one of the most extreme environments in which humans live. Archaeological evidence suggests two main phases for modern human peopling of the Tibetan Plateau. The first can be traced back to at least ∼160,000 years ago probably by Denisovans^32^ and then to 40,000-30,000 years ago as reflected in abundant blade tool assemblages^33^. However, it is only in the last ∼3,600 years that there is evidence for continuous permanent occupation of this region with the advent of agriculture^34^. We grouped 17 present-day populations from the highlands into three categories based on genetic clustering patterns (Figure S3): “Core Tibetans” who are closely related to the ancient Nepal individuals such as Chokopani with a minimal amount of admixture with groups related to West Eurasians and lowland East Asians in the last dozens of generations, “northern Tibetans” who are admixed between lineages related to Core Tibetans and West Eurasians, and “Tibeto-Yi Corridor” populations (the eastern edge of the Tibetan Plateau connecting the highlands to the lowlands) that includes not just Tibetan speakers but also Qiang and Lolo-Burmese speakers who we estimate using *qpAdm*^4,35^ have 30-70% Southeast Asian Cluster-related ancestry (Table S9). We computed *f*_*3*_ (Mbuti; Core Tibetan, non-Tibetan East Asian) to search for non-Tibetans that share the most genetic drift with Tibetans. Neolithic Wuzhuangguoliang, Han and Qiang appear at the top of the list (Table S10), suggesting that Tibetans harbor ancestry from a population closely related to Wuzhuangguoliang that also contributed more to Qiang and Han than to other present-day East Asian groups. We estimate that the mixture occurred 60-80 generations ago (2240-1680 years ago assuming 28 years per generation^36^ under a model of a single pulse of admixture (Table S11). This represents an average date and so only provides a lower bound on when these two populations began to mix; the start of their period of admixture could plausibly be as old as the ∼3,600-year-old date for the spread of agriculture onto the Tibetan plateau. These findings are therefore consistent with archaeological evidence that expansions of farmers from the Upper and Middle Yellow River Basin influenced populations of the Tibetan Plateau from the Neolithic to the Bronze Age as they spread across the China Central plain^37,38^, and with Y chromosome evidence that the shared common haplogroup Oα-F5 between Han and Tibetans coalesced to a common ancestry less than 5,800 years ago^39^.

In the south, we find that the ancient Taiwan Hanben and Gongguan culture individuals dating from at least a span of 1400 BCE – 600 CE are genetically most similar to present-day Austronesian speakers and ancient Lapita individuals from Vanuatu as shown in outgroup *f*_*3*_-statistics and significantly positive *f*_*4*_-statistics (Taiwan_Hanben/Gongguan, Mbuti; Ami/Atayal/Lapita, other Asians) (Table S8). The similarity to Austronesian-speakers is also evident in the Iron Age dominant paternal Y chromosome lineage O3a2c2-N6 and maternal mtDNA lineages E1a, B4a1a, F3b1, and F4b, which are widespread lineages among Austronesian-speakers^40,41^. We compared the present-day Austronesian-speaking Ami and Atayal of Taiwan with diverse Asian populations using statistics like *f*_*4*_ (Taiwan Iron Age/Austronesian, Mbuti; Asian1, Asian2). Ancient Taiwan groups and Austronesian-speakers share significantly more alleles with Tai-Kadai speakers in southern mainland China and in Hainan Island^42^ than they do with other East Asians (Table S8), consistent with the hypothesis that ancient populations related to present-day Tai-Kadai speakers are the source for the spread of agriculture to Taiwan island around 5000 years ago^43^. The Jomon share alleles at an elevated rate with ancient Taiwan individuals and Ami/Atayal as measured by statistics of the form *f*_*4*_ (Jomon, Mbuti; Ancient Taiwan/Austronesian-speaker, other Asians) compared with other East Asian groups, with the exception of groups in the Amur Basin Cluster (Table S8)^44^.

The Han Chinese are the world’s largest ethnic group. It has been hypothesized based on the archaeologically documented spread of material culture and farming technology, as well as the linguistic evidence of links among Sino-Tibetan languages, that one of the ancestral populations of the Han might have consisted of early farmers along the Upper and Middle Yellow River in northern China, some of whose descendants also may have spread to the Tibetan Plateau and contributed to present-day Tibeto-Burmans^45^. Archaeological and historical evidence document how during the past two millennia, the Han expanded south into regions inhabited by previously established agriculturalists^46^. Analysis of genome-wide variation among present-day populations has revealed that the Han Chinese are characterized by a “North-South” cline^47,48^, which is confirmed by our analysis. The Neolithic Wuzhuangguoliang, present-day Tibetans, and Amur River Basin populations, share significantly more alleles with Han Chinese compared with the Southeast Asian Cluster, while the Southeast Asian Cluster groups share significantly more alleles with the majority of Han Chinese groups when compared with the Neolithic Wuzhuangguoliang (Table S12, Table S13). These findings suggest that Han Chinese may be admixed in variable proportions between groups related to Neolithic Wuzhuangguoliang and people related to those of the Southeast Asian Cluster. To determine the minimum number of source populations needed to explain the ancestry of the Han, we used *qpWave*^4,49^ to study the matrix of all possible statistics of the form *f*_*4*_ (Han_1_, Han_2_; O_1_, O_2_), where “O_1_” and “O_2_” are outgroups that are unlikely to have been affected by recent gene flow from Han Chinese. This analysis confirms that two source populations are consistent with all of the ancestry in most Han Chinese groups (with the exception of some West Eurasian-related admixture that affects some northern Han Chinese in proportions of 2-4% among the groups we sampled; Table S14 and Table S15). Specifically, we can model almost all present-day Han Chinese as mixtures of two ancestral populations, in a variety of proportions, with 77-93% related to Neolithic Wuzhuangguoliang from the Yellow River basin, and the remainder from a population related to ancient Taiwan that we hypothesize was closely related to the rice farmers of the Yangtze River Basin. This is also consistent with our inference that the Yangtze River farmer related ancestry contributed nearly all the ancestry of Austronesian speakers and Tai-Kadai speakers and about 2/3 of some Austroasiatic speakers^17,20^ (Figure 4). A caveat is that there is a modest level of modern contamination in the Wuzhuangguoliang we use as a source population for this analysis (Online Table 1), but this would not bias admixture estimates by more than the contamination estimate of 3-4%. The average dates of West Eurasian-related admixture in northern Han Chinese populations Han_NChina and Han_Shanxi are 32-45 generations ago, suggesting that mixture was continuing at the time of the Tang Dynasty (618-907 CE) and Song Dynasty (960-1279 BCE) during which time there are historical records of integration of Han Chinese amd western ethnic groups, but this date is an average so the mixture between groups could have begun earlier.

**Figure 4:**
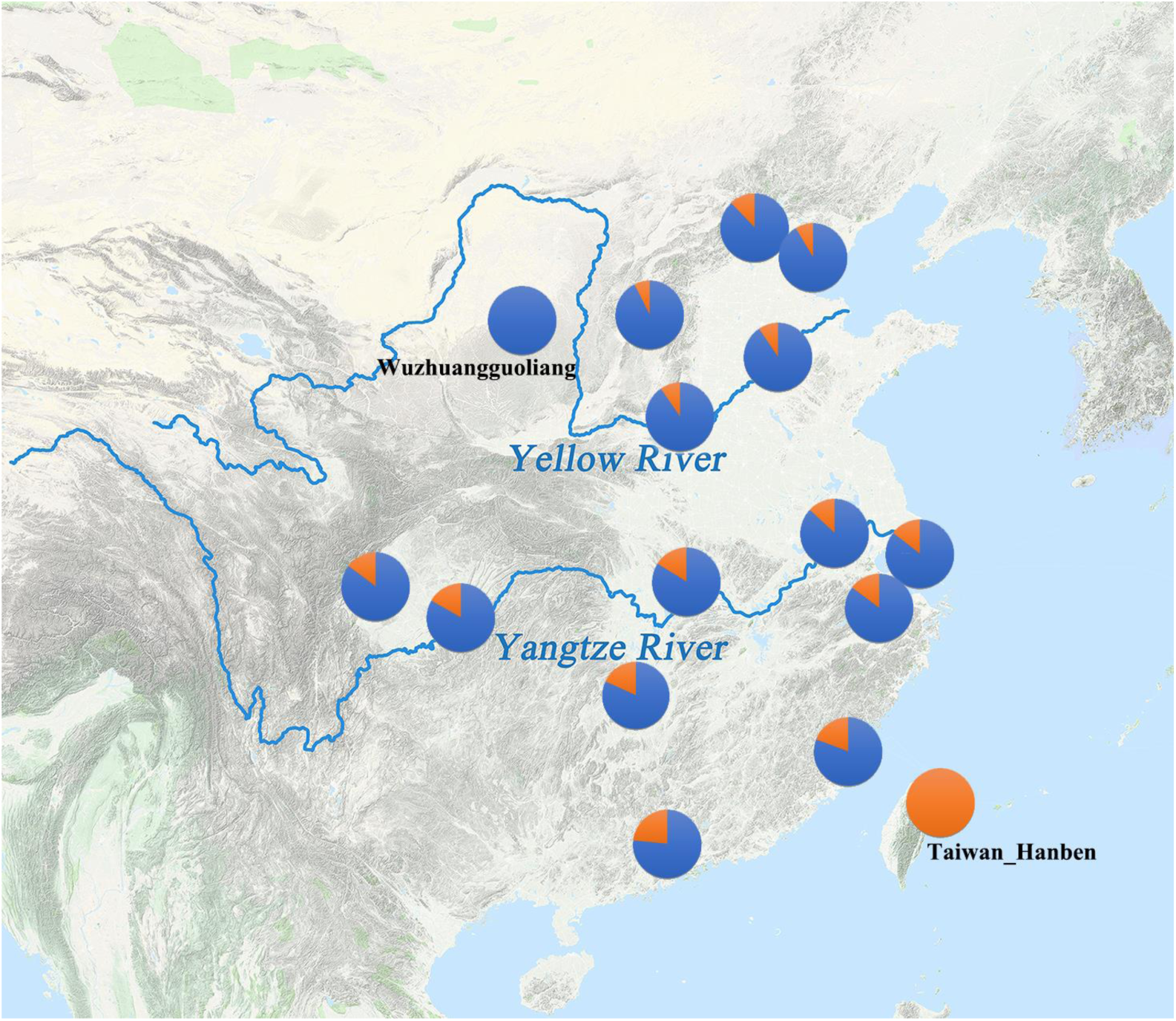
*qpAdm* modeling of Han Chinese cline. We used the ancient Wuzhuangguoliang as a proxy for Yellow River Farmers and Taiwan_Hanben as a proxy for Yangtze River Farmers related ancestry.

To obtain insight into the formation of present-day Japanese archipelago populations, we searched for groups that contribute most strongly to present-day Japanese through admixture *f*_*3*_-statistics. The most strongly negative signals come from mixtures of Han Chinese and ancient Jomon (*f*_*3*_(Japanese; Han Chinese, Jomon)) (Table S16). We can model present-day Japanese as two-way mixtures of 84.3% Han Chinese and 15.7% Jomon or 87.6% Korean and 12.4% Jomon (we cannot distinguish statistically between these two sources; Table S17 and Table S18). This analysis by no means suggests that the mainland ancestry in Japan was contributed directly by the Han Chinese or Koreans themselves, but does suggest that it is from an ancestral population related to those that contributed in large proportion to Han Chinese as well as to Koreans for which we do not yet have ancient DNA data.

We used *qpGraph*^35^ to explore models with population splits and gene flow, and tested their fit to the data by computing *f*_*2*_-, *f*_*3*_- and *f*_*4*_-statistics measuring allele sharing among pairs, triples, and quadruples of populations, evaluating fit based on the maximum |Z|-score comparing predicted and observed values. We further constrained the models by using estimates of the relative population split times between the selected pairs of populations based on the output of the MSMC software^50^. While admixture graph modeling based on allele frequency correlation statistics is not able to reject a model in which ancient Taiwan individuals and Boisman share substantial ancestry with each other more recently than either does with the ancestors of Chokopani and Core Tibetans, this model cannot be correct because our MSMC analysis reveals that Core Tibetans (closely related to Chokopani) and Ulchi (closely related to Boisman) share ancestry more recently in time on average than either does with Ami (related to Taiwan_Hanben). This MSMC-based constraint allowed us to identify a parsimonious working model for the deep history of key lineages discussed in this study (Supplementary Information section 3: *qpGraph* Modeling). Our fitted model (Figure 5), suggests that much of East Asian ancestry today can be modelled as derived from two ancient populations: one from the same lineage as the approximately ∼40,000-year-old Tianyuan individual and the other more closely related to Onge, with groups today having variable proportions of ancestry from these two deep sources. In this model, the Mongolia_East_N and Amur River Basin Boisman related lineages derive the largest proportion of their ancestry from the Tianyuan-related lineage and the least proportion of ancestry from the Onge-related lineage compared with other East Asians. A sister lineage of Mongolia_East_N is consistent with expanding into the Tibetan Plateau and mixing with the local hunter-gatherers who represent an Onge-related branch in the tree. The Taiwan Hanben are well modelled as deriving about 14% of their ancestry from a lineage remotely related to Onge and the rest of their ancestry from a lineage that also contributed to Jomon and Boisman on the Tianyuan side, a scenario that would explain the observed affinity among Jomon, Boisman and Taiwan Hanben. We estimate that Jomon individuals derived 45% of their ancestry from a deep basal lineage on the Onge side. These results are consistent with the scenario a Late Pleistocene coastal route of human migration linking Southeast Asia, the Japanese Archipelago and the Russian Far East^51^. Due to the paucity of ancient genomic data from Upper Paleolithic East Asians, there are limited constraints at present for reconstructing the deep branching patterns of East Asian ancestral populations, and it is certain that this admixture graph is an oversimplification and that additional features of deep population relationships will be revealed through future work.

**Figure 5:**
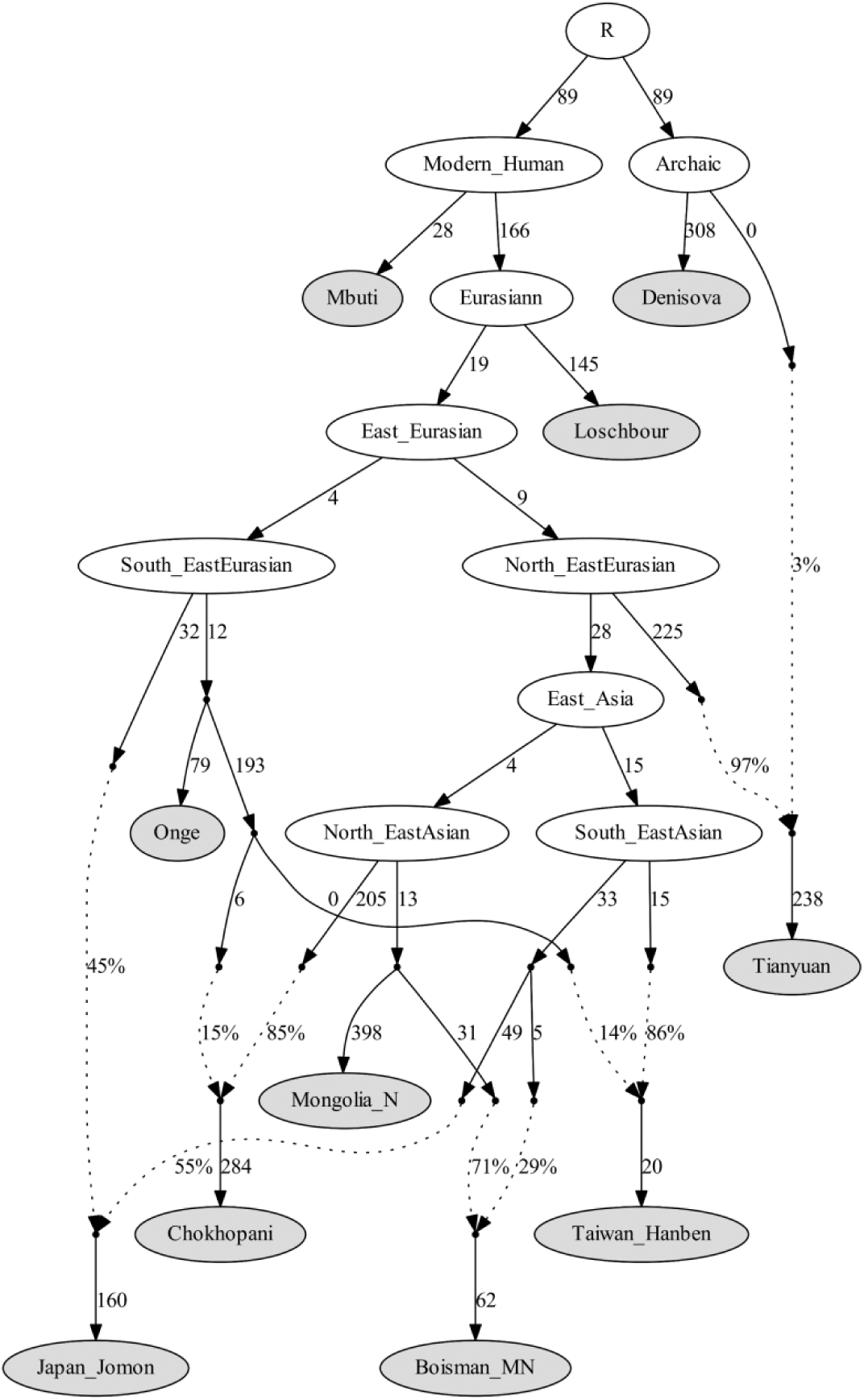
*qpGraph* modeling of a subset of East Asians. We used all available sites in the 1240K dataset, restricting to transversions only to replicate key results (Supplementary Information). We started with a skeleton tree that fits the data with Denisova, Mbuti, Onge, Tianyuan and Loschbour and one admixture event. We then grafted on Mongolia_East_N, Jomon, Taiwan_Hanben, Chokhopani, and Boisman in turn, adding them consecutively to all possible edges in the tree and retaining only graph solutions that provided no differences of |Z|>3 between fitted and estimated statistics. We used the MSMC relative population split time to constrain models (the maximum discrepancy for this model is |Z|=2.8). Drifts along edges are multiplied by 1000. Dashed lines represent admixture. Deep population splits are not well constrained due to a lack of data from Upper Paleolithic East Asians.

At the end of the last Ice Age, there were multiple highly differentiated populations in East as well as West Eurasia, and it is now clear that these groups mixed in both regions, instead of one population displacing the others. In West Eurasia, there were at least four divergent populations each as genetically differentiated from each other as Europeans and East Asians today (average F_ST_=0.10), which mixed in the Neolithic, reducing heterogeneity (average F_ST_=0.03) and mixed further in the Bronze Age and Iron Age to produce the present-relatively low differentiation that characterizes modern West Eurasia (average F_ST_=0.01)^52^. In East Eurasia, our study suggests an analogous process, with the differentiation characteristic of the Amur River Basin groups, Neolithic Yellow River farmers, and people related to those of the Taiwan Iron Age (average F_ST_=0.06 in our data) collapsing through mixture to today’s relatively low differentiation (average F_ST_=0.01-0.02) (Figure 6). A priority should be to obtain ancient DNA data for the hypothesized Yangtze River population (the putative source for the ancestry prevalent in the Southeast Asian Cluster of present-day groups), which should, in turn, make it possible to test and further extend these models, and in particular to understand if dispersals of people in Southeast Asia do or do not correlate to ancient movements of people.

**Figure 6.**
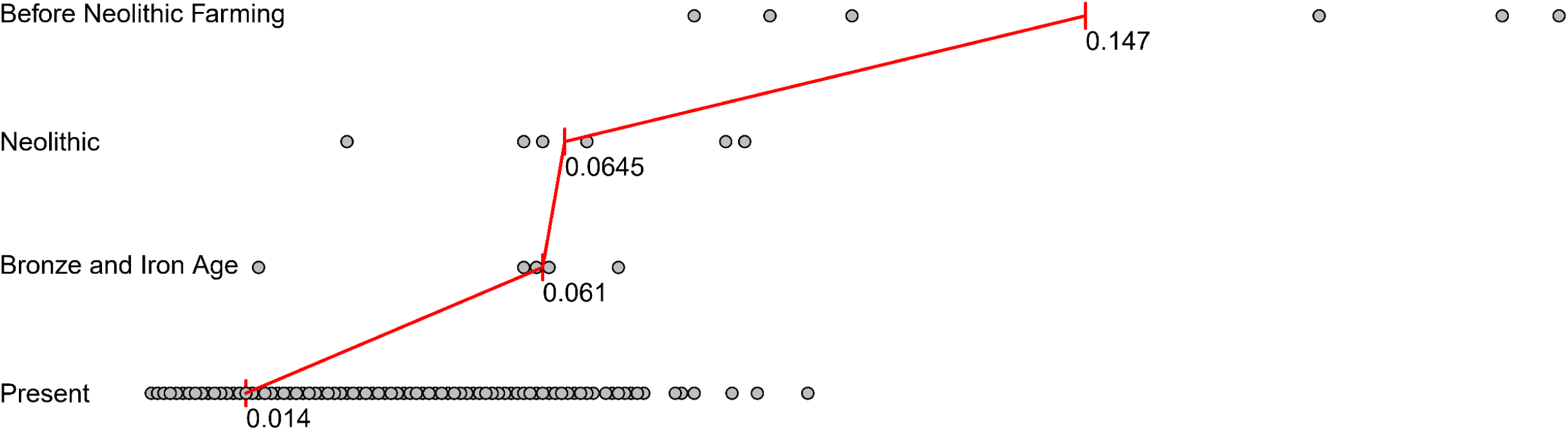
Homogenization of East Asian populations through mixture. Pairwise F_ST_ distribution among populations belonging to four time slices in East Asia; the median (red) of F_ST_ is shown.

## Supporting information

Supplementary Information sections 1-3

## Methods

### Ancient DNA laboratory work

All samples except those from Wuzhuangguoliang were prepared in dedicated clean room facilities at Harvard Medical School, Boston, USA. Online Table 2 lists experimental settings for each sample and library included in the dataset. Skeletal samples were surface cleaned and drilled or sandblasted and milled to produce a fine powder for DNA extraction^53,54^. We then either followed the extraction protocol by Dabney et al^55^ replacing the extender-MinElute-column assembly with the columns from the Roche High Pure Viral Nucleic Acid Large Volume Kit^56^ (manual extraction) or, for samples prepared later, used DNA extraction protocol based on silica beads instead of spin columns (and Dabney buffer) to allow for automated DNA purification^57^ (robotic extraction). We prepared individually barcoded double-stranded libraries for most samples using a protocol that included a DNA repair step with Uracil-DNA-glycosylase (UDG) treatment to cut molecules at locations containing ancient DNA damage that is inefficient at the terminal positions of DNA molecules (Online Table 1, UDG: “half”)^58^, or, without UDG pre-treatment (double stranded minus). For a few samples processed later, single stranded DNA libraries^59^ were prepared with USER (NEB) addition in the dephosphorylation step that results in inefficient uracil removal at the 5’end of the DNA molecules, and does not affect deamination rates at the terminal 3’ end^60^. We performed target enrichment via hybridization of these libraries with previously reported protocols^10^. We either enriched for the mitochondrial genome and 1.2M SNPs in two separate experiments or together in a single experiment. If split over two experiments, the first enrichment was for sequences aligning to mitochondrial DNA^58,61^ with some baits overlapping nuclear targets spiked in to screen libraries for nuclear DNA content. The second in-solution enrichment was for a targeted set of 1,237,207 SNPs that comprises a merge of two previously reported sets of 394,577 SNPs (390k capture)^4^ and 842,630 SNPs^9^. We sequenced the enriched libraries on an Illumina NextSeq500 instrument for 2×76 cycles (and both indices) or on Hiseq X10 instruments at the Broad Institute of MIT and Harvard for 2×101 cycles. We also shotgun sequenced each library for a few hundred thousand reads to assess the fraction of human reads.

Ancient DNA extractions of the Wuzhuangguoliang samples were performed in the clean room at Xi’an Jiaotong University and Xiamen University following the protocol by Rohland and Hofreiter^62^. Each sample extract was converted into double-stranded Illumina libraries following the manufacturer’s protocol (Fast Library Prep Kit, iGeneTech, Beijing, China). Sample-specific indexing barcodes were added to both sides of the fragments via amplification. Nuclear DNA capture was performed with AIExome Enrichment Kit V1 (iGeneTech, Beijing, China) according to the manufacturer’s protocol and sequenced on an Illumina NovaSeq instrument with 150 base pair paired-end reads. Sequences that did not perfectly match one of the expected index combinations were discarded.

For the AH1-7 and AH1-17 DNA extracts, we prepared whole genome sequencing libraries. The two DNA extracts were converted into barcoded Illumina sequencing libraries using commercially available library kits (NEBNext^®^ Ultra™ II DNA Library Prep Kit) and Illumina-specific primers^63^. DNA libraries were not treated with uracil-DNA-glycosylase (UDG) ^59^. We used a MinElute Gel Extraction Kit (Qiagen, Hilden, Germany) for purification. Two libraries were sequenced on a HiSeqX10 instrument (2×150 bp, PE) at the Novogene Sequencing Centre (Beijing, China). The base calling was performed using CASAVA software.

### Bioinformatic processing

For the sequencing data produced at Harvard Medical School, we used one of two pipelines (“pipeline 1” or “pipeline 2”; Online Table 2). An up-to-date description of both pipelines and analyses showing that the differences between them do not cause systematic bias in population genetic analysis can be found in Fernandes et al^64^. For both pipelines we began by de-multiplexed the data and assigning sequences to samples based on the barcodes and/or indices, allowing up to one mismatch per barcode or index. We trimmed adapters and restricted to fragments where the two reads overlapped by at least 15 nucleotides. In pipeline 1 we merged the sequences (allowing up to one mismatch) using a modified version of *Seqprep*^65^ where bases in the merged region are chosen based on highest quality in case of a conflict, and in pipeline 2 we used custom software (https://github.com/DReichLab/ADNA-Tools). For mitochondrial DNA analysis, we aligned the resulting merged sequences to the RSRS reference genome^66^ using *bwa* (version 0.6.1 for pipeline 1 and version 0.7.15 for pipeline 2)^67^, and removed duplicates with the same orientation, start and stop positions, and molecular barcodes. We determined mitochondrial DNA haplogroups using *HaploGrep2*^68^. We also analyzed the sequences to generate two assessments of ancient DNA authenticity. The first assessment estimated the rate of cytosine to thymine substitution in the final nucleotide, which is expected to be at least 3% at cytosines in libraries prepared with a partial UDG treatment protocol and at least 10% for untreated libraries (minus) and single stranded libraries; all libraries we analyzed met this threshold. The second assessment used *contamMix* (version 1.0.9 for pipeline 1 and 1.0.12 for pipeline 2)^10^ to determine the fraction of mtDNA sequences in an ancient sample that match the endogenous majority consensus more closely than a comparison set of 311 worldwide present-day human mtDNAs (Online Table 1). Computational processing of the sequence data from the whole genome was the same as the mtDNA enrichment except that the human genome (hg19) was used as the target reference. Due to the low coverage, diploid calling was not possible; instead, we randomly selected a single sequence covering every SNP position of interest (“pseudo-haploid” data) using custom software, only using nucleotides that were a minimum distance from the ends of the sequences to avoid deamination artifacts (https://github.com/DReichLab/adna-workflow). The coverages and numbers of SNPs covered at least once on the autosomes (chromosomes 1-22) are in Online Table 1.

For the sequencing data from the Wuzhuangguoliang samples, we clipped adaptors with *leehom*^69^ and then further processed using *EAGER*^70^, including mapping with *bwa* (v0.6.1)^67^ against the human genome reference GRCh37/hg19 (or just the mitochondrial reference sequence), and removing duplicate reads with the same orientation and start and end positions. To avoid an excess of remaining C-to-T and G-to-A transitions at the ends of the sequences, we clipped three bases of the ends of each read for each sample using trimBam (https://genome.sph.umich.edu/wiki/BamUtil:_trimBam). We generated pseudo-haploid calls by selecting a single read randomly for each individual using pileupCaller (https://github.com/stschiff/sequenceTools/tree/master/srcpileupCaller).

### Accelerator Mass Spectrometry Radiocarbon Dating

We generated 94 direct AMS (Accelerator Mass Spectrometry) radiocarbon (^14^C) dates as part of this study; 87 at Pennsylvania State University (PSU) and 7 at Poznan Radiocarbon Laboratory. The methods used at both laboratories are published, and here we summarize the methods from PSU. Bone collagen from petrous, phalanx, or tooth (dentine) samples was extracted and purified using a modified Longin method with ultrafiltration (>30kDa gelatin)^71^. If bone collagen was poorly preserved or contaminated we hydrolyzed the collagen and purified the amino acids using solid phase extraction columns (XAD amino acids)^72^. Prior to extraction we sequentially sonicated all samples in ACS grade methanol, acetone, and dichloromethane (30 minutes each) at room temperature to remove conservants or adhesives possibly used during curation. Extracted collagen or amino acid preservation was evaluated using crude gelatin yields (% wt), %C, %N and C/N ratios. Stable carbon and nitrogen isotopes were measured on a Thermo DeltaPlus instrument with a Costech elemental analyzer at Yale University. C/N ratios between 3.14 and 3.45 indicate that all radiocarbon dated samples are well preserved. All samples were combusted and graphitized at PSU using methods described in Kennett et al. 2017^71^. ^14^C measurements were made on a modified National Electronics Corporation 1.5SDH-1 compact accelerator mass spectrometer at either the PSUAMS facility or the Keck-Carbon Cycle AMS Facility. All dates were calibrated using the IntCal13 curve^73^ in OxCal v 4.3.2^74^ and are presented in calendar years BCE/CE.

### Y chromosomal haplogroup analysis

We performed Y-haplogroup determination by examining the state of SNPs present in ISOGG version 11.89 (accessed March 31, 2016) and our unpublished updated phylogeny.

### X-chromosome contamination estimates

We performed an X-chromosomal contamination test for the male individuals following an approach introduced by Rasmussen et al^75^ and implemented in the *ANGSD* software suite^11^. We used the “MoM” (Methods of Moments) estimates. The estimates for some males are not informative because of the limited number of X-chromosomal SNPs covered by at least two sequences, and hence we only report results for individuals with at least 200 SNPs covered at least twice. The estimated contamination rates for the male samples are low (Online Table 1). The contamination rates for all samples are quite low except those from Wuzhuangguoliang. We detected 3-6% contamination in the Wuzhuangguoliang samples, and restricted population genetic modeling analysis only to three males with 3-4% contamination.

### Data merging

We merged the data with previously published datasets genotyped on Affymetrix Human Origins arrays^3,35^, restricting to individuals with >95% genotyping completeness. We manually curated the data using ADMIXTURE^76^ and EIGENSOFT^21^ to identify samples that were outliers compared with other samples from their own populations. We removed seven individuals from subsequent analysis; the population IDs for these individuals are prefixed by the string “Ignore_” in the dataset we release, so users who wish to analyze these samples are still able to do so.

### Principal Components Analysis

We carried out principal components analysis in the *smartpca* program of EIGENSOFT^21^, using default parameters and the lsqproject: YES and numoutlieriter: 0 options.

### ADMIXTURE Analysis

We carried out ADMIXTURE analysis in unsupervised mode^76^ after pruning for linkage disequilibrium in PLINK^77^ with parameters --indep-pairwise 200 25 0.4 which retained 256,427 SNPs for Human Origin Dataset. We ran ADMIXTURE with default 5-fold cross-validation (--cv=5), varying the number of ancestral populations between K=2 and K=18 in 100 bootstraps with different random seeds.

### *f*-statistics

We computed *f*_3_-statistics and *f*_4_-statistics using ADMIXTOOLS^35^ with default parameters. We computed standard errors using a block jackknife^78^.

### F_ST_ computation

We estimated F_ST_ using EIGENSOFT^21^ with default parameters, inbreed: YES, and fstonly: YES. We found that the inbreeding corrected and uncorrected F_ST_ were nearly identical (within ∼0.001), and in this study, always analyzed uncorrected F_ST_.

### Admixture graph modeling

Admixture graph modeling was carried out with the *qpGraph* software as implemented in ADMIXTOOLS^35^ using Mbuti as an outgroup.

### Testing for the number of streams of ancestry

We used *qpWave*^4,35^ as implemented in ADMIXTOOLS to test whether a set of test populations is consistent with being related via *N* streams of ancestry from a set of outgroup populations.

### Inferring mixture proportions without an explicit phylogeny

We used *qpAdm*^4^ as implemented in ADMIXTOOLS to estimate mixture proportions for a *Test* population as a combination of *N* ‘reference’ populations by exploiting (but not explicitly modeling) shared genetic drift with a set of ‘Outgroup’ populations.

### Weighted linkage disequilibrium (LD) analysis

LD decay was calculated using ALDER^22^ to infer admixture parameters including dates and mixture proportions.

### MSMC

We used MSMC^50^ following the procedures in Mallick et al^79^ to infer cross-coalescence rates and population sizes among Ami/Atayal, Tibetan, and Ulchi.

### Kinship analysis

We used READ software^80^ as well as a custom method^81^ to determine genetic kinship between individual pairs.

### Data availability

The aligned sequences are available through the European Nucleotide Archive under accession number [to be made available on publication]. Genotype data used in analysis are available at https://reich.hms.harvard.edu/datasets. Any other relevant data are available from the corresponding author upon reasonable request.

## Acknowledgements

We thank David Anthony, Ofer Bar-Yosef, Katherine Brunson, Rowan Flad, Pavel Flegontov, Qiaomei Fu, Wolfgang Haak, Iosif Lazaridis, Mark Lipson, Iain Mathieson, Richard Meadow, Inigo Olalde, Nick Patterson, Pontus Skoglund, and Dan Xu for valuable conversations and critical comments. We thank Naruya Saitou and the Asian DNA Repository Consortium for sharing genotype data from present-day Japanese groups. We thank Toyohiro Nishimoto and Takashi Fujisawa from the Rebun Town Board of Education for providing the Funadomari Jomon samples, and Hideyo Tanaka and Watru Nagahara from the Archeological Center of Chiba City who are excavators of the Rokutsu Jomon site. The excavations at Boisman-2 site (Boisman culture), the Pospelovo-1 site (Yankovsky culture), and the Roshino-4 site (Heishui Mohe culture) were funded by the Far Eastern Federal University and the Institute of History Far Eastern Branch of the Russian Academu of Sciences, researches Pospelovo-1 funded by RFBR project number 18-09-40101. C.C.W was funded by the Max Planck Society, the National Natural Science Foundation of China (NSFC 31801040), the Nanqiang Outstanding Young Talents Program of Xiamen University (X2123302), and Fundamental Research Funds for the Central Universities (ZK1144). O.B. and Y.B. were funded by Russian Scientific Foundation grant 17-14-01345. H.M. was supported by the grant JSPS 16H02527. The research of M.R. and C.C.W has received funding from the European Research Council (ERC) under the European Union’s Horizon 2020 research and innovation programme (grant agreement No 646612) granted to M.R. The research of C.S. is supported by the Calleva Foundation and the Human Origins Research Fund. H.L was funded NSFC (91731303, 31671297), B&R International Joint Laboratory of Eurasian Anthropology (18490750300). J.K. was funded by DFG grant KR 4015/1-1, the Baden Württemberg Foundation, and the Max Planck Institute. Accelerator Mass Spectrometry radiocarbon dating work was supported by the National Science Foundation (BCS-1460369) to D.J.K. and B.J.C). D.R. was funded by NSF HOMINID grant BCS-1032255, NIH (NIGMS) grant GM100233, the Paul Allen Foundation, the John Templeton Foundation grant 61220, and the Howard Hughes Medical Institute.

## Author Contributions

Conceptualization, C.-C.W., H.-Y.Y., A.N.P., H.M., A.M.K., L.J., H.L., J.K., R.P., and D.R.; Formal Analysis, C.-C.W., R.B., M.Ma., S.M., Z.Z., B.J.C, and D.R.; Investigation, C.-C.W., K.Si., O.C., A.K., N.R., A.M.K., M.Ma., S.M., K.W., N.A., N.B., K.C., B.J.C, L.E., A.M.L., M.Mi., J.O., K.S., S.W., S.Y., F.Z., J.G., Q.D., L.K., Da.L, Do.L, R.L., W.C., R.S., L.-X.W., L.W., G.X., H.Y., M.Z., G.H., X.Y., R.H., S.S., D.J.K., L.J., H.L., J.K., R.P., and D.R.; Resources, H.-Y.Y., A.N.P., R.B., D.T., J.Z., Y.-C.L, J.-Y.L., M.Ma., S.M., Z.Z., R.C., C.-J. H., C.-C.S., Y.G.N., A.V.T., A.A.T., S.L., Z.-Y.S., X.-M.W., T.-L.Y., X.H., L.C., H.D., J.B., E.Mi., D.E., T.-O.I., E.My., H.K.-K., M.N., K.Sh., D.J.K., R.P., and D.R.; Data Curation, C.-C.W., K.Si., O.C., A.K., N.R., R.B., M.Ma., S.M., B.J.C, L.E., A.A.T., and D.R.; Writing, C.-C.W., H.-Y.Y., A.N.P., H.M., A.K., and D.R.; Supervision, C.-C.W., H.-Q.Z., N.R., M.R., S.S., D.J.K., L.J., H.L., J.K., R.P., and D.R.

