## Supplementary Tables and Figures for "The Genomic Formation of Human Populations in East Asia"

**Table S1. Population information for newly genotyped individuals.**

| **Population** | **Language** | **Location** | **Latitude** | **Longitude** | **N** |
| --- | --- | --- | --- | --- | --- |
| Tibetan | Tibetic, Sino-Tibetan | Chamdo, Tibet, China | 31.1 | 97.2 | 12 |
| Tibetan | Tibetic, Sino-Tibetan | Gangcha, Qinghai, China | 37.3 | 100.2 | 20 |
| Tibetan | Tibetic, Sino-Tibetan | Gannan, Gansu, China | 35.0 | 102.9 | 5 |
| Tibetan | Tibetic, Sino-Tibetan | Lhasa, Tibet, China | 30.0 | 91.1 | 9 |
| Tibetan | Tibetic, Sino-Tibetan | Nagqu, Tibet, China | 31.5 | 92.1 | 7 |
| Tibetan | Tibetic, Sino-Tibetan | Shannan, Tibet, China | 29.2 | 91.8 | 10 |
| Tibetan | Tibetic, Sino-Tibetan | Shigatse, Tibet, China | 29.3 | 88.9 | 10 |
| Tibetan | Tibetic, Sino-Tibetan | Xinlong, Sichuan, China | 31.0 | 100.3 | 10 |
| Tibetan | Tibetic, Sino-Tibetan | Xunhua, Qinghai, China | 35.8 | 102.5 | 4 |
| Tibetan | Tibetic, Sino-Tibetan | Yajiang, Sichuan, China | 30.0 | 101.0 | 10 |
| Tibetan | Tibetic, Sino-Tibetan | Yunnan, China | 27.8 | 99.7 | 4 |
| Qiang | Qiangic, Sino-Tibetan | Daofu, Sichuan, China | 30.9 | 101.1 | 11 |
| Qiang | Qiangic, Sino-Tibetan | Danba, Sichuan, China | 30.8 | 101.9 | 9 |
| Han | Chinese, Sino-Tibetan | Chongqing, China | 29.3 | 106.3 | 3 |
| Han | Chinese, Sino-Tibetan | Fujian, China | 26.1 | 119.3 | 5 |
| Han | Chinese, Sino-Tibetan | Guangdong, China | 23.2 | 113.2 | 7 |
| Han | Chinese, Sino-Tibetan | Henan, China | 34.8 | 113.6 | 5 |
| Han | Chinese, Sino-Tibetan | Hubei, China | 30.5 | 114.3 | 5 |
| Han | Chinese, Sino-Tibetan | Jiangsu, China | 32.1 | 118.8 | 7 |
| Han | Chinese, Sino-Tibetan | Shandong, China | 36.6 | 117.0 | 10 |
| Han | Chinese, Sino-Tibetan | Shanghai, China | 31.2 | 121.5 | 2 |
| Han | Chinese, Sino-Tibetan | Shanxi, China | 37.9 | 112.5 | 8 |
| Han | Chinese, Sino-Tibetan | Sichuan, China | 30.7 | 104.1 | 7 |
| Han | Chinese, Sino-Tibetan | Zhejiang, China | 30.3 | 120.2 | 5 |
| Zhuang | Tai, Tai-Kadai | Guangxi, China | 22.8 | 108.4 | 22 |
| Li | Hlai, Tai–Kadai | Hainan, China | 18.5 | 110 | 4 |
| Dong | Kam-Sui, Tai–Kadai | Guizhou, China | 26.7 | 106.6 | 13 |
| Dong | Kam-Sui, Tai–Kadai | Hunan, China | 27.4 | 109.2 | 7 |
| Mulam | Kam-Sui, Tai–Kadai | Luocheng, Guangxi, China | 24.8 | 108.9 | 17 |
| Maonan | Kam-Sui, Tai–Kadai | Huanjiang, Guangxi, China | 24.8 | 108.3 | 17 |
| Gelao | Kra, Tai-Kadai | Longlin, Baise, Guangxi, China | 24.8 | 105.3 | 10 |
| Bonan | Mongolic | Jishishan, Gansu, China | 35.7 | 102.8 | 10 |
| Dongxiang | Mongolic | Linxia, Gansu, China | 35.6 | 103.2 | 7 |
| Yugur-Eastern | Mongolic | Sunan, Gansu, China | 38.9 | 99.6 | 16 |
| Kazakh | Kipchak, Turkic | Kazak Autonomous County of Aksay, Gansu, China | 38.5 | 94.3 | 8 |
| Kyrgyz | Kipchak, Turkic | Urumqi, Xinjiang, China | 43.8 | 87.7 | 13 |
| Yugur-Western | Turkic | Sunan, Gansu, China | 38.9 | 99.6 | 1 |
| Salar | Oghuz, Turkic | Xunhua, Qinghai, China | 35.8 | 102.5 | 8 |
| Bahun | Nepali, Indo-European | Nepal | 27.4 | 85.3 | 5 |
| Gurung | Tamangic, Sino-Tibetan | Nepal | 27.4 | 86.2 | 5 |
| Magar | Magaric, Sino-Tibetan | Nepal | 27.4 | 86.2 | 6 |
| Newar | Sino-Tibetan | Nepal | 27.4 | 85.3 | 8 |
| Rai | Kiranti/Nepali | Nepal | 27.4 | 85.3 | 5 |
| Sherpa | Tibetic, Bodish, Sino-Tibetan | Nepal | 27.4 | 85.3 | 4 |
| Tamang | Tamangic, Sino-Tibetan | Nepal | 27.4 | 86.2 | 8 |
| Tharu | Indo-Aryan, Indo-European | Nepal | 27.4 | 86.2 | 5 |

**Table S2. Kinship detected between pairs of individuals.**

| **Country** | **Site** | **Family ID** | **N** | **Individuals** | **Relationship** | **Date** |
| --- | --- | --- | --- | --- | --- | --- |
| Japan | Rokutsu | Rokutsu.Family | 2 | I13886-I13887 | Brothers | 1500-1000 BCE |
| China | Wuzhuangguoliang | Wuzh.Family1 | 3 | 18R21264-S120-18R21267 | 18R21264 is a 1^st^ or 2^nd^ degree relative of the other two individuals | 150-465 CE |
| China | Wuzhuangguoliang | Wuzh.Family2 | 2 | AH17-18R21268 | 1^st^ degree relatives | 150-465 CE |
| China | Wuzhuangguoliang | Wuzh.Family3 | 2 | S95-S97 | 1^st^ degree relatives | 150-465 CE |
| Taiwan | Hanben | Hanben.Family1 | 2 | I3611-I3612 | 2nd or 3rd degree relatives | 150-465 CE |
| Taiwan | Hanben | Hanben.Family2 | 2 | I15156-I8072 | 1st degree relatives | 510-870 CE |
| Taiwan | Hanben | Hanben.Family3 | 2 | I3734-I3735 | 2nd or 3rd degree relatives | 510-870 CE |
| Taiwan | Hanben | Hanben.Family4 | 3 | I8078-I3735-I3734 | I8078-I1375 1st degree relatives;  I3734 is a 2-3rd relative of I8078 | 510-870 CE |
| Russia | Boisman-2 | Boisman.Family1 | 6 | I3556-I14819-I14771-I14772-I14773-I14774 | father-mother-son-daughter-son2-daughter2 | 3696-3638 calBCE [based on I3556] |
| Russia | Boisman-2 | Boisman.Family2 | 2 | I1206-I1192 | 1st degree relatives | 4932-4834 calBCE [intersection] |
| Russia | Boisman-2 | Boisman.Family3 | 2 | I14307-I14308 | 1st degree relatives | 5400-3600 BCE |
| Mongolia | Marzyn | Marzyn.Family | 3 | I11696-I11697-I11698 | 2nd or 3rd degree relatives | 5617-5531 calBCE [intersection] |
| Mongolia | Ulaangom | Ulaangom.Family1 | 2 | I7029-I6230 | father-son | 354-114 calBCE [based on I7029] |
| Mongolia | Ulaangom | Ulaangom.Family2 | 2 | I6231-I6232 | 2nd or 3rd degree relatives | 387-209 calBCE [based on I6232] |
| Mongolia | Ulaangom | Ulaangom.Family3 | 2 | I12970-I7028 | 1st or 2nd degree relatives | 381-235 calBCE [intersection] |
| Mongolia | Ulaangom | Ulaangom.Family4 | 2 | I6224-I6225 | siblings | 361-203 calBCE [based on I6224] |

Note: We also detect a family of two individuals (I12506-I12955) as 1st or 2nd degree relatives. However, I12506 has low quality data (0.004x average coverage) and does not cluster with other individuals from the same archaeological culture assignment, while individual I12955 has high quality data (4.69x average coverage) and does cluster with other individuals from the same archaeological culture assignment. The direct radiocarbon dates on the two individuals are separated in time by more than 300 years, and in addition, the two sites are separated by 15 degrees of longitude. We therefore assume the detected relatedness is a false-positive and do not declare this to be a family.

**Table S3. Admixture time with West Eurasian-related groups and lower bound of proportion estimated by ALDER. Populations (restricted to those with ≥4 individuals) are sorted by 1-ref Z-score for French.**

| **Population** | **N** | **2-ref decay for Li and French (generations)** | **2-ref z-score** | **1-ref decay for**  **French (gens)** | **1-ref Z score** | **Mixture fraction % lower bound** |
| --- | --- | --- | --- | --- | --- | --- |
| **Tibetan_Gangcha** | 20 | 26.84 ± 3.64 | 7.37 | 28.66 ± 2.68 | 10.68 | 5.0 ± 0.4 |
| **Kyrgyz_China** | 13 | 19.36 ± 2.17 | 8.93 | 22.87 ± 2.15 | 10.64 | 23.3 ± 1.1 |
| **Dongxiang** | 7 | 31.78 ± 5.20 | 6.11 | 24.94 ± 2.72 | 9.19 | 11.7 ± 1.0 |
| **Salar** | 8 | 22.79 ± 3.41 | 6.69 | 23.74 ± 2.73 | 8.7 | 12.5 ± 0.8 |
| **Tu** | 10 | 28.94 ± 3.14 | 9.22 | 31.38 ± 4.17 | 7.52 | 6.3 ± 0.5 |
| **Kazakh_China** | 8 | 23.10 ± 3.07 | 7.52 | 26.55 ± 4.07 | 6.53 | 20.6 ± 1.3 |
| **Uygur** | 10 | 15.52 ± 1.77 | 8.76 | 17.50 ± 2.71 | 6.47 | 28.6 ± 1.9 |
| **Bonan** | 10 | 24.12 ± 3.67 | 6.57 | 23.39 ± 3.75 | 6.23 | 7.8 ± 0.9 |
| **Yugur** | 16 | 21.63 ± 4.75 | 4.56 | 28.49 ± 4.80 | 5.93 | 5.1 ± 0.5 |
| **Even** | 9 | 6.50 ± 1.58 | 4.12 | 8.28 ± 1.40 | 5.92 | 19.8 ± 1.0 |
| **CHB** | 103 | 43.26 ± 17.53 | 2.47 | 39.60 ± 6.92 | 5.72 | 1.2 ± 0.2 |
| **Tibetan_Shigatse** | 10 | 76.73 ± 117.93 | 0.63 | 26.18 ± 4.64 | 5.64 | 2.0 ± 0.3 |
| **Evenk_FarEast** | 5 | 4.17 ± 0.73 | 5.73 | 4.87 ± 0.89 | 5.49 | 13.0 ± 1.3 |
| **Oroqen** | 9 | 34.98 ± 6.49 | 5.39 | 30.37 ± 5.54 | 5.48 | 3.6 ± 0.6 |
| **KHV** | 91 | 31.42 ± 16.09 | 1.95 | 29.40 ± 5.74 | 5.12 | 1.2 ± 0.1 |
| **Xibo** | 7 | 14.12 ± 3.63 | 3.89 | 14.56 ± 3.12 | 4.66 | 3.5 ± 0.5 |
| **Cambodian** | 8 | 20.73 ± 6.80 | 3.05 | 50.36 ± 11.49 | 4.38 | 4.9 ± 0.8 |
| **Ulchi** | 25 | 8.31 ± 3.83 | 2.17 | 27.86 ± 5.01 | 4.37 | 1.6 ± 0.3 |
| **Newar** | 7 | 39.41 ± 6.28 | 6.27 | 35.55 ± 9.58 | 3.71 | 14.8 ± 2.1 |
| **Lahu** | 8 | 149.71 ± 149.73 | 0.9 | 10.59 ± 2.87 | 3.69 | 1.9 ± 0.4 |
| **Tibetan_Nagqu** | 8 | 27.42 ± 15.34 | 1.79 | 33.06 ± 9.17 | 3.61 | 2.2 ± 0.5 |
| **Evenk_Transbaikal** | 8 | 6.50 ± 1.98 | 3.28 | 6.31 ± 1.76 | 3.58 | 5.1 ± 0.6 |
| **Tibetan_Lhasa** | 9 | 68.85 ± 17.84 | 3.61 | 47.71 ± 14.20 | 3.36 | 3.4 ± 0.8 |
| **Nanai** | 10 | 27.71 ± 11.19 | 2.48 | 31.75 ± 9.04 | 3.26 | 1.8 ± 0.5 |
| **Tibetan_Yajiang** | 10 | 15.84 ± 4.51 | 3.51 | 35.33 ± 11.33 | 3.12 | 2.0 ± 0.4 |
| **Tharu** | 5 | 40.44 ± 22.44 | 1.8 | 9.94 ± 3.31 | 3.00 | 8.6 ± 1.3 |
| **Nivh** | 10 | 500.00 ± inf | 0 | 36.15 ± 12.15 | 2.97 | 3.1 ± 0.8 |
| **Daur** | 9 | 20.51 ± 6.24 | 3.29 | 18.17 ± 6.21 | 2.93 | 3.4 ± 0.7 |
| **Thai** | 10 | 26.01 ± 9.41 | 2.76 | 36.36 ± 12.49 | 2.91 | 6.4 ± 0.8 |
| **Han_NChina** | 10 | 32.78 ± 12.79 | 2.56 | 45.45 ± 16.05 | 2.83 | 2.5 ± 0.8 |
| **Rai** | 5 | 4.49 ± 2.55 | 1.76 | 14.38 ± 5.52 | 2.61 | 2.5 ± 0.6 |
| **Tibetan_Xinlong** | 10 | 17.52 ± 6.99 | 2.48 | 24.67 ± 9.54 | 2.59 | 1.1 ± 0.4 |
| **Han_Henan** | 5 | 11.02 ± 7.21 | 1.53 | 58.22 ± 20.36 | 2.58 | 3.0 ± 1.1 |
| **Tamang** | 7 | 27.15 ± 10.41 | 2.61 | 17.55 ± 6.87 | 2.55 | 4.6 ± 0.9 |
| **Kusunda** | 10 | 13.23 ± 5.74 | 2.31 | 5.44 ± 2.14 | 2.54 | 4.1 ± 0.6 |
| **Tibetan_Gannan** | 5 | 19.97 ± 7.48 | 2.67 | 20.53 ± 8.40 | 2.45 | 7.9 ± 1.3 |
| **Miao** | 10 | 367.64 ± inf | 0 | 110.12 ± 45.69 | 2.32 | 2.0 ± 0.9 |
| **Gurung** | 5 | 48.55 ± 26.51 | 1.83 | 10.66 ± 4.71 | 2.26 | 2.2 ± 0.7 |
| **Sherpa** | 4 | 110.97 ± inf | 0 | 54.38 ±24.38 | 2.23 | 4.6 ± 1.8 |
| **Tibetan_Shannan** | 9 | 76.89 ± 68.37 | 1.12 | 70.86 ± 31.86 | 2.22 | 3.7 ± 1.6 |
| **CHS** | 105 | 324.08 ± 159.01 | 0.51 | 173.98 ± 56.15 | 2.18 | 1.1 ± 0.5 |
| **Han_Shanxi** | 8 | 32.41 ± 7.85 | 4.13 | 35.04 ± 17.52 | 2.00 | 1.7 ± 0.4 |
| **Han_Shandong** | 10 | 59.13 ± 27.02 | 1.93 | 82.53 ± 42.66 | 1.93 | 1.7 ± 0.7 |
| **Hezhen** | 8 | 15.36 ± 7.03 | 2.19 | 42.94 ± 22.96 | 1.87 | 3.9 ± 1.1 |
| **Mulam** | 17 | 52.74 ± 30.42 | 1.13 | 5.80 ± 3.10 | 1.79 | 0.1 ± 0.1 |
| **Gelao** | 10 | 2.00 ± inf | 0 | 2.79 ± 1.17 | 1.72 | 0.3 ± 0.2 |
| **Magar** | 6 | 8.64 ± 31.02 | 0.28 | 11.27 ± 6.62 | 1.70 | 4.9 ± 1.1 |
| **Mongola** | 6 | 25.27 ± 10.53 | 2.4 | 22.56 ± 13.97 | 1.62 | 4.4 ± 1.1 |
| **Dong_Guizhou** | 13 | 147.00 ± 463.38 | 0.14 | 174.70 ± 57.27 | 1.60 | 2.1 ± 1.3 |
| **Tibetan_Chamdo** | 12 | 48.36 ± 22.90 | 2.11 | 94.35 ± 61.85 | 1.53 | 2.7 ± 1.8 |
| **She** | 10 | 164.17 ± 186.17 | 0.52 | 15.30 ± 7.87 | 1.46 | 0.3 ± 0.2 |
| **Han_Guangdong** | 7 | 270.31 ± inf | 0 | 241.96 ± 134.25 | 1.08 | 6.0 ± 5.1 |
| **Han_Jiangsu** | 7 | 33.92 ± 65.64 | 0.52 | 61.55 ± 40.34 | 1.08 | 0.6 ± 0.6 |
| **Dai** | 10 | 20.09 ± 28.38 | 0.71 | 168.40 ± 113.10 | 1.05 | 2.7 ± 2.5 |
| **Vietnamese** | 10 | 61.77 ± 45.03 | 1.37 | 38.57 ± 37.43 | 1.03 | 0.8 ± 0.6 |
| **Dong_Hunan** | 7 | 22.94 ± 5.69 | 3.14 | 42.68 ± 26.78 | 1.01 | 0.5 ± 0.5 |
| **CDX** | 93 | 42.53 ± 28.85 | 1.42 | 39.20 ± 51.81 | 0.76 | 0.5 ± 0.4 |
| **Korean** | 6 | 73.51 ± 79.60 | 0.8 | 96.59 ± 121.94 | 0.76 | 1.4 ± 1.9 |
| **Han_HGDP** | 33 | 20.08 ± 5.44 | 2.91 | 7.05 ± 10.01 | 0.70 | 0.2 ± 0.1 |
| **Naxi** | 9 | 40.09 ± 60.84 | 0.66 | 150.52 ± 170.60 | 0.68 | 2.0 ± 2.8 |
| **Qiang_Daofu** | 11 | 17.16 ± 9.27 | 1.85 | 15.63 ± 23.61 | 0.66 | 0.6 ± 0.4 |
| **Kinh** | 8 | 24.08 ± 12.56 | 1.92 | 193.23 ± 191.37 | 0.64 | 7.6 ± 10.4 |
| **Qiang_Danba** | 9 | 32.79 ± inf | 0 | 127.69 ± 218.08 | 0.40 | 2.7 ± 6.1 |
| **Atayal** | 9 | 3.50 ± inf | 0 | 17.56 ± 6.84 | 0.27 | 1.2 ± 2.1 |
| **Tujia** | 10 | 163.12 ± 202.59 | 0.54 | 34.50 ± 204.42 | 0.17 | 0.2 ± 0.8 |
| **Yi** | 10 | 26.37 ± 10.44 | 2.13 | 16.04 ± 138.25 | 0.12 | 0.4 ± 1.9 |
| **Ami** | 10 | 197.52 ± inf | 0 | 22.94 ± inf | 0 | 0.3 ± 95.2 |
| **Bahun** | 4 | 14.74 ± 8.90 | 1.66 | 2.00 ± inf | 0 | 5.0 ± 0.0 |
| **Han_Fujian** | 5 | 10.77 ± inf | 0 | 3.65 ± inf | 0 | 0.0 ± 95.4 |
| **Han_Hubei** | 5 | 17.21 ± 9.14 | 1.42 | 19.10 ± inf | 0 | 0.0 ± 154.9 |
| **Han_Sichuan** | 7 | 2.00 ± inf | 0 | 74.54 ± inf | 0 | 1.0 ± 94.5 |
| **Japanese** | 29 | 38.12 ± inf | 0 | 73.78 ± inf | 0 | 0.5 ± 95.0 |
| **JPT** | 104 | 500.00 ± inf | 0 | 454.75 ± inf | 0 | 4.5 ± 205.4 |
| **Maonan** | 17 | 14.05 ± inf | 0 | 2.00 ± inf | 0 | 0.0 ± 225.1 |
| **Tibetan_Yunnan** | 4 | 2.00 ± inf | 0 | 2.00 ± inf | 0 | 2.1 ± 155.0 |
| **Zhuang** | 22 | 470.02 ± inf | 0 | 500.00 ± inf | 0 | 8.8 ± 203.5 |
| **Han_Zhejiang** | 5 | 5.30 ± 6.08 | 0.87 | 113.59 ± 40.89 | -2.12 | 0.0 ± 0.0 |

**Table S4. West Eurasian-related admixture proportions estimated using *qpAdm*. Here “p” refers to the P-value for rank=1 and “std.err” is the standard error estimated using a block jackknife. We used *qpAdm* to estimate West Eurasia-related ancestry in present-day East Asians using Mbuti.DG, Ust_Ishim.DG, Russia_Kostenki14, Papuan.DG, Australian.DG, and Onge.DG as outgroups and Li and French as proxies for the source populations.**

|  | **p** | **Li** | **French** | **std.err** |
| --- | --- | --- | --- | --- |
| Bahun | 0.138 | 0.498 | 0.502 | 0.019 |
| Uyghur | 0.540 | 0.536 | 0.464 | 0.013 |
| Kyrgyz_China | 0.275 | 0.648 | 0.352 | 0.014 |
| Even | 0.027 | 0.649 | 0.351 | 0.016 |
| Kazakh_China | 0.103 | 0.683 | 0.317 | 0.016 |
| Newar | 0.288 | 0.740 | 0.260 | 0.017 |
| Magar | 0.091 | 0.787 | 0.213 | 0.016 |
| Tharu | 0.282 | 0.794 | 0.206 | 0.019 |
| Evenk_FarEast | 0.431 | 0.830 | 0.170 | 0.019 |
| Salar | 0.581 | 0.837 | 0.163 | 0.017 |
| Evenk_Transbaikal | 0.354 | 0.864 | 0.136 | 0.019 |
| Dongxiang | 0.298 | 0.866 | 0.134 | 0.017 |
| Kusunda | 0.856 | 0.871 | 0.129 | 0.019 |
| Bonan | 0.719 | 0.897 | 0.103 | 0.017 |
| Tibetan_Gannan | 0.010 | 0.897 | 0.103 | 0.019 |
| Tu | 0.736 | 0.913 | 0.087 | 0.016 |
| Mongola | 0.229 | 0.916 | 0.084 | 0.017 |
| Tibetan_Gangcha | 0.478 | 0.922 | 0.078 | 0.017 |
| Xibo | 0.436 | 0.929 | 0.071 | 0.017 |
| Oroqen | 0.229 | 0.931 | 0.069 | 0.017 |
| Tamang | 0.694 | 0.934 | 0.066 | 0.019 |
| Tibetan_Xunhua | 0.571 | 0.936 | 0.064 | 0.019 |
| Yugur | 0.246 | 0.941 | 0.059 | 0.016 |
| Daur | 0.950 | 0.945 | 0.055 | 0.018 |
| Ulchi | 0.655 | 0.952 | 0.048 | 0.017 |
| Hezhen | 0.811 | 0.955 | 0.045 | 0.018 |
| Rai | 0.876 | 0.957 | 0.043 | 0.020 |
| Nanai | 0.271 | 0.957 | 0.043 | 0.019 |
| Han_Shanghai | 0.033 | 0.960 | 0.040 | 0.023 |
| Thai | 0.839 | 0.960 | 0.040 | 0.017 |
| Tibetan_Chamdo | 0.420 | 0.962 | 0.038 | 0.018 |
| Tibetan_Shigatse | 0.270 | 0.965 | 0.035 | 0.019 |
| Gurung | 0.286 | 0.966 | 0.034 | 0.020 |
| Tibetan_Yunnan | 0.040 | 0.967 | 0.033 | 0.022 |
| Han_NChina | 0.737 | 0.968 | 0.032 | 0.016 |
| Tibetan_Yajiang | 0.422 | 0.969 | 0.031 | 0.018 |
| Tibetan_Nagqu | 0.201 | 0.971 | 0.029 | 0.020 |
| Tibetan_Lhasa | 0.175 | 0.971 | 0.029 | 0.019 |
| Negidal | 0.256 | 0.974 | 0.026 | 0.022 |
| Tibetan_Shannan | 0.017 | 0.978 | 0.022 | 0.019 |
| Nivkh | 0.456 | 0.979 | 0.021 | 0.020 |
| Sherpa | 0.392 | 0.982 | 0.018 | 0.022 |
| Cambodian | 0.026 | 0.984 | 0.016 | 0.018 |
| Qiang_Danba | 0.216 | 0.984 | 0.016 | 0.018 |
| Korean | 0.546 | 0.985 | 0.015 | 0.018 |
| Qiang_Daofu | 0.065 | 0.986 | 0.014 | 0.018 |
| Tibetan_Xinlong | 0.362 | 0.987 | 0.013 | 0.017 |
| Han_Guangdong | 0.807 | 0.991 | 0.009 | 0.017 |
| Han_Henan | 0.744 | 0.992 | 0.008 | 0.020 |
| Han_Shanxi | 0.746 | 0.993 | 0.007 | 0.018 |
| Yi | 0.677 | 0.995 | 0.005 | 0.018 |
| Han_Shandong | 0.414 | 0.997 | 0.003 | 0.017 |
| Miao | 0.508 | 1.000 | 0.000 | 0.016 |
| Dong_Hunan | 0.099 | 1.000 | 0.000 | 0.018 |
| Kinh | 0.759 | 1.001 | -0.001 | 0.016 |
| Tujia | 0.492 | 1.001 | -0.001 | 0.017 |
| Han_Jiangsu | 0.914 | 1.001 | -0.001 | 0.018 |
| Japanese | 0.397 | 1.002 | -0.002 | 0.016 |
| CHB.SG | 0.125 | 1.002 | -0.002 | 0.015 |
| Naxi | 0.947 | 1.003 | -0.003 | 0.018 |
| Han_Sichuan | 0.204 | 1.003 | -0.003 | 0.018 |
| Han_HGDP | 0.449 | 1.007 | -0.007 | 0.015 |
| Han_Fujian | 0.807 | 1.009 | -0.009 | 0.019 |
| Mulam | 0.434 | 1.009 | -0.009 | 0.016 |
| She | 0.124 | 1.011 | -0.011 | 0.017 |
| Maonan | 0.214 | 1.012 | -0.012 | 0.015 |
| KHV.SG | 0.182 | 1.012 | -0.012 | 0.014 |
| JPT.SG | 0.051 | 1.013 | -0.013 | 0.016 |
| CHS.SG | 0.146 | 1.013 | -0.013 | 0.015 |
| Dong_Guizhou | 0.882 | 1.014 | -0.014 | 0.016 |
| Han_Zhejiang | 0.088 | 1.014 | -0.014 | 0.019 |
| Dai | 0.269 | 1.015 | -0.015 | 0.016 |
| Han_Chongqing | 0.292 | 1.016 | -0.016 | 0.020 |
| Zhuang | 0.775 | 1.016 | -0.016 | 0.015 |
| Vietnamese | 0.242 | 1.020 | -0.020 | 0.016 |
| Gelao | 0.669 | 1.020 | -0.020 | 0.017 |
| CDX.SG | 0.180 | 1.022 | -0.022 | 0.015 |
| Ami | 0.214 | 1.024 | -0.024 | 0.018 |
| Han_Hubei | 0.440 | 1.030 | -0.030 | 0.018 |
| Lahu | 0.153 | 1.032 | -0.032 | 0.019 |
| Nicobarese | 0.256 | 1.036 | -0.036 | 0.024 |

**Table S5: *f_3_* and *f_4_*-statistics documenting key findings of this study.** **|Z|>3 scores are highlighted.**

| **Populations** | **Genetic continuity from ancient Taiwan to present-day Austronesian speaking populations** | | **Ancient Taiwan and Austronesians have extra affinity with Tai-Kadai populations** | | **Extra affinity between Amur Basin Neolithic and Mongolia Neolithic populations with Native Americans but not with Mal'ta1** | | **Extra affinity between Jomon, ancient Taiwan / Austronesian-speaker and Amur River Basin populations** | |
| --- | --- | --- | --- | --- | --- | --- | --- | --- |
|  | ***f_3_* (Taiwan_Hanben,X; Mbuti)** | ***f_4_* (Taiwan_Hanben/Gongguan, Mbuti; Ami, X)** | ***f_4_* (Taiwan_Hanben, Mbuti; Maonan, X)** | ***f_4_* (Ami, Mbuti; Maonan, X)** | ***f_4_* (Mixe, Mbuti; X, Boisman/Mongolia_Neolithic)** | ***f_4_* (Mal'ta1, Mbuti; X, Boisman/ Mongolia_Neolithic)** | ***f_4_* (Japan_Jomon, Mbuti; X, Taiwan_Hanben)** | ***f_4_* (Japan_Jomon, Mbuti; X, Ami/Atayal)** |
| **Tibetan_Chamdo** | 0.277602 | 37.804/23.464 | 28.264 | 27.238 | -8.55/-5.435 | -0.035/-0.865 | -9.451 | -9.158/-7.408 |
| **CDX.SG** | 0.291633 | 25.312/16.764 | 5.137 | 5.768 | -10.565/-6.272 | -0.589/-1.076 | -6.029 | -5.829/-3.934 |
| **Dai** | 0.291423 | 21.708/13.673 | 4.084 | 3.586 | -9.542/-6.192 | 0.408/-0.636 | -5.46 | -5.288/-3.999 |
| **Maonan** | 0.293168 | 20.622/12.868 | - | - | -8.497/-5.478 | -0.262/-1.102 | -4.901 | -4.79/-3.295 |
| **KHV.SG** | 0.290154 | 27.832/18.581 | 9.887 | 11.594 | -10.501/-6.264 | -0.083/-0.734 | -7.111 | -6.767/-4.595 |
| **Gelao** | 0.290894 | 21.421/14.218 | 4.794 | 4.637 | -8.145/-5.413 | -0.458/-1.075 | -4.953 | -4.904/-3.517 |
| **CHS.SG** | 0.29152 | 25.128/16.168 | 5.187 | 6.875 | -6.949/-3.697 | -0.28/-0.676 | -3.985 | -3.949/-2.579 |
| **Zhuang** | 0.292306 | 22.421/14.027 | 2.44 | 1.424 | -8.679/-5.382 | -0.267/-1.067 | -4.165 | -4.185/-2.776 |
| **Dong_Hunan** | 0.292078 | 19.187/12.155 | 2.222 | 2.680 | -7.202/-4.904 | -0.647/-1.264 | -3.785 | -3.895/-2.815 |
| **Japanese** | 0.28588 | 32.028/20.113 | 17.136 | 15.958 | -4.548/-2.75 | 0.093/-0.963 | 17.144 | 16.137/13.45 |
| **Han_HGDP** | 0.290582 | 24.967/16.287 | 7.376 | 7.231 | -6.681/-4.072 | -0.524/-1.26 | -4.893 | -4.669/-3.311 |
| **Han_Hubei** | 0.29078 | 19.728/11.316 | 4.22 | 3.773 | -4.422/-2.864 | 0.754/-0.375 | -2.828 | -2.949/-2.045 |
| **Mulam** | 0.292378 | 21.526/13.858 | 2.092 | 1.083 | -7.958/-5.304 | -0.637/-1.366 | -5.25 | -5.205/-3.59 |
| **Lahu** | 0.286693 | 26.271/18.113 | 11.956 | 11.893 | -9.813/-6.703 | -0.286/-1.058 | -5.683 | -5.665/-4.317 |
| **CHB.SG** | 0.289108 | 27.692/17.723 | 11.488 | 13.716 | -5.948/-3.034 | 0.061/-0.568 | -4.05 | -4.125/-2.77 |
| **Vietnamese** | 0.290574 | 22.674/14.918 | 6.175 | 6.175 | -9.665/-6.231 | -0.673/-1.33 | -5.325 | -5.226/-3.892 |
| **JPT.SG** | 0.286757 | 31.629/19.186 | 16.318 | 17.928 | -5.007/-1.992 | -0.011/-0.342 | 18.132 | 17.226/14.406 |
| **Han_Chongqing** | 0.29057 | 18.187/10.614 | 3.83 | 3.469 | -4.825/-3.268 | -0.276/-0.884 | -2.934 | -3.01/-2.399 |
| **Ami** | 0.304245 | - | -20.622 | - | -7.846/-5.385 | -0.017/-0.956 | 0.322 | -0.322/0.243 |
| **She** | 0.290866 | 22.04/13.608 | 4.715 | 3.804 | -6.221/-3.979 | -0.62/-1.284 | -4.282 | -4.325/-3.176 |
| **Han_Fujian** | 0.290577 | 20.362/13.181 | 4.52 | 3.746 | -5.957/-3.901 | -0.413/-1.093 | -2.76 | -2.955/-2.131 |
| **Han_Shanghai** | 0.290151 | 15.286/9.708 | 3.772 | 3.432 | -2.879/-2.093 | 0.548/-0.236 | -1.534 | -1.618/-1.209 |
| **Dong_Guizhou** | 0.291619 | 21.968/14.478 | 3.915 | 3.484 | -8.365/-5.352 | -0.794/-1.46 | -4.438 | -4.618/-3.15 |
| **Tujia** | 0.28939 | 23.953/14.97 | 8.356 | 7.182 | -6.505/-4.214 | 0.053/-0.754 | -4.127 | -4.046/-2.885 |
| **Yi** | 0.283827 | 31.685/19.889 | 18.975 | 17.524 | -7.137/-4.823 | 0.372/-0.657 | -6.601 | -6.397/-4.814 |
| **Han_Guangdong** | 0.291912 | 19.738/11.457 | 2.548 | 1.923 | -6.667/-4.359 | -0.464/-1.284 | -3.373 | -3.449/-2.39 |
| **Nicobarese** | 0.279463 | 27.547/18.014 | 17.016 | 16.238 | -15.081/-11.433 | -1.332/-1.927 | -9.495 | -9.543/-8.649 |
| **Atayal** | 0.304265 | -0.031/2.83 | -17.364 | -14.989 | -7.216/-4.946 | -0.415/-1.165 | -0.243 | -0.466/0.466 |
| **Kinh** | 0.289884 | 24.383/15.394 | 7.057 | 7.188 | -9.808/-6.614 | -0.424/-1.151 | -6.059 | -6.1/-4.563 |
| **Naxi** | 0.28238 | 30.983/20.624 | 20.893 | 18.948 | -7.444/-4.976 | 0.215/-0.827 | -6.014 | -5.81/-4.828 |
| **Korean** | 0.286861 | 25.235/16.442 | 11.08 | 10.706 | -4.287/-2.932 | 0.768/-0.57 | -0.872 | -1.078/-0.54 |
| **Cambodian** | 0.279637 | 36.869/25.164 | 25.854 | 24.911 | -15.85/-10.525 | -0.317/-1.181 | -12.819 | -12.121/-9.973 |
| **Miao** | 0.289672 | 23.151/14.769 | 7.338 | 6.867 | -6.748/-4.307 | -0.608/-1.307 | -3.636 | -3.7/-2.647 |
| **Han_Shandong** | 0.286383 | 28.923/17.477 | 13.994 | 13.027 | -5.152/-3.364 | 0.04/-0.875 | -5.168 | -5.161/-3.846 |
| **Han_Shanxi** | 0.285519 | 29.076/18.105 | 14.747 | 14.041 | -4.665/-3.094 | -0.288/-1.172 | -4.893 | -4.769/-3.757 |
| **Han_Sichuan** | 0.289021 | 23.777/14.704 | 8.276 | 7.350 | -5.932/-3.978 | 0.277/-0.68 | -4.24 | -4.283/-3.17 |
| **Qiang_Daofu** | 0.281697 | 33.675/21.063 | 21.933 | 20.934 | -6.983/-4.588 | 0.203/-0.802 | -6.655 | -6.686/-5.145 |
| **Han_Jiangsu** | 0.289173 | 23.35/15.172 | 7.909 | 7.105 | -5.249/-3.366 | -0.506/-1.237 | -2.662 | -2.822/-1.923 |
| **Tibetan_Xinlong** | 0.283367 | 32.819/20.839 | 19.754 | 18.139 | -7/-4.656 | 0.028/-0.964 | -7.151 | -6.94/-5.374 |
| **Han_Zhejiang** | 0.288948 | 22.848/13.492 | 7.395 | 6.390 | -5.519/-3.72 | -0.554/-1.275 | -3.827 | -3.894/-3.083 |
| **Qiang_Danba** | 0.282884 | 32.193/21.4 | 19.549 | 18.485 | -6.517/-4.154 | 0.149/-0.815 | -7.55 | -7.381/-5.936 |
| **Han_Henan** | 0.28684 | 23.362/16.632 | 10.208 | 10.164 | -4.999/-3.293 | 0.664/-0.403 | -3.659 | -3.73/-2.919 |
| **Tibetan_Lhasa** | 0.276771 | 37.9/24.467 | 28.051 | 26.913 | -9.288/-6.078 | 1.213/-0.174 | -9.614 | -9.256/-7.697 |
| **Sherpa** | 0.276231 | 34.318/23.115 | 23.237 | 22.486 | -8.5/-5.996 | 0.338/-0.568 | -7.709 | -7.842/-6.615 |
| **Li** | 0.292836 | 15.477/11.359 | 0.518 | 0.300 | -8.838/-6.117 | -1.678/-2.031 | -3.375 | -3.373/-2.539 |
| **Tibetan_Yunnan** | 0.280409 | 28.636/19.219 | 17.239 | 16.575 | -7.128/-5.033 | 0.665/-0.379 | -6.271 | -6.141/-5.201 |
| **Yankovsky_IA** | 0.287918 | 13.245/9.064 | 5.288 | 8.192 | 0.476/0.218 | -0.49/-1.44 | 1.843 | 1.204/1.004 |
| **Tibetan_Nagqu** | 0.277312 | 36.899/24.004 | 26.881 | 25.600 | -8.238/-5.48 | 0.979/-0.379 | -8.731 | -8.685/-7.197 |
| **Han_NChina** | 0.284791 | 29.501/19.433 | 16.618 | 15.998 | -6.333/-4.112 | 0.134/-0.873 | -5.464 | -5.604/-4.168 |
| **Nivkh** | 0.27926 | 32.165/20.134 | 21.451 | 19.007 | 1.365/1.067 | 2.728/0.788 | 7.96 | 7.593/7.183 |
| **Tibetan_Shigatse** | 0.276058 | 38.422/24.289 | 28.75 | 27.513 | -8.658/-5.657 | 0.716/-0.485 | -9.676 | -9.232/-7.667 |
| **Thai** | 0.27756 | 39.698/26.424 | 28.948 | 28.002 | -15.958/-10.325 | -1.016/-1.651 | -14.156 | -13.28/-10.888 |
| **Nanai** | 0.28015 | 33.331/20.783 | 21.888 | 20.195 | 0.529/0.545 | 2.797/0.828 | -2.737 | -2.943/-2.177 |
| **Tibetan_Shannan** | 0.275693 | 40.538/25.781 | 30.535 | 28.487 | -9.247/-6.147 | 0.553/-0.589 | -9.328 | -9.163/-7.666 |
| **Ulchi** | 0.278795 | 36.553/22.926 | 25.923 | 23.788 | 1.212/0.904 | 3.232/0.848 | 2.079 | 1.764/1.942 |
| **Oroqen** | 0.279729 | 34.161/20.953 | 23.021 | 21.696 | -1.654/-0.906 | 2.854/0.785 | -5.42 | -5.436/-4.29 |
| **Rai** | 0.275709 | 35.599/21.698 | 24.962 | 23.726 | -8.681/-6.221 | 0.836/-0.339 | -9.353 | -9.504/-7.732 |
| **Hezhen** | 0.280583 | 32.785/22.334 | 21.034 | 19.741 | -2.918/-1.812 | 1.772/0.146 | -6.209 | -6.096/-5.046 |
| **Gurung** | 0.272375 | 39.23/25.544 | 30.135 | 27.647 | -11.271/-7.464 | 0.826/-0.37 | -10.725 | -10.309/-8.805 |
| **Negidal** | 0.279915 | 27.705/17.91 | 17.053 | 15.632 | 2.238/1.948 | 3.294/1.515 | -0.582 | -0.765/-0.39 |
| **Tibetan_Yajiang** | 0.278515 | 37.43/23.837 | 26.908 | 25.579 | -8.685/-5.578 | 0.012/-0.876 | -9.74 | -9.37/-7.717 |
| **Boisman** | 0.283179 | 25.137/17.105 | 14.042 | 16.403 | -/-0.496 | -/-1.203 | 5.151 | 4.655/4.528 |
| **Yugur** | 0.281702 | 35.545/23.663 | 23.851 | 22.479 | -7.561/-4.735 | 1.447/-0.039 | -9.195 | -8.689/-6.752 |
| **Xibo** | 0.280948 | 33.207/20.559 | 21.048 | 20.180 | -5.336/-3.418 | 0.542/-0.632 | -7.104 | -6.887/-5.697 |
| **Nepal_ancient** | 0.275416 | 36.693/22.493 | 25.59 | 25.212 | -10.271/-7.092 | 0.445/-0.375 | -10.322 | -10.02/-8.63 |
| **Daur** | 0.279991 | 35.672/22.791 | 24.1 | 22.361 | -4.564/-2.864 | 1.193/-0.231 | -6.746 | -6.752/-5.4 |
| **Mongola** | 0.281067 | 32.488/19.973 | 19.893 | 18.976 | -5.318/-3.296 | 0.725/-0.448 | -6.975 | -7.242/-5.773 |
| **Russia_SakhalinAinu** | 0.27611 | 14.776/7.32 | 9.312 | 10.389 | -5.412/-4.077 | -0.865/0.27 | 24.473 | 23.642/23.354 |
| **Magar** | 0.255378 | 64.706/44.099 | 61.865 | 31.996 | -12.367/-14.929 | 1.313/0.924 | -28.349 | -25.919/-22.11 |
| **Tibetan_Xunhua** | 0.276823 | 33.745/22.022 | 24.009 | 17.670 | -6.977/-5.836 | 0.485/-0.592 | -9.822 | -9.317/-8.218 |
| **Tibetan_Gangcha** | 0.275853 | 42.705/27.537 | 34.955 | 32.611 | -9.432/-6.042 | 1.482/0.023 | -12.158 | -11.772/-9.289 |
| **Tamang** | 0.268011 | 46.684/31.777 | 38.694 | 36.307 | -14.108/-9.84 | 1.365/0.01 | -15.468 | -14.936/-12.648 |
| **Tibetan_Gannan** | 0.278438 | 33.035/22.438 | 23.096 | 22.306 | -8.501/-5.708 | 1.086/-0.186 | -10.583 | -10.217/-8.544 |
| **Evenk_Transbaikal** | 0.272125 | 39.466/23.984 | 31.106 | 28.174 | -0.168/0.166 | 6.03/2.97 | -7.887 | -7.994/-6.9 |
| **Tu** | 0.27837 | 37.476/25.201 | 27.97 | 25.288 | -8.659/-5.802 | 0.75/-0.552 | -11.292 | -10.716/-8.853 |
| **Heishui_Mohe** | 0.279504 | 18.185/11.605 | 10.749 | 11.110 | -6.184/-5.489 | -0.494/-1.438 | -2.869 | -3.485/-3.204 |
| **Bonan** | 0.278244 | 38.93/25.889 | 28.193 | 27.566 | -9.547/-6.054 | 1.678/0.231 | -11.846 | -11.093/-8.952 |
| **Dongxiang** | 0.275011 | 40.267/25.563 | 30.412 | 28.888 | -9.57/-6.375 | 2.173/0.612 | -12.291 | -12.093/-9.896 |
| **Kusunda** | 0.260627 | 50.915/35.272 | 45.391 | 42.245 | -18.467/-12.384 | 0.892/-0.25 | -19.138 | -17.827/-16.137 |
| **Evenk_FarEast** | 0.269116 | 42.694/28.912 | 33.305 | 31.982 | -5.421/-3.485 | 6.594/3.578 | -10.813 | -10.557/-9.026 |
| **Salar** | 0.272736 | 41.95/27.256 | 34.81 | 32.002 | -12.168/-8.087 | 1.306/-0.124 | -14.902 | -14.116/-11.873 |
| **Tharu** | 0.253894 | 58.001/40.022 | 50.384 | 46.783 | -20.5/-14.527 | 2.538/0.892 | -23.749 | -22.919/-20.015 |
| **Newar** | 0.252509 | 60.607/41.066 | 54.161 | 49.481 | -21.542/-15.485 | 3.839/1.83 | -24.461 | -23.191/-20.422 |
| **Even** | 0.256735 | 57.877/37.713 | 52.111 | 48.297 | -11.887/-8.21 | 8.707/5.026 | -20.32 | -20.041/-17.371 |
| **Kazakh_China** | 0.260765 | 53.971/36.489 | 47.476 | 44.619 | -13.529/-9.31 | 5.398/2.769 | -21.798 | -20.841/-18.288 |
| **Kyrgyz_China** | 0.25848 | 57.641/39.062 | 53.097 | 51.152 | -15.261/-10.306 | 6.435/3.33 | -24.076 | -23.023/-19.896 |
| **Bahun** | 0.23523 | 66.89/46.388 | 62.95 | 59.084 | -26.486/-19.505 | 5.757/3.37 | -30.233 | -28.217/-26.9 |
| **Uygur** | 0.249183 | 65.487/44.14 | 60.933 | 57.834 | -21.577/-14.82 | 6.739/3.721 | -30.177 | -28.882/-25.299 |
| **DevilsCave_N.SG** | 0.279428 | 23.893/14.509 | 14.097 | 6.891 | -0.636/-0.426 | -1.071/-2.299 | -1.11 | -1.258/-0.868 |
| **Japan_Jomon** | 0.270377 | 29.588/18.629 | 20.36 | 18.759 | -11.523/-10.579 | 0.868/-0.773 | - | - |
| **Wuzhuangguoliang** | 0.296561 | 8.928/6.112 | 4.213 | 5.846 | -1.764/0.097 | 0.108/0.624 | -1.176 | -1.402/-0.67 |
| **Lapita** | 0.301996 | 2.939/2.816 | -6.062 | -0.779 | -9.631/-6.946 | -2.456/-2.825 | 0.249 | -0.209/-0.176 |
| **Mongolia_N_East** | 0.278015 | 20.385/12.333 | 12.598 | 13.440 | 0.496 | 1.203 | -2.717 | -3.212/-3.017 |

**Table S6. Statistics of the form of *f_3_* (Mongolia ancient groups; Mongolia_N, X), where X represents worldwide populations. Here we only list the top 5 negative values for each Mongolia ancient populations. We restrict analysis to cases with >50000 overlapping SNPs.**

|  | **Source 1** | **Source 2** | **Target** | ***f_3_*** | **std.err** | **Z** | **SNPs** |
| --- | --- | --- | --- | --- | --- | --- | --- |
| **top 5 negative Z-score values** | Russia_Sintashta_MLBA | Mongolia_N_East | Mongolia_LBA_3_MongunTaiga | -0.026 | 0.001046 | -24.858 | 596995 |
|  | Russia_Srubnaya | Mongolia_N_East | Mongolia_LBA_3_MongunTaiga | -0.0257 | 0.001058 | -24.283 | 589835 |
|  | Russia_Afanasievo | Mongolia_N_East | Mongolia_LBA_3_MongunTaiga | -0.02488 | 0.001047 | -23.755 | 593793 |
|  | England_MBA | Mongolia_N_East | Mongolia_LBA_3_MongunTaiga | -0.02468 | 0.001047 | -23.57 | 593666 |
|  | Russia_Krasnoyarsk_MLBA | Mongolia_N_East | Mongolia_LBA_3_MongunTaiga | -0.0246 | 0.001044 | -23.566 | 572493 |
|  | Russia_Poltavka_lc | Mongolia_N_East | Mongolia_EIA_1_SlabGrave | -0.00767 | 0.00116 | -6.611 | 505347 |
|  | Kazakhstan_Botai.SG | Mongolia_N_East | Mongolia_EIA_1_SlabGrave | -0.00574 | 0.000975 | -5.891 | 619312 |
|  | WSHG | Mongolia_N_East | Mongolia_EIA_1_SlabGrave | -0.00627 | 0.001093 | -5.735 | 516494 |
|  | Kazakhstan_Kumsay_EBA | Mongolia_N_East | Mongolia_EIA_1_SlabGrave | -0.00545 | 0.000981 | -5.551 | 577357 |
|  | Russia_Yamnaya_Kalmykia.SG | Mongolia_N_East | Mongolia_EIA_1_SlabGrave | -0.00502 | 0.000912 | -5.508 | 616480 |
|  | Sweden_BA.SG | Mongolia_N_East | Mongolia_LBA_2_Ulaanzukh | -0.00387 | 0.002334 | -1.657 | 81159 |
|  | Russia_Poltavka_lc | Mongolia_N_East | Mongolia_LBA_2_Ulaanzukh | -0.00238 | 0.001531 | -1.557 | 392759 |
|  | Kazakhstan_Zevakinskiy_MLBA | Mongolia_N_East | Mongolia_LBA_2_Ulaanzukh | -0.00217 | 0.001633 | -1.33 | 336342 |
|  | WSHG | Mongolia_N_East | Mongolia_LBA_2_Ulaanzukh | -0.0018 | 0.001475 | -1.222 | 411688 |
|  | Latvia_LN_CW | Mongolia_N_East | Mongolia_LBA_2_Ulaanzukh | -0.0023 | 0.002047 | -1.123 | 135918 |
|  | Russia_Yamnaya_Samara | Mongolia_N_East | Mongolia_LBA_6_Khovsgol | -0.015 | 0.000674 | -22.245 | 733047 |
|  | Russia_Afanasievo | Mongolia_N_East | Mongolia_LBA_6_Khovsgol | -0.01548 | 0.000708 | -21.851 | 713662 |
|  | Kazakhstan_Botai.SG | Mongolia_N_East | Mongolia_LBA_6_Khovsgol | -0.01812 | 0.000841 | -21.536 | 641092 |
|  | Russia_Yamnaya_Kalmykia.SG | Mongolia_N_East | Mongolia_LBA_6_Khovsgol | -0.0166 | 0.000809 | -20.503 | 635516 |
|  | Russia_Srubnaya | Mongolia_N_East | Mongolia_LBA_6_Khovsgol | -0.01417 | 0.000703 | -20.154 | 714587 |
|  | WSHG | Mongolia_N_East | Mongolia_EBA_1_Ulgii | -0.01806 | 0.001563 | -11.552 | 416120 |
|  | Kazakhstan_Kumsay_EBA | Mongolia_N_East | Mongolia_EBA_1_Ulgii | -0.01669 | 0.001478 | -11.288 | 468669 |
|  | Kazakhstan_Botai.SG | Mongolia_N_East | Mongolia_EBA_1_Ulgii | -0.01535 | 0.001375 | -11.168 | 482894 |
|  | Russia_Afanasievo | Mongolia_N_East | Mongolia_EBA_1_Ulgii | -0.01479 | 0.001344 | -11.004 | 599413 |
|  | Russia_Yamnaya_Kalmykia.SG | Mongolia_N_East | Mongolia_EBA_1_Ulgii | -0.01502 | 0.001392 | -10.783 | 489777 |
|  | Russia_Sintashta_MLBA | Mongolia_N_East | Mongolia_EIA_4_Sagly | -0.02368 | 0.000535 | -44.286 | 725404 |
|  | Russia_Srubnaya | Mongolia_N_East | Mongolia_EIA_4_Sagly | -0.02351 | 0.000547 | -42.957 | 721237 |
|  | Russia_Yamnaya_Samara | Mongolia_N_East | Mongolia_EIA_4_Sagly | -0.02262 | 0.000528 | -42.853 | 737092 |
|  | England_Bell_Beaker | Mongolia_N_East | Mongolia_EIA_4_Sagly | -0.02246 | 0.000529 | -42.414 | 732894 |
|  | CEU.SG | Mongolia_N_East | Mongolia_EIA_4_Sagly | -0.02147 | 0.000507 | -42.334 | 782275 |
|  | Russia_AfontovaGora3 | Mongolia_N_East | Mongolia_MBA_1_Munkhkhairkhan | -0.00988 | 0.00319 | -3.097 | 112225 |
|  | WSHG | Mongolia_N_East | Mongolia_MBA_1_Munkhkhairkhan | -0.0061 | 0.002393 | -2.55 | 374706 |
|  | Russia_MA1_HG.SG | Mongolia_N_East | Mongolia_MBA_1_Munkhkhairkhan | -0.00407 | 0.002745 | -1.481 | 277929 |
|  | Kazakhstan_Botai.SG | Mongolia_N_East | Mongolia_MBA_1_Munkhkhairkhan | -0.00318 | 0.002331 | -1.363 | 433825 |
|  | Russia_Karelia_HG.SG | Mongolia_N_East | Mongolia_MBA_1_Munkhkhairkhan | -0.00266 | 0.002666 | -0.996 | 334514 |
|  | Kazakhstan_Botai.SG | Mongolia_N_East | Mongolia_LBA_4_CenterWest | -0.01634 | 0.000876 | -18.648 | 625014 |
|  | WSHG | Mongolia_N_East | Mongolia_LBA_4_CenterWest | -0.01891 | 0.001021 | -18.52 | 524158 |
|  | Russia_Afanasievo | Mongolia_N_East | Mongolia_LBA_4_CenterWest | -0.01387 | 0.000782 | -17.744 | 706280 |
|  | Russia_Yamnaya_Samara | Mongolia_N_East | Mongolia_LBA_4_CenterWest | -0.01313 | 0.000748 | -17.545 | 726251 |
|  | Kazakhstan_Kumsay_EBA | Mongolia_N_East | Mongolia_LBA_4_CenterWest | -0.01604 | 0.000946 | -16.956 | 582655 |
| **top 5 negative *f3* values** | Kazakhstan_Solyanka_MLBA | Mongolia_N_East | Mongolia_LBA_3_MongunTaiga | -0.0275 | 0.001787 | -15.392 | 203771 |
|  | Russia_Potapovka_published | Mongolia_N_East | Mongolia_LBA_3_MongunTaiga | -0.02737 | 0.00181 | -15.122 | 205948 |
|  | Kazakhstan_Kyzlbulak_MLBA1 | Mongolia_N_East | Mongolia_LBA_3_MongunTaiga | -0.02736 | 0.001624 | -16.849 | 367353 |
|  | Latvia_LN_CW | Mongolia_N_East | Mongolia_LBA_3_MongunTaiga | -0.02735 | 0.001956 | -13.987 | 139325 |
|  | Russia_Sintashta_MLBA.SG | Mongolia_N_East | Mongolia_LBA_3_MongunTaiga | -0.0267 | 0.001266 | -21.086 | 445747 |
|  | Russia_Poltavka_lc | Mongolia_N_East | Mongolia_EIA_1_SlabGrave | -0.00767 | 0.00116 | -6.611 | 505347 |
|  | Latvia_MN_Comb_Ware.SG | Mongolia_N_East | Mongolia_EIA_1_SlabGrave | -0.00719 | 0.001323 | -5.43 | 384215 |
|  | Russia_AfontovaGora3 | Mongolia_N_East | Mongolia_EIA_1_SlabGrave | -0.00704 | 0.001603 | -4.393 | 155972 |
|  | Russia_MA1_HG.SG | Mongolia_N_East | Mongolia_EIA_1_SlabGrave | -0.00634 | 0.001353 | -4.684 | 422200 |
|  | Colonist.SG | Mongolia_N_East | Mongolia_EIA_1_SlabGrave | -0.00633 | 0.001912 | -3.312 | 101753 |
|  | Sweden_BA.SG | Mongolia_N_East | Mongolia_LBA_2_Ulaanzukh | -0.00387 | 0.002334 | -1.657 | 81159 |
|  | Kazakhstan_Kanai_MBA | Mongolia_N_East | Mongolia_LBA_2_Ulaanzukh | -0.00271 | 0.002433 | -1.113 | 65582 |
|  | Russia_Poltavka_lc | Mongolia_N_East | Mongolia_LBA_2_Ulaanzukh | -0.00238 | 0.001531 | -1.557 | 392759 |
|  | Latvia_LN_CW | Mongolia_N_East | Mongolia_LBA_2_Ulaanzukh | -0.0023 | 0.002047 | -1.123 | 135918 |
|  | Kazakhstan_Zevakinskiy_MLBA | Mongolia_N_East | Mongolia_LBA_2_Ulaanzukh | -0.00217 | 0.001633 | -1.33 | 336342 |
|  | Russia_AfontovaGora3 | Mongolia_N_East | Mongolia_LBA_6_Khovsgol | -0.02226 | 0.001562 | -14.25 | 163439 |
|  | WSHG | Mongolia_N_East | Mongolia_LBA_6_Khovsgol | -0.01949 | 0.001008 | -19.331 | 535913 |
|  | Russia_Samara_HG | Mongolia_N_East | Mongolia_LBA_6_Khovsgol | -0.01939 | 0.001367 | -14.189 | 278515 |
|  | Russia_MA1_HG.SG | Mongolia_N_East | Mongolia_LBA_6_Khovsgol | -0.0193 | 0.00132 | -14.627 | 443186 |
|  | Latvia_MN_Comb_Ware.SG | Mongolia_N_East | Mongolia_LBA_6_Khovsgol | -0.01859 | 0.001213 | -15.321 | 403611 |
|  | Russia_AfontovaGora3 | Mongolia_N_East | Mongolia_EBA_1_Ulgii | -0.01939 | 0.002204 | -8.8 | 124673 |
|  | Russia_MA1_HG.SG | Mongolia_N_East | Mongolia_EBA_1_Ulgii | -0.01811 | 0.001861 | -9.729 | 316068 |
|  | WSHG | Mongolia_N_East | Mongolia_EBA_1_Ulgii | -0.01806 | 0.001563 | -11.552 | 416120 |
|  | Russia_Poltavka_lc | Mongolia_N_East | Mongolia_EBA_1_Ulgii | -0.0176 | 0.00176 | -9.997 | 398163 |
|  | Latvia_MN_Comb_Ware.SG | Mongolia_N_East | Mongolia_EBA_1_Ulgii | -0.01734 | 0.001792 | -9.676 | 283350 |
|  | Kazakhstan_Solyanka_MLBA | Mongolia_N_East | Mongolia_EIA_4_Sagly | -0.02604 | 0.001182 | -22.026 | 265008 |
|  | Russia_Poltavka_lc | Mongolia_N_East | Mongolia_EIA_4_Sagly | -0.02493 | 0.0011 | -22.669 | 560582 |
|  | Russia_Sintashta_MLBA.SG | Mongolia_N_East | Mongolia_EIA_4_Sagly | -0.02463 | 0.000826 | -29.82 | 608444 |
|  | Kazakhstan_Aktogai_MLBA | Mongolia_N_East | Mongolia_EIA_4_Sagly | -0.02449 | 0.000693 | -35.35 | 652317 |
|  | England_Bell_Beaker_EBA | Mongolia_N_East | Mongolia_EIA_4_Sagly | -0.02435 | 0.001193 | -20.405 | 335246 |
|  | Russia_AfontovaGora3 | Mongolia_N_East | Mongolia_MBA_1_Munkhkhairkhan | -0.00988 | 0.00319 | -3.097 | 112225 |
|  | WSHG | Mongolia_N_East | Mongolia_MBA_1_Munkhkhairkhan | -0.0061 | 0.002393 | -2.55 | 374706 |
|  | Russia_MA1_HG.SG | Mongolia_N_East | Mongolia_MBA_1_Munkhkhairkhan | -0.00407 | 0.002745 | -1.481 | 277929 |
|  | Kazakhstan_Botai.SG | Mongolia_N_East | Mongolia_MBA_1_Munkhkhairkhan | -0.00318 | 0.002331 | -1.363 | 433825 |
|  | Russia_Karelia_HG.SG | Mongolia_N_East | Mongolia_MBA_1_Munkhkhairkhan | -0.00266 | 0.002666 | -0.996 | 334514 |
|  | Russia_AfontovaGora3 | Mongolia_N_East | Mongolia_LBA_4_CenterWest | -0.02045 | 0.001531 | -13.353 | 158638 |
|  | WSHG | Mongolia_N_East | Mongolia_LBA_4_CenterWest | -0.01891 | 0.001021 | -18.52 | 524158 |
|  | Russia_MA1_HG.SG | Mongolia_N_East | Mongolia_LBA_4_CenterWest | -0.01652 | 0.001346 | -12.279 | 429786 |
|  | Kazakhstan_Botai.SG | Mongolia_N_East | Mongolia_LBA_4_CenterWest | -0.01634 | 0.000876 | -18.648 | 625014 |
|  | Russia_Khvalynsk_EN | Mongolia_N_East | Mongolia_LBA_4_CenterWest | -0.0163 | 0.0011 | -14.82 | 441359 |

**Table S7. Mongolia_Neolithic-related admixture proportions in Neolithic and Early Bronze Age samples from Lake Baikal region estimated using *qpAdm*. Here “p” refers to the P-value for rank=1 and “std.err” is the standard error estimated using a block jackknife. We used Mbuti.DG, Ust_Ishim.DG, Russia_Kostenki14, Papuan.DG, Australian.DG, and Onge.DG as outgroups.**

|  | p-value | MA1 | Mongolia_N_East | std.err |
| --- | --- | --- | --- | --- |
| Russia_Lokomotiv_EN.SG | 0.419 | 0.099 | 0.901 | 0.036 |
| Russia_Shamanka_EBA.SG | 0.868 | 0.179 | 0.821 | 0.035 |
| Russia_Shamanka_EN.SG | 0.740 | 0.064 | 0.936 | 0.032 |
| Russia_UstIda_EBA.SG | 0.720 | 0.208 | 0.792 | 0.034 |
| Russia_UstIda_LN.SG | 0.354 | 0.229 | 0.771 | 0.033 |

**Table S8. Han Chinese-related admixture proportions (α) estimated using *qpAdm*. Here “p” refers to the P-value for rank=1 and “std.err” is the standard error estimated using a block jackknife. We selected Mbuti, Ust_Ishim, Kostenki14, Onge, Papuan, Australian, Nasioi, Mixe, Saqqaq, and Bichon as outgroups and Boisman, Han_Shandong and French as proxies for the source populations. We used Mixe and ancient Saqqaq as outgroups because we are interested in the relative proportions of Boisman and Han Chinese related ancestry, and wanted to populations on the right sensitive to Boisman ancestry.**

| **Target population** | **p** | **α** | **Std.err** |
| --- | --- | --- | --- |
| Xibo | 0.899 | 0.650 | 0.051 |
| Daur | 0.351 | 0.462 | 0.050 |
| Hezhen | 0.790 | 0.414 | 0.060 |
| Oroqen | 0.181 | 0.220 | 0.062 |
| Nanai | 0.279 | 0.129 | 0.068 |
| Negidal | 0.162 | -0.059 | 0.098 |
| Nivh | 0.168 | 0.086 | 0.084 |
| Ulchi | 0.171 | 0.009 | 0.072 |
| Yankovsky_IronAge | 0.121 | 0.113 | 0.146 |
| Heishui_Mohe_I1209_Medieval | 0.970 | 0.198 | 0.245 |
| Heishui_Mohe_I3358_Medieval | 0.205 | 0.570 | 0.150 |

**Table S9. Mixture proportions estimated in *qpAdm* using Core Tibetan as one source population and Tai-Kadai groups as the other. We chose Mbuti, Ju_hoan_North, French, BedouinB, Papuan, Onge, Mala and Nasioi as outgroups and Core Tibetans and Southeast Asian Cluster (here we use Tai-Kadai speaking populations in south China as the source, since they are geographically closer to Corridor populations) as proxies for the source populations. “p” is the P-value for rank 1, “α” refers to the proportion of gene flow from Southeast Asian Cluster, and the proportion from Tibetan related population could be calculated by “1-α”. The “std.err” is the standard error estimated using a Block Jackknife. Core Tibetan is a group combining all five Tibetan populations in Lhasa, Nagqu, Chamdo, Shannan, and Shigatse. We merged Maonan, Mulam, Zhuang, Dong_Guizhou, Dong_Hunan, Li, Gelao, CDX, and Dai into one group as Tai-Kadai with 193 individuals. We also used each Tai-Kadai speaking population alone as one source, and we found the results are qualitatively consistent with using the merged group.**

| **Population** | **p** | **α** | **std.err** |
| --- | --- | --- | --- |
| Lahu | 0.050 | 0.724 | 0.084 |
| Yi | 0.301 | 0.399 | 0.063 |
| Naxi | 0.121 | 0.339 | 0.073 |
| Qiang_Danba | 0.877 | 0.441 | 0.065 |
| Qiang_Daofu | 0.775 | 0.263 | 0.065 |
| Tibetan_Xinlong | 0.977 | 0.401 | 0.061 |
| Tibetan_Yajiang | 0.229 | -0.043 | 0.066 |
| Tibetan_Yunnan | 0.234 | 0.303 | 0.106 |

**Table S10. Populations with top 20 values of *f_3_* statistics (Mbuti; Core Tibetan, X), here “X” refers to worldwide ancient and modern populations except Tibetans in our Affymetrix Human Origins Dataset.**

| **Order** | | **Tibetan_Shigatse** | **Tibetan_Chamdo** | **Tibetan_Nagqu** | **Tibetan_Lhasa** | **Tibetan_Shannan** |
| --- | --- | --- | --- | --- | --- | --- |
|  | **1** | Wuzhuangguoliang | Wuzhuangguoliang | Wuzhuangguoliang | Wuzhuangguoliang | Wuzhuangguoliang |
|  | **2** | Qiang_Daofu | Qiang_Daofu | Qiang_Danba | Qiang_Daofu | Qiang_Daofu |
|  | **3** | Qiang_Danba | Qiang_Danba | Qiang_Daofu | Qiang_Danba | Qiang_Danba |
|  | **4** | Han_Shanghai | Han_Shanghai | Han_Shanghai | Han_Shanghai | Han_Shanghai |
|  | **5** | Yi | Yi | Yi | Yi | Yi |
|  | **6** | Han_Jiangsu | Naxi | Naxi | Naxi | Naxi |
|  | **7** | Han_Henan | Han_Henan | Han_Jiangsu | Han_Jiangsu | Han_Henan |
|  | **8** | Naxi | Han_Jiangsu | Han_Henan | Han_Henan | Han_Hubei |
|  | **9** | Han_Hubei | Han_Hubei | Han_Shandong | Han_Chongqing | Han_Jiangsu |
|  | **10** | Han_Chongqing | Korean | Han_Shanxi | Han_Hubei | Korean |
|  | **11** | Han_Shanxi | Han_Shanxi | Han_Hubei | Han_Shandong | Sherpa |
|  | **12** | CHB | CHB | Korean | Korean | Han_Shanxi |
|  | **13** | Han_Shandong | Han_Chongqing | Han_Chongqing | CHB | Han_Chongqing |
|  | **14** | Sherpa | Han_Shandong | CHB | Han_Shanxi | Han_Shandong |
|  | **15** | Korean | Han_Zhejiang | Yankovsky_IA | Sherpa | CHB |
|  | **16** | Han_Zhejiang | Han_Sichuan | Han_Zhejiang | Tujia | Han_Zhejiang |
|  | **17** | Han_Sichuan | Tujia | Han_Sichuan | Han_Zhejiang | Tujia |
|  | **18** | Han_NChina | Han_NChina | Tujia | Han_Sichuan | Han_Sichuan |
|  | **19** | Tujia | Miao | Han_HGDP | Han_NChina | Miao |
|  | **20** | Han_Fujian | She | Han_NChina | Miao | Yankovsky_IA |

**Table S11. ALDER test for admixture LD in Core Tibetans.** **We computed single-reference weighted LD using the following 8 groups with larger sample sizes as reference populations:**

**Han_HO: 13 Han Chinese populations with 107 individuals in our Human Origin Dataset--Han_Hubei, Han_Chongqing, Han_Shanghai, Han_Fujian, Han_HGDP, Han_Guangdong, Han_Shandong, Han_Shanxi, Han_Sichuan, Han_Jiangsu, Han_Zhejiang, Han_Henan, Han_NChina; CHB: CHB with 103 individuals from the 1000 Genome Project; CHS: CHS with 105 individuals from the 1000 Genome Project; Han_all: combined Han_HO, CHB, and CHS with a total of 315 individuals; Tai-Kadai: 9 Tai-Kadai populations with 193 individuals—Maonan, Mulam, Li, Gelao, Dong_Hunan, Dong_Guizhou, Zhuang, Dai, and CDX of 1000 Genome Project; Austronesian: Ami and Atayal, 19 individuals; Japanese: JPT of 1000 Genome Project and Japanese in our Human Origin Dataset, 133 individuals; Amur Basin: populations in Amur River Basin, including 56 individuals from Tungusic (Nanai, Negidal, Hezhen, and Ulchi) and Nivh; Siberian: 143 individuals from Altaian, Dolgan, Tubalar, Tuvinian, Yakut, Yukagir, Kalmyk, Chukchi, Koryak, and Eskimo.** **Core Tibetan is a group combining all five Tibetan populations in Lhasa, Nagqu, Chamdo, Shannan, and Shigatse.**

| **d(cM)** | **Han_HO** | **z-score** | **Mixture fraction % lower bound** |
| --- | --- | --- | --- |
| d>0.70 | 69.22 ± 19.44 | 3.56 | 22.3 ± 6.5 |
| d>0.80 | 64.59 ± 21.55 | 3.00 |  |
| d>0.90 | 60.15 ± 19.60 | 3.07 |  |
| d>1.00 | 61.96 ± 22.31 | 2.78 |  |
| d>1.10 | 63.93 ± 26.13 | 2.45 |  |
| **d(cM)** | **CHB** | **z-score** | **Mixture fraction % lower bound** |
| d>0.80 | 65.88 ± 20.33 | 3.24 | 25.0 ± 7.1 |
| d>0.90 | 64.25 ± 19.57 | 3.28 |  |
| d>1.00 | 61.56 ± 19.99 | 3.08 |  |
| d>1.10 | 65.28 ± 24.00 | 2.72 |  |
| d>1.20 | 61.77 ± 20.99 | 2.94 |  |
| **d(cM)** | **CHS** | **z-score** | **Mixture fraction % lower bound** |
| d>0.60 | 77.93 ± 18.51 | 4.21 | 20.0 ± 5.6 |
| d>0.70 | 75.11 ± 19.37 | 3.88 |  |
| d>0.80 | 67.70 ± 19.75 | 3.43 |  |
| d>0.90 | 66.85 ± 19.93 | 3.35 |  |
| d>1.00 | 67.30 ± 21.23 | 3.17 |  |
| **d(cM)** | **Han_all** | **z-score** | **Mixture fraction % lower bound** |
| d>0.80 | 66.23 ± 19.92 | 3.33 | 22.8 ± 7.0 |
| d>0.90 | 64.11 ± 19.25 | 3.33 |  |
| d>1.00 | 64.08 ± 20.72 | 3.09 |  |
| d>1.10 | 67.51 ± 24.98 | 2.70 |  |
| d>1.20 | 63.27 ± 22.15 | 2.86 |  |
| **d(cM)** | **Tai-Kadai** | **z-score** | **Mixture fraction % lower bound** |
| d>0.70 | 85.35 ± 39.18 | 2.18 | 9.9 ± 6.7 |
| d>0.80 | 70.60 ± 44.44 | 1.59 |  |
| d>0.90 | 68.03 ± 45.25 | 1.50 |  |
| d>1.00 | 64.10 ± 48.53 | 1.32 |  |
| d>1.10 | 56.00 ± 40.14 | 1.40 |  |
| **d(cM)** | **Austronesian** | **z-score** | **Mixture fraction % lower bound** |
| d>0.30 | 183.03 ± 70.75 | 2.59 | 4.8 ± 4.9 |
| d>0.40 | 137.78 ± 52.06 | 2.65 |  |
| d>0.50 | 119.25 ± 131.75 | 0.91 |  |
| d>0.60 | 27.61 ± 28.34 | 0.97 |  |
| d>0.70 | 23.60 ± 19.08 | 1.24 |  |
| **d(cM)** | **Japanese** | **z-score** | **Mixture fraction % lower bound** |
| d>0.80 | 66.24 ± 18.30 | 3.62 | 16.2 ± 4.3 |
| d>0.90 | 67.23 ± 18.73 | 3.59 |  |
| d>1.00 | 72.38 ± 18.52 | 3.91 |  |
| d>1.10 | 79.73 ± 17.17 | 4.64 |  |
| d>1.20 | 86.25 ± 20.20 | 4.27 |  |
| **d(cM)** | **Amur Basin** | **z-score** | **Mixture fraction % lower bound** |
| d>0.60 | 49.88 ± 19.27 | 2.59 | 5.2 ± 1.8 |
| d>0.70 | 48.45 ± 17.55 | 2.76 |  |
| d>0.80 | 50.57 ± 17.94 | 2.82 |  |
| d>0.90 | 54.24 ± 18.27 | 2.97 |  |
| d>1.00 | 66.99 ± 23.28 | 2.88 |  |
| **d(cM)** | **Siberian** | **z-score** | **Mixture fraction % lower bound** |
| d>0.80 | 35.86 ± 8.81 | 4.07 | 5.6 ± 1.1 |
| d>0.90 | 36.67 ± 8.66 | 4.23 |  |
| d>1.00 | 37.94 ± 8.68 | 4.37 |  |
| d>1.10 | 37.69 ± 8.20 | 4.60 |  |
| d>1.20 | 36.35 ± 7.61 | 4.78 |  |

**Table S12: Statistics of the form of *f_3_* (Han Chinese; X, Y), where X and Y are worldwide populations. Here we only list the top 3 negative values for each Han Chinese population.**

| **Source 1** | **Source 2** | **Target** | ***f_3_*** | **std.err** | **Z** |
| --- | --- | --- | --- | --- | --- |
| CDX | Nanai | CHB | -0.00256 | 0.000137 | -18.683 |
| CDX | Ulchi | CHB | -0.00221 | 0.000124 | -17.851 |
| Mulam | Nanai | CHB | -0.00292 | 0.000173 | -16.863 |
| CDX | Nanai | CHS | -0.00246 | 0.000121 | -20.315 |
| CDX | Ulchi | CHS | -0.00219 | 0.000112 | -19.489 |
| Maonan | Ulchi | CHS | -0.00278 | 0.000171 | -16.223 |
| Ami | Tibetan_Chamdo | Han_Chongqing | -0.00461 | 0.000859 | -5.362 |
| Ami | Tibetan_Lhasa | Han_Chongqing | -0.00445 | 0.000855 | -5.203 |
| Atayal | Tibetan_Lhasa | Han_Chongqing | -0.00480 | 0.000926 | -5.183 |
| Ami | Tibetan_Chamdo | Han_Fujian | -0.00325 | 0.000567 | -5.726 |
| Ami | Tibetan_Shigatse | Han_Fujian | -0.00322 | 0.000567 | -5.685 |
| Ami | Qiang_Daofu | Han_Fujian | -0.00283 | 0.000562 | -5.040 |
| Ami | Tibetan_Chamdo | Han_Guangdong | -0.00384 | 0.000444 | -8.632 |
| Maonan | Tibetan_Chamdo | Han_Guangdong | -0.00286 | 0.000377 | -7.594 |
| Li | Tibetan_Chamdo | Han_Guangdong | -0.00353 | 0.000476 | -7.410 |
| Ami | Tibetan_Chamdo | Han_Henan | -0.00423 | 0.000574 | -7.372 |
| Atayal | Tibetan_Chamdo | Han_Henan | -0.00431 | 0.000593 | -7.266 |
| Li | Tibetan_Chamdo | Han_Henan | -0.00381 | 0.000557 | -6.847 |
| CDX | Japanese | Han_HGDP | -0.00184 | 0.000102 | -18.063 |
| Maonan | Tibetan_Chamdo | Han_HGDP | -0.00319 | 0.000181 | -17.594 |
| Zhuang | Tibetan_Chamdo | Han_HGDP | -0.00282 | 0.000167 | -16.911 |
| Ami | Tibetan_Chamdo | Han_Hubei | -0.00480 | 0.000549 | -8.738 |
| Atayal | Tibetan_Chamdo | Han_Hubei | -0.00447 | 0.000575 | -7.773 |
| Ami | Tibetan_Gangcha | Han_Hubei | -0.00394 | 0.000511 | -7.703 |
| Ami | Tibetan_Chamdo | Han_Jiangsu | -0.00405 | 0.000450 | -9.003 |
| Ami | Tibetan_Gangcha | Han_Jiangsu | -0.00314 | 0.000418 | -7.497 |
| Maonan | Tibetan_Chamdo | Han_Jiangsu | -0.00281 | 0.000382 | -7.363 |
| Dai | Tibetan_Chamdo | Han_NChina | -0.00256 | 0.000330 | -7.749 |
| Mulam | Yakut | Han_NChina | -0.00223 | 0.000316 | -7.058 |
| Dai | Nanai | Han_NChina | -0.00239 | 0.000348 | -6.857 |
| Ami | Tibetan_Chamdo | Han_Shandong | -0.00309 | 0.000377 | -8.217 |
| Ami | Tibetan_Nagqu | Han_Shandong | -0.00302 | 0.000404 | -7.464 |
| Mulam | Nanai | Han_Shandong | -0.00234 | 0.000332 | -7.037 |
| Atayal | Tibetan_Chamdo | Han_Shanghai | -0.00290 | 0.001386 | -2.095 |
| Ami | Tibetan_Chamdo | Han_Shanghai | -0.00286 | 0.001398 | -2.045 |
| Ami | Tibetan_Xunhua | Han_Shanghai | -0.00295 | 0.001550 | -1.902 |
| Ami | Tibetan_Chamdo | Han_Shanxi | -0.00281 | 0.000426 | -6.606 |
| Dong_Guizhou | Yakut | Han_Shanxi | -0.00230 | 0.000372 | -6.183 |
| Mulam | Yakut | Han_Shanxi | -0.00221 | 0.000376 | -5.876 |
| Ami | Tibetan_Chamdo | Han_Sichuan | -0.00498 | 0.000425 | -11.709 |
| Maonan | Tibetan_Chamdo | Han_Sichuan | -0.00404 | 0.000369 | -10.955 |
| Ami | Tibetan_Shigatse | Han_Sichuan | -0.00446 | 0.000425 | -10.481 |
| Ami | Tibetan_Chamdo | Han_Zhejiang | -0.00376 | 0.000558 | -6.729 |
| Ami | Tibetan_Shigatse | Han_Zhejiang | -0.00347 | 0.000584 | -5.941 |
| Ami | Tibetan_Nagqu | Han_Zhejiang | -0.00338 | 0.000601 | -5.618 |

**Table S13. *f_4_*-statistics highlight mixture in Han Chinese. Significant values are highlighted. The top panel uses all samples. The bottom panel only uses Han, Tibetan_Chamdo and Ulchi individuals with *f_4_* (French, Mbuti; X, Ami) consistent with zero |Z|<2 (40 Han and 16 Ulchi are removed).**

| **Population** | **Ulchi shares more alleles with Han Chinese than with Ami or Tibetan** | | **Tibetan shares more alleles with Han Chinese than with Ami or Ulchi** | | **Ami shares more alleles with Han Chinese than with Ulchi or Tibetan** | | **Ami shares more alleles with Han Chinese than with Wuzhuangguoliang** |
| --- | --- | --- | --- | --- | --- | --- | --- |
|  | ***f_4_*(Ulchi, Mbuti;**  **Tibetan_Chamdo, Han)** | ***f_4_*(Ulchi, Mbuti;**  **Ami, Han)** | ***f_4_*(Tibetan_Chamdo,**  **Mbuti; Ami, Han)** | ***f_4_*(Tibetan_Chamdo,**  **Mbuti; Ulchi, Han)** | ***f_4_*(Ami, Mbuti;**  **Ulchi, Han)** | ***f_4_*(Ami, Mbuti;**  **Tibetan_Chamdo, Han)** | ***f4*(Ami, Mbuti; Han, Wuzhuangguoliang)** |
| **Han_Henan** | -8.297 | -7.465 | -10.010 | -8.267 | -11.621 | -14.868 | 2.096 |
| **Han_Shandong** | -7.645 | -7.330 | -10.196 | -7.174 | -13.305 | -17.135 | 2.549 |
| **Han_Shanxi** | -7.608 | -7.007 | -9.992 | -7.328 | -11.423 | -14.908 | 1.737 |
| **Han_NChina** | -5.151 | -4.947 | -8.697 | -5.984 | -10.159 | -13.605 | 1.344 |
| **CHB** | -9.488 | -8.221 | -12.060 | -9.250 | -19.318 | -24.759 | 3.601 |
| **Han_Jiangsu** | -8.39 | -7.954 | -11.163 | -8.427 | -16.660 | -20.118 | 3.585 |
| **Han_Shanghai** | -7.077 | -7.213 | -9.926 | -7.601 | -12.460 | -15.204 | 3.623 |
| **Han_Sichuan** | -4.196 | -4.620 | -8.916 | -5.724 | -15.575 | -19.752 | 3.786 |
| **Han_Zhejiang** | -5.693 | -6.173 | -8.888 | -5.811 | -15.565 | -18.630 | 3.013 |
| **Han_Fujian** | -4.382 | -5.012 | -7.419 | -3.946 | -17.398 | -20.235 | 4.590 |
| **Han_Hubei** | -7.043 | -7.285 | -10.565 | -7.830 | -17.777 | -20.301 | 4.355 |
| **Han_Chongqing** | -4.815 | -5.427 | -8.576 | -5.454 | -14.346 | -16.887 | 4.568 |
| **CHS** | -5.984 | -6.255 | -10.288 | -6.111 | -24.102 | -29.305 | 5.221 |
| **Han_HGDP** | -5.968 | -6.122 | -9.624 | -6.057 | -22.212 | -26.789 | 4.427 |
| **Han_Guangdong** | -2.092 | -2.990 | -6.651 | -2.976 | -20.998 | -24.472 | 4.982 |
| **Han_Henan** | -6.363 | -5.680 | -8.547 | -7.274 | -9.708 | -12.807 | 1.925 |
| **Han_Shandong** | -7.334 | -6.623 | -9.903 | -7.489 | -12.324 | -16.732 | 2.567 |
| **Han_Shanxi** | -6.134 | -5.821 | -8.949 | -7.047 | -9.735 | -11.941 | 1.878 |
| **Han_NChina** | -4.242 | -3.909 | -7.180 | -5.406 | -8.414 | -11.004 | 0.704 |
| **CHB** | -8.761 | -7.243 | -11.566 | -9.051 | -17.374 | -24.428 | 4.247 |
| **Han_Jiangsu** | -8.29 | -7.416 | -11.163 | -8.775 | -15.423 | -20.118 | 3.585 |
| **Han_Sichuan** | -3.851 | -3.865 | -8.203 | -5.84 | -14.59 | -19.088 | 3.936 |
| **Han_Zhejiang** | -5.893 | -6.040 | -8.888 | -6.693 | -15.13 | -18.63 | 3.013 |
| **Han_Fujian** | -4.613 | -4.829 | -7.419 | -4.969 | -17.031 | -20.235 | 4.590 |
| **Han_Hubei** | -6.142 | -6.048 | -9.784 | -7.725 | -15.592 | -18.705 | 4.237 |
| **Han_Chongqing** | -4.939 | -5.195 | -8.576 | -6.182 | -13.849 | -16.887 | 4.568 |
| **CHS** | -5.843 | -5.572 | -10.148 | -6.785 | -21.376 | -29.01 | 5.856 |
| **Han_HGDP** | -5.922 | -5.52 | -9.534 | -6.751 | -19.985 | -26.866 | 4.831 |
| **Han_Guangdong** | -2.207 | -2.624 | -6.651 | -4.096 | -20.081 | -24.472 | 4.982 |

**Table S14: The number of source populations for Han Chinese inferred from *qpWave*.** **As a check on the robustness of this analysis, we also dropped each outgroup and Han Chinese population to determine that the number of source populations is not driven by one outgroup with a specific affinity to the tested populations or by a Han group having unusual ancestry. Values of P < 0.05 are highlighted. The source populations are particularly related to outgroup Karitiana, Papuan and Nasioi. When we drop Han_NChina which has West Eurasian-related admixture, 2-sources (rank1) are enough to explain the variations of Han Chinese.**

| **Dropped** | **all outgroups (no drop)** | | **Mbuti** | | **BedouinB** | | **Chechen** | | **Kalash** | | **Karitiana** | | **Papuan** | | **Nasioi** | |
| --- | --- | --- | --- | --- | --- | --- | --- | --- | --- | --- | --- | --- | --- | --- | --- | --- |
|  | **rank1** | **rank2** | **rank1** | **rank2** | **rank1** | **rank2** | **rank1** | **rank2** | **rank1** | **rank2** | **rank1** | **rank2** | **rank1** | **rank2** | **rank1** | **rank2** |
| **Han (no drop)** | 1.608E-02 | 4.938E-01 | 9.728E-03 | 4.513E-01 | 9.931E-03 | 2.774E-01 | 1.770E-02 | 4.539E-01 | 7.939E-03 | 3.339E-01 | 1.955E-01 | 7.851E-01 | 1.701E-01 | 6.993E-01 | 5.400E-02 | 6.144E-01 |
| **Han_Henan** | 2.025E-02 | 4.914E-01 | 1.009E-02 | 4.264E-01 | 1.556E-02 | 3.043E-01 | 9.163E-03 | 3.753E-01 | 6.583E-03 | 2.889E-01 | 2.258E-01 | 8.229E-01 | 1.572E-01 | 7.237E-01 | 4.430E-02 | 6.155E-01 |
| **Han_Shandong** | 4.743E-02 | 6.496E-01 | 4.041E-02 | 6.671E-01 | 2.332E-02 | 4.654E-01 | 5.287E-02 | 6.524E-01 | 2.981E-02 | 5.165E-01 | 3.990E-01 | 8.727E-01 | 1.037E-01 | 6.306E-01 | 3.544E-02 | 5.286E-01 |
| **Han_Shanxi** | 1.569E-02 | 4.823E-01 | 8.634E-03 | 4.321E-01 | 1.167E-02 | 2.772E-01 | 1.800E-02 | 4.799E-01 | 7.116E-03 | 3.272E-01 | 2.168E-01 | 7.541E-01 | 1.327E-01 | 6.600E-01 | 6.859E-02 | 6.664E-01 |
| **Han_NChina** | 1.755E-01 | 5.613E-01 | 1.105E-01 | 6.242E-01 | 1.409E-01 | 3.221E-01 | 1.752E-01 | 5.093E-01 | 9.275E-02 | 4.218E-01 | 3.157E-01 | 7.784E-01 | 4.659E-01 | 7.739E-01 | 2.917E-01 | 6.591E-01 |
| **CHB** | 1.335E-02 | 4.984E-01 | 9.231E-03 | 4.736E-01 | 9.274E-03 | 3.065E-01 | 1.591E-02 | 4.390E-01 | 6.218E-03 | 3.446E-01 | 1.231E-01 | 7.072E-01 | 2.293E-01 | 6.737E-01 | 8.700E-02 | 6.498E-01 |
| **Han_Sichuan** | 1.282E-02 | 4.263E-01 | 7.172E-03 | 3.927E-01 | 9.359E-03 | 2.532E-01 | 1.458E-02 | 3.941E-01 | 7.349E-03 | 2.939E-01 | 1.357E-01 | 7.440E-01 | 1.528E-01 | 6.478E-01 | 4.701E-02 | 5.722E-01 |
| **Han_Zhejiang** | 1.537E-02 | 4.056E-01 | 9.339E-03 | 3.504E-01 | 5.072E-03 | 1.862E-01 | 1.654E-02 | 3.707E-01 | 8.515E-03 | 2.664E-01 | 1.689E-01 | 7.657E-01 | 1.614E-01 | 6.654E-01 | 6.057E-02 | 5.892E-01 |
| **Han_Fujian** | 8.360E-03 | 3.931E-01 | 4.929E-03 | 3.565E-01 | 4.298E-03 | 1.960E-01 | 7.974E-03 | 3.430E-01 | 4.410E-03 | 2.683E-01 | 1.463E-01 | 7.114E-01 | 1.182E-01 | 6.438E-01 | 3.493E-02 | 5.736E-01 |
| **CHS** | 9.728E-02 | 8.556E-01 | 7.379E-02 | 8.673E-01 | 7.034E-02 | 7.393E-01 | 7.598E-02 | 7.474E-01 | 6.981E-02 | 8.028E-01 | 1.683E-01 | 7.837E-01 | 4.900E-01 | 8.961E-01 | 3.498E-01 | 9.651E-01 |
| **Han_Chongqing** | 8.628E-03 | 4.203E-01 | 6.501E-03 | 4.431E-01 | 5.888E-03 | 2.389E-01 | 1.040E-02 | 4.200E-01 | 3.534E-03 | 2.566E-01 | 1.417E-01 | 7.542E-01 | 1.230E-01 | 6.310E-01 | 3.470E-02 | 5.224E-01 |
| **Han_HGDP** | 9.949E-03 | 4.455E-01 | 6.200E-03 | 4.315E-01 | 5.801E-03 | 2.478E-01 | 8.687E-03 | 3.803E-01 | 6.827E-03 | 3.379E-01 | 1.458E-01 | 6.765E-01 | 1.275E-01 | 6.645E-01 | 5.425E-02 | 6.581E-01 |
| **Han_Guangdong** | 1.616E-02 | 4.597E-01 | 1.303E-02 | 4.542E-01 | 7.561E-03 | 2.236E-01 | 1.910E-02 | 4.335E-01 | 1.018E-02 | 3.372E-01 | 2.327E-01 | 7.177E-01 | 1.190E-01 | 6.956E-01 | 4.083E-02 | 6.339E-01 |
| **Han_Hubei** | 6.466E-02 | 5.004E-01 | 2.843E-02 | 4.460E-01 | 3.415E-02 | 3.229E-01 | 6.859E-02 | 5.473E-01 | 4.579E-02 | 3.925E-01 | 2.606E-01 | 8.284E-01 | 3.785E-01 | 8.218E-01 | 1.931E-01 | 5.891E-01 |
| **Han_Shanghai** | 4.294E-02 | 6.052E-01 | 2.984E-02 | 4.202E-01 | 3.038E-02 | 4.618E-01 | 3.811E-02 | 6.041E-01 | 1.915E-02 | 4.365E-01 | 4.738E-01 | 9.318E-01 | 2.810E-01 | 8.241E-01 | 6.482E-02 | 6.169E-01 |
| **Han_Jiangsu** | 7.813E-03 | 4.242E-01 | 4.227E-03 | 3.826E-01 | 4.057E-03 | 2.156E-01 | 9.375E-03 | 4.074E-01 | 3.754E-03 | 2.469E-01 | 1.311E-01 | 7.210E-01 | 1.089E-01 | 6.220E-01 | 3.280E-02 | 5.619E-01 |

**Table S15: The number of sources for all Han Chinese groups and French inferred from *qpWave*. We drop two outgroups at a time. The values in the bottom left are *P* values for rank 1, and the top right are *P* values for rank 2. We highlight all scores at P<0.01. The analyses indicate that at least three sources of ancestry are needed to explain the joint set of Han Chinese and French, which is no more than the three sources that are needed to explain Han Chinese alone. The fact that adding French into the set of tested populations does not increase the required number of sources of ancestry supports the hypothesis that one of the sources of ancestry is West Eurasian-related, which as shown in Table S14, has contributed at a low level to Han Chinese populations especially in northern China. After accounting for this ancestry, diverse Han Chinese appear to be consistent with deriving from just two sources of ancestry.**

| Dropped | Mbuti | BedouinB | Chechen | Kalash | Karitiana | Papuan | Nasioi |
| --- | --- | --- | --- | --- | --- | --- | --- |
| Mbuti |  | 8.076E-02 | 2.221E-01 | 1.895E-01 | 2.822E-01 | 5.736E-01 | 4.614E-01 |
| BedouinB | 6.972E-21 |  | 2.397E-01 | 2.154E-01 | 1.933E-01 | 4.163E-01 | 2.873E-01 |
| Chechen | 4.770E-21 | 1.605E-19 |  | 2.149E-01 | 4.435E-01 | 7.541E-01 | 7.324E-01 |
| Kalash | 1.755E-19 | 2.804E-19 | 5.370E-19 |  | 7.365E-01 | 6.612E-01 | 4.911E-01 |
| Karitiana | 2.387E-04 | 6.516E-03 | 1.011E-03 | 4.301E-03 |  | 4.909E-01 | 2.513E-01 |
| Papuan | 8.146E-19 | 1.016E-18 | 8.293E-19 | 1.565E-16 | 6.246E-02 |  | 7.115E-01 |
| Nasioi | 3.241E-08 | 2.210E-10 | 6.603E-09 | 9.799E-09 | 1.173E-01 | 1.443E-03 |  |

**Table S16: Statistics of the form of *f_3_* (Japanese; Japan_Jomon, X), where X are worldwide populations. Here we only list the top 30 negative *f_3_* values.**

| **Pop1** | **Pop2** | **Target** | ***f_3_*** | **std.err** | **Z** | **SNP** |
| --- | --- | --- | --- | --- | --- | --- |
| Japan_Jomon | Han_Zhejiang | Japanese | -0.01295 | 0.000622 | -20.820 | 352038 |
| Japan_Jomon | Han_Shandong | Japanese | -0.01283 | 0.000495 | -25.911 | 355396 |
| Japan_Jomon | Han_Shanghai | Japanese | -0.01276 | 0.00086 | -14.844 | 350092 |
| Japan_Jomon | CHB.SG | Japanese | -0.01257 | 0.000367 | -34.229 | 369512 |
| Japan_Jomon | Han_Henan | Japanese | -0.01248 | 0.000626 | -19.922 | 352728 |
| Japan_Jomon | Han_Jiangsu | Japanese | -0.01233 | 0.00057 | -21.622 | 353044 |
| Japan_Jomon | Korean | Japanese | -0.01232 | 0.000588 | -20.939 | 352576 |
| Japan_Jomon | Han_Hubei | Japanese | -0.01228 | 0.000606 | -20.251 | 352233 |
| Japan_Jomon | Han_Shanxi | Japanese | -0.0122 | 0.00054 | -22.595 | 354952 |
| Japan_Jomon | Han_Chongqing | Japanese | -0.01202 | 0.000765 | -15.715 | 351001 |
| Japan_Jomon | Han_HGDP | Japanese | -0.01189 | 0.000404 | -29.45 | 363573 |
| Japan_Jomon | She | Japanese | -0.01179 | 0.000532 | -22.174 | 354801 |
| Japan_Jomon | Qiang_Danba | Japanese | -0.01165 | 0.000536 | -21.743 | 355798 |
| Japan_Jomon | CHS.SG | Japanese | -0.01165 | 0.000376 | -30.975 | 366956 |
| Japan_Jomon | Hakka_Taiwan | Japanese | -0.01159 | 0.000551 | -21.028 | 349474 |
| Japan_Jomon | Han_Sichuan | Japanese | -0.01153 | 0.00054 | -21.336 | 353822 |
| Japan_Jomon | Xibo | Japanese | -0.01148 | 0.00056 | -20.478 | 356740 |
| Japan_Jomon | Han_Taiwan | Japanese | -0.01145 | 0.000499 | -22.943 | 350320 |
| Japan_Jomon | Hezhen | Japanese | -0.01139 | 0.000546 | -20.869 | 356310 |
| Japan_Jomon | Han_NChina | Japanese | -0.01129 | 0.000487 | -23.166 | 357393 |
| Japan_Jomon | Daur | Japanese | -0.01119 | 0.000516 | -21.667 | 358517 |
| Japan_Jomon | Tujia | Japanese | -0.0111 | 0.000521 | -21.308 | 356118 |
| Japan_Jomon | Han_Fujian | Japanese | -0.01098 | 0.000621 | -17.676 | 352386 |
| Japan_Jomon | Yugur | Japanese | -0.01092 | 0.000441 | -24.746 | 364642 |
| Japan_Jomon | Mongola | Japanese | -0.01084 | 0.000555 | -19.524 | 356569 |
| Japan_Jomon | Han_Guangdong | Japanese | -0.01064 | 0.000551 | -19.312 | 354168 |
| Japan_Jomon | Tibetan_Xinlong | Japanese | -0.0106 | 0.000518 | -20.455 | 357729 |
| Japan_Jomon | Mongolia_XiongNu.SG | Japanese | -0.01055 | 0.00084 | -12.556 | 352821 |
| Japan_Jomon | Wuzhuangguoliang | Japanese | -0.0105 | 0.001742 | -6.029 | 46331 |
| Japan_Jomon | Tibetan_Yajiang | Japanese | -0.01048 | 0.000539 | -19.436 | 360260 |

**Table S17. The Jomon related ancestry in present-day Japanese and Korean estimated from *qpAdm* modelling. We used CHB and Jomon as two sources and the following populations as outgroups: Mbuti.DG, Ust_Ishim.DG, Russia_Kostenki14, Papuan.DG, Australian.DG, Onge.DG, Tianyuan, Yana_UP.SG, Atayal.DG.**

| **Population** | **p** | **Source1** | **Source2** | **std.err** |
| --- | --- | --- | --- | --- |
|  |  | **CHB** | **Jomon** |  |
| Japanese | 0.095 | 0.843 | 0.157 | 0.019 |
| Korean | 0.079 | 0.958 | 0.042 | 0.031 |
|  |  | **Korean** | **Jomon** |  |
| Japanese | 0.457 | 0.876 | 0.124 | 0.031 |

**Table S18. Modeling present-day Japanese as an admixture of three sources. We used CHB, Korean and Jomon as three sources and the following populations as outgroups:** **Mbuti.DG, Ust_Ishim.DG, Russia_Kostenki14, Papuan.DG, Australian.DG, Onge.DG, Tianyuan, Yana_UP.SG, Atayal, Tibetan.DG, Li.**

|  | p | CHB | | Korean | | Japan_Jomon | |
| --- | --- | --- | --- | --- | --- | --- | --- |
|  |  | proportion | std.err | proportion | std.err | proportion | std.err |
| Japanese | 0.478 | 0.446 | 0.141 | 0.429 | 0.144 | 0.125 | 0.011 |

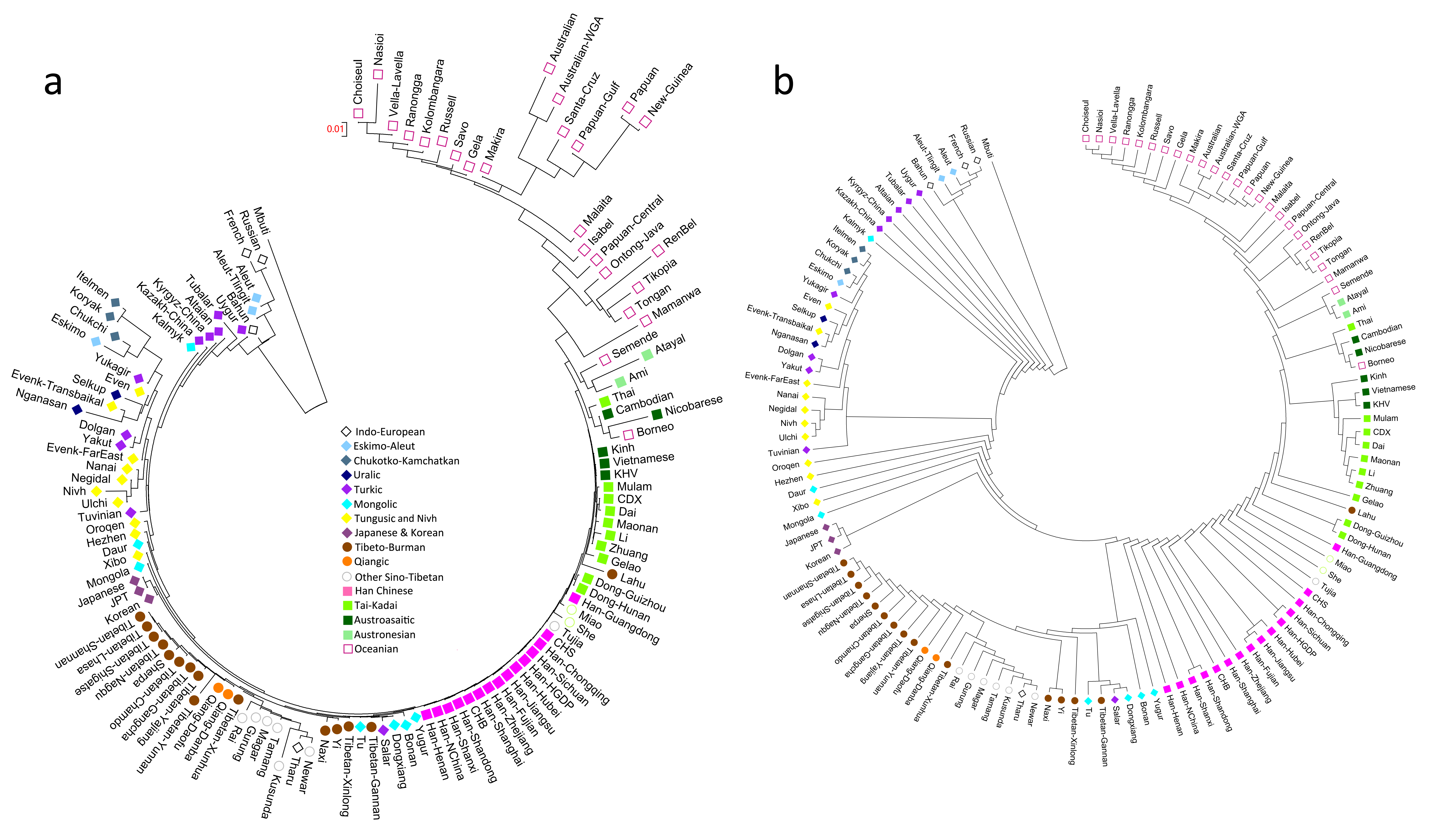

**Figure S1. Neighbor-joining tree of East Eurasians based on F_st_ distances using the Human Origin dataset. (a)** **The branch length is shown in F_st_ distance, (b) Version of the same figure where internal branches are all shown as having the same length for better visualization.**

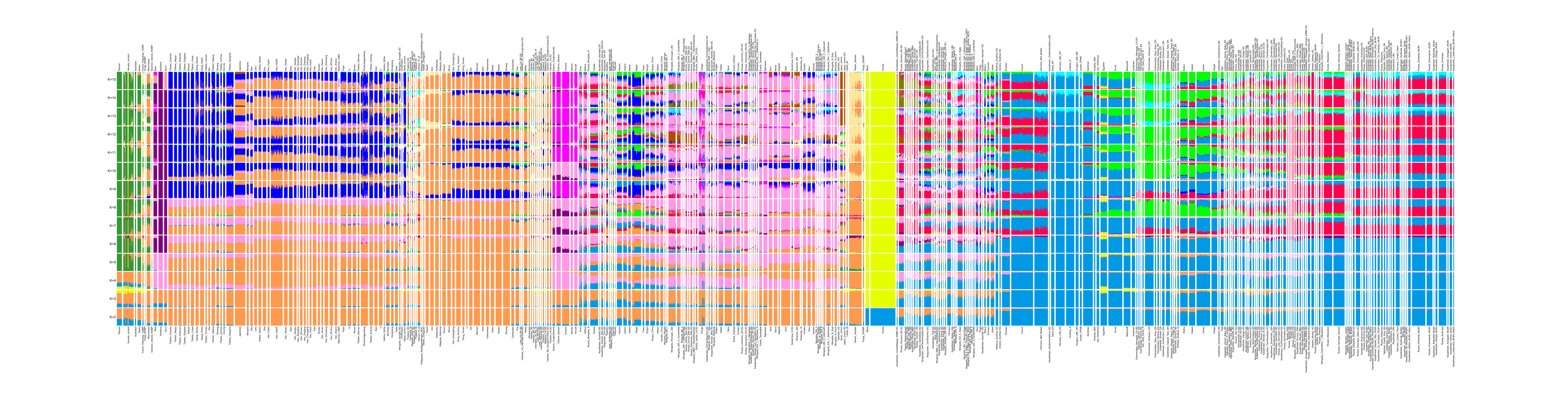

**Figure S2. Admixture plot from K=2 to K=15 using the Human Origin dataset**

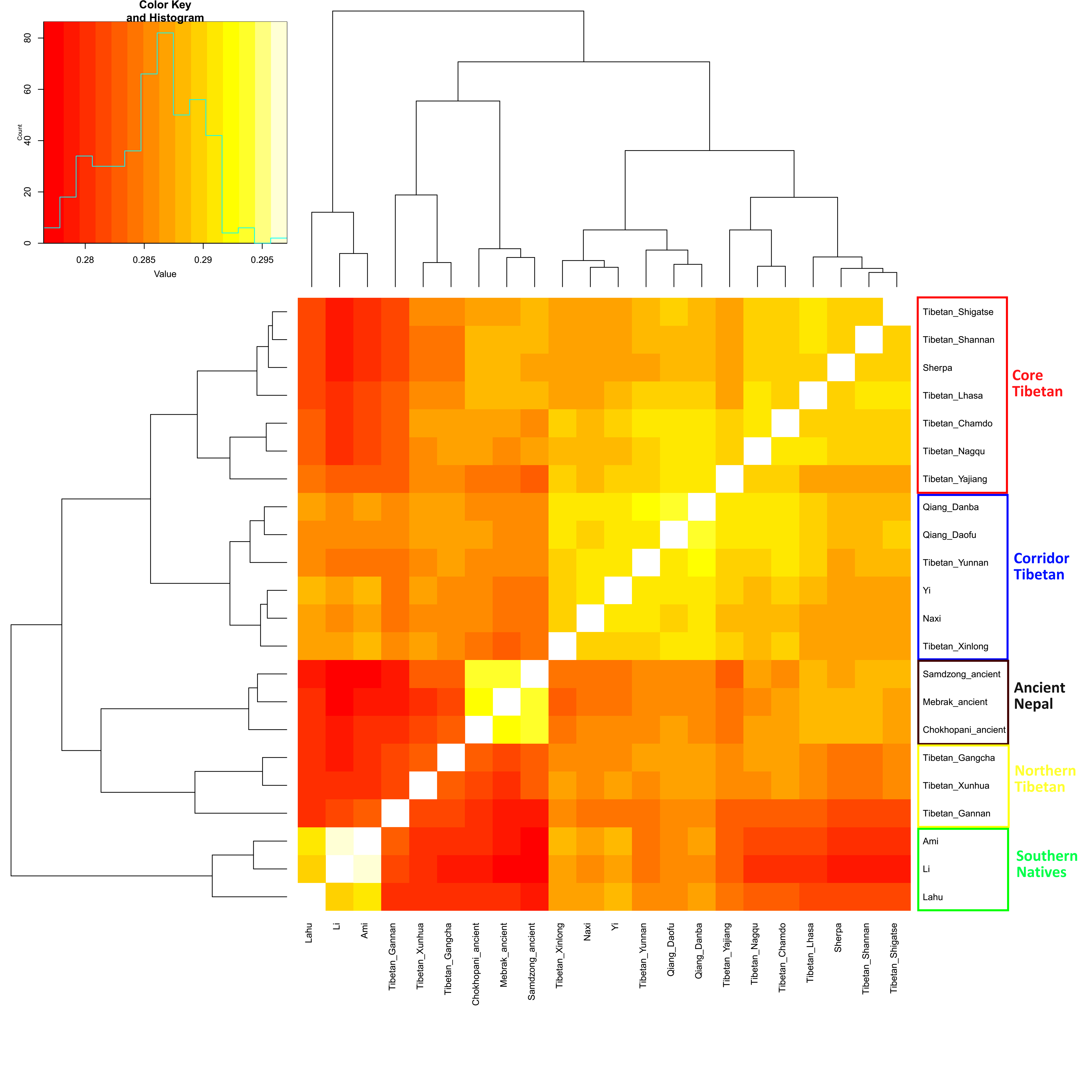

**Figure S3: Shared genetic drift among Tibetans, measured by *f_3_* (X, Y; Mbuti). Lighter colors indicate more shared drift. Lahu group with the Southeast Asian Cluster probably due to the substantial gene flow. The Tibetan_Yajiang are geographically in the Tibeto-Burman Corridor but group with Core Tibetans, presumably reflecting less genetic admixture from the Southeast Asian Cluster.**
