## Supplementary Information sections 1-3 for "The Genomic Formation of Human Populations in East Asia"

Supplementary Information section 1: Context of newly reported individuals

Supplementary Information section 2: Overview of genetic substructure

Supplementary Information section 3: Admixture graph modeling

**Supplementary Information section 1: Context of newly reported individuals**

**Newly reported samples from present-day groups**

We collected blood samples from 383 healthy and unrelated individuals from Han Chinese, Tibeto-Burman, Tai-Kadai, Austroasiatic, Mongolic, Turkic, Tungusic, and Indo-European speaking groups from China and Nepal. Table S1 lists information on these samples. The sample collection was carried out in 2014 in accordance with the human ethical research principles of The Ministry of Science and Technology of the People’s Republic of China (Interim Measures for the Administration of Human Genetic Resources, June 10, 1998) and genotyping of the samples was reviewed and approved by the Ethics Committee of the School of Life Sciences, Fudan University. Study staff informed potential participants about the goals of the project, and individuals who chose to participate gave informed consent consistent with broad studies of population history. Genomic DNA was extracted using the QIAamp DNA Mini Kit (QIAGEN, Hilden, Germany). The individuals all contributed DNA samples voluntarily, and the informed consents of all newly reported individuals are consistent with the full public dissemination of de-identified genotype data. The collection of genome-wide variation data on de-identified samples was approved by the Harvard Human Research Protection Program (Protocol 11681), re-reviewed on 12 July 2016. We genotyped all the samples on the Affymetrix Human Origins single nucleotide polymorphism (SNP) array at the Shenzhen-Beijing Genomics Institute.

**Ancient samples**

We newly report data from 191 ancient skeletons including 89 from 43 sites in Mongolia from ~6000 BCE to ~1000 CE, 20 ~3000 BCE individuals from Shannxi in northern China (Neolithic farmers from the Wuzhuangguoliang site), 52 from Taiwan (Iron Age and earlier samples from the Hanben and Gongguan sites), 18 from the Amur River Basin in the Russian Far East dating to around ~5000 BCE (Middle Neolithic hunter-gatherers from the Boisman-2 Cemetery), one ~1000 BCE individual (Iron Age from the Yankovsky Culture), three early Medieval individuals dating to ~1000 CE (from the Heishui Mohe and Bohai Mohe culture), 7 Jomon individuals from about 1500-1000 BCE and 3500-3200 BCE, and one hunter-gatherer dating to 1600-1950 CE from Sakhalin Island. Supplementary Data file 1 gives tabular individuls on the samples.

**Boisman culture^1-3^**: The sites of the Middle Neolithic Boisman culture are distributed over the Pacific coastal area of the southwestern Maritime Province of Russia and are represented archaeologically by settlements with shell mounds. The subsistence of the Boisman culture was based on hunting, fishing, and gathering terrestrial and marine resources. The archaeological assemblage of the Boisman culture and other similar cultures of the Middle Neolithic of the southern region of the Russian Far East as well as some Neolithic sites of Northeast China and the northeast of North Korea are part of a distinctive archaeological complex that differs from synchronous archaeological complexes of northeastern China and the Chulmun culture of the Korean Peninsula. The radiocarbon dates of this culture range from 4900-2500 BCE, and the direct radiocarbon dates on bone obtained for this study ranged from 5000-3600 BCE. According to some scholars, anthropometric features of skulls from Boisman­2 are particularly reminiscent of Arctic populations, such as Chukchi. Morphological similarity has also been documented between the burials at Boisman-2 and two present-day populations in the Lower Amur River basin, the Nivh and Ulchi. Some scholars have suggested that the Boisman-2 remains are “the earliest known representatives” of a much broader “Pacific” branch of East Eurasian populations comprising coastal groups from Chukotka to Taiwan^4^. We successfully obtained ancient DNA data from 18 individuals from the Boisman-2 site.

**Yankovsky culture^5^**: Early Iron Age people of the Yankovsky culture (850-350 BCE) inhabited mainly the coastal and subcontinental zones from the northern Korea peninsula to the Eastern Primorye coast. The subsistence of the Yankovsky culture was based on hunting, millet cultivation and widespread exploitation of marine resources. Millet cultivation played some role in subsistence, but not at all sites. We successfully obtained ancient DNA from one individual assigned to the Yankovsky culture at Pospelovo-1, Burial 1, Primorsky Krai, Russia.

**Heishui Mohe^6,7^**: Historical accounts indicate that in the 7th-10th centuries to the period of the Sui and Tang dynasties on the territory from the lower reaches of Sungari, Usury and the adjacent Amur river valley, there were 16 tribes of the Heishui Mohe, divided into northern and southern Heishui. The latter occupied the northern and northeastern parts of the present-day Maritime Province of Russia. After the decline of Bohai (10th century) they are known in the Chinese historical records as the Eastern Jurchen. They have been hypothesized to be ancestral to the Jurchen, present-day Manchu and other Tungusic peoples. We successfully obtained ancient DNA from two individuals belonging to the Heishui Mohe culture from Heishui Mohe Cemetery, Burial 2 and 6, Primorsky Krai, Russia, and one Mohe Bohai individual from Primorsky Krai, Chernyatino 5, Mohe-Bohai Graveyard.

**Hunter-gatherers of Japan and Sakhalin**: We obtained ancient DNA from 7 Jomon individuals from Funadomari site and Rokutsu site at about 1500-1000 BCE and 3500-3200 BCE, and one Sakhalin island hunter-gatherer dating to 1600-1950 CE.

The Funadomari site is a shell-midden site located in the northern part of Rebun Island, Hokkaido Prefecture northernmost of the Japanese archipelago. The site is elevated approximately 5 metres above sea level on a sand dune along the Funadomari Gulf. In 1998, Nishimoto led a team that uncovered 28 primary inhumation burials in good preservation in a flexed position and associated with numerous artefacts including necklaces, bracelets and anklets made of marine shell (*Mercenaria stimpsoni*)^8^. The typology of the cord marking decorations on the unearthed pottery is related to the "Kasori B" type found on the island of Honshu, Chiba Prefecture Japan, suggesting that these individuals were buried during the early to middle phase of the Late Jomon Period (ca. 1800-1500 BCE)^8^. Despite their geographical remoteness, the Funadomari Jomon crania show morphological resemblances to other known Jomon crania from Honshu Island^9^ belonging to the first layer of *Homo sapiens* that dispersed in the pre-farming period^10^. The direct 14C date for individual 23 gave a date of 4025±20 (1σ) calBP, 3846-3644 calBP (68.2%), 3960-3550 calBP (95.4%) (Paleo Labo Co., Ltd.)^11,12^. Another 14C dating of charcoal from the hearth resulted in a range between the top layer of 3635±65 calBP and lowest layer of 3850±55 cal BP^8^.

The Jomon shell-midden site of “Rokutsu” is located in Ohkanza town in Chiba City, Chiba Prefecture adjacent to metropolitan Tokyo. The large-scale rescue excavation took place in 2006 by the archaeological center of Chiba city, and was led by Hideyo Tanaka and Wataru Nagahara (report unpublished). The analysis uncovered the remains of seven inhumation burials. The individuals in burial No 1 (adult female), 2 (adult male), 5 (adult female), 6 (adult male), and 7 (adult male) were laid down on the side with limbs extended, while individuals 3 and 4 (female adult) were buried with flexed joints at the elbow, hip and knee. The local typology of Jomon pottery assigned to these burials was Late Jomon in the range 2000-1500 BCE (pottery typology, Shoumyoji, Horinouchi and Kasori B types).

The Nevelsk 2 site is located on the southwestern coast of Sakhalin Island in the southern part of the Tatar Strait. Currently, it is almost completely destroyed by modern buildings. In 1979, road construction work uncovered a cultural layer from the Okhotsk culture (7-10^th^ centuries CE) as well as three burials of a later time introduced into the Okhotsk layer. All three burials of the Nevelsk 2 site were accompanied by funerary objects of styles that were common for Sakhalin hunter-gatherers from the 17^th^ century to the mid-20^th^ century CE. Burial # 2 from which we obtained DNA was oriented with his head to the north, and laid on his right side (facing west) with bent legs and arm bones extended in front of the chest. The left temporal part of the skull was damaged. Funerary objects associated with this burial included 13 blue glass beads (found in the region of the cervical vertebrae; they made up a small necklace), the remains of an iron knife with a wooden handle, and a bone tip. Near the leg bones there were remains of wooden red lacquerware and fragments of an earthenware (faience) saucer. These items of Japanese production were used by Sakhalin hunter-gatherers in everyday life, and are the most frequent finds usually found during the excavation of such burials. The date of Burial #2 is likely to be from the 19th to early 20th centuries CE based on cultural context.

**Wuzhuangguoliang Neolithic of China**: The traditional archaeological culture classification in China identifies six large regional systems in the Neolithic: North, Central Plains, East, Southeast, Southwest and South^13^. Our study focused on the site of Wuzhuangguoliang (WZGL), in the contact zone between the Central Plains and the North cultural regions. The WZGL site is located at Xiaojie village, Jingbian County, Shaanxi Province, adjacent to Inner Mongolia in the north and Yan'an in the south. The excavators divided the site into three areas, A, B and C, with a total area of about 1,740 m^2^, exposing 21 houses, 88 ash pits and 2 pottery kilns. The exposed human bones were all from the ash pits. A total of 20 human bones for DNA analysis were collected from the ash pits distributed in the A and B areas. The site was comprehensively excavated by the Shaanxi Provincial Institute of Archaeology in 1996 and 2001. Radiocarbon dating, based primarily on zooarchaeological remains in ash pits from this site, suggests dates of 3317-2921 cal BCE (4422±29 BP, XA-8399). Two phases can be discerned in this site based on architecture, ash pits, pottery kilns, and archaeological stratigraphy and typology, and it was suggested that the two phases might belong to the Haishengbulang culture and Ashan culture respectively, ranging from the late Yangshao period to the early Longshan period^14^. The distribution of Phase One is located in the east of the site coded as area A while Phase Two is located in the southern parts of the sites coded as areas B and C. The archaeological remains included slipped and corded pottery in various forms including jars, basins, jugs, vats, bottles and bowls, with fine clay and sand-tempered in red, gray and reddish-brown. It also included polished stone axes and elaborate houses with gardens and fireplaces. The pointed-bottomed bottles with cup-shaped mouths, which are representative of the Yangshao culture^15^, suggest a connection with the Central Plains cultural region. Both hunting and gatherering and raising of domesticated animals was practiced at WZGL, with some ancient animal bones being from domestic animals such as *canis familiaris* and *sus domestica*, and others being from wild animals such as *lepus capensis*, *procapra gutturosa*, *equus sp*. and nearly complete felid skeletons identified as *coun alpinus* and *Felis silvestris* in wild^16^. In the Neolithic WZGL had a steppe ecology, but today it is in a desert environment and the vegetation is accordingly very different^17^. The domestication practices of *sus domestica* are similar to those of the Guanzhong Area which belongs to the Central Plains cultural region, further highlighting a connection between the areas^18^.

Physical anthropological and molecular anthropological analyses have been carried out on ancient human bones from three ash pits (AH1, BH23 and BH35). Previous ancient mtDNA study of human bones of this site suggested that ancient WZGL people shared lineages with East Eurasians, identifying haplogroups A, B, D, R9a, N9a, F1a and Z^19^. Morphological observation and measurements highlight the close relationship of the WZGL individuals to the present-day Liuwan people (eastern Qinghai province), Yuchisi people (northern Anhui province) and Xixiahou people (middle Shandong province). The WZGL site provides comprehensively verified evidence of the use of ash pit burial in northern Shaanxi and contributes to our understanding of living habits and ideology, as well as the physical characteristics of ancient inhabitants in that period. Radiocarbon dating of two of the ancient bone specimens from the WZGL site that we analyzed for this study (AHI-7 and AHI-17) was undertaken by accelerator mass spectroscopy (AMS). The radiocarbon dates were calibrated using CALIB Rev 7.0.2 software^20^ and the intcal13 radiocarbon age calibration datasets^21^.

**Hanben site:** The Blihun Hanben site is located in the south of Yilan County in eastern Taiwan. It was discovered in 2012 during the construction of the Suhua Highway Improvement. The Hanben site is situated on the coastline near the Pacific Ocean, only about 200 m from the sea at current levels and about 19 m above sea level. The site is 1500 to 2000 m from north to south and 150 to 200 m from west to east. There are at least two prehistoric cultural layers at this site. The first (L4) has been dated to 350-750 CE, thus being part of the Taiwanese Iron Age; the second (L6) to 50 BCE to 350 CE, within the transition from Neolithic to Iron Age in Taiwan. Based on the ceramics and burial practices, the remains within L4 could be considered part of the Shisanhang culture. One of the key characteristics of the Hanben site is that all of the burials in L4 are sideways-flexed, similar to other Shisanhang sites. The other similar characteristic is the iron metallurgy at BHB, another important feature of Shisanhang culture. The remains from L6 cannot be assigned to any other existing prehistoric culture in eastern Taiwan. However, the flexed in a seated position and sideways flexed burial practices in L6 indicate relationships to other prehistoric cultures in eastern Taiwan.

**Green Island, Gongguan site:** The site is located on the north coast of Green Island, which is a first-level coastal terrace that is curved in a concave shape. The square coordinates are E299580m×N2508550m and the altitude is about 5-10 meters. The Gongguan site was dated to 260-650 CE using charcoal from the cultural layer.

**Mongolian sites**

In what follows we describe the archaeological context for the newly reported Mongolian individuals in our study. We also justify the material culture classification that we use to group the individuals prior to genetic analysis.

**Neolithic of Eastern Mongolia:** Burials belonging to the Neolithic of Eastern Mongolia (Dornog aimag) probably belong to the so-called Tamsagbulag culture of the 6^th^ to 5^th^ millennia BCE. To date, numerous sites of this culture have been investigated and permanent settlements with durable semi-subterranean houses were excavated, with the largest house recorded as 120 m^2^ in area. Finds include a large number of knife-like flakes and tools made from the flakes (including cutters, knives, knife-blades, awls and scrapers), as well composite knives and daggers, arrowheads, polished tools, bone tools, bone and stone ornaments, and fragments of pottery. People of the Tamsagbulag culture engaged in primitive hoe agriculture combined with food-gathering, fishing, and hunting; whether or not stockbreeding was practiced is questionable. Some features of the Tamsagbulag culture have analogies in the Transbaikalia region^22^.

**Neolithic of Northern Mongolia:** Excavations in the Egiin gol river basin (Bulgan aimag) revealed a group of burials under stone heaps. Burial goods include composite knives, inset flakes, arrowheads, and stone beads. Burial custom and burial goods have analogies with Cis-Baikalian Neolithic cultures^23,24^.

**Afanasievo culture^25,26^:** People of this culture are believed to have migrated from the Volga-Ural steppe region to the area of the Russian Altai in the late 4^th^ millennium BCE and then to have spread to Middle Yenisei basin, Eastern Kazakhstan, Northern Xinjiang, and Central Mongolia between 3000-2600 BCE. The basis of the economy of this culture was sheep and cattle breeding, and small seasonal settlements have been identified. The Afanasievo is assigned to the Chalcolithic Period based on the lack of metal artifacts made from bronze, though artifacts made from copper, gold and meteorite iron have been found. Peoples of the Afanasievo culture used egg-shaped and ball-shaped pottery and so-called “censers” (incense burners) with different stamp ornamentation, similar to the pottery of the earliest Yamnaya culture (or pre-Yamnaya period) of Eastern Europe.

**Chemurchek (Qiemuerqieke) phenomenon:** The Chemurchek phenomenon is named for the river 切木尔切克Qiemuerqieke in the Altai county of Xinjiang^27,28^. A complex of cultural features resembling those of contemporary peoples in Western Europe appeared in the slopes of Mongolian Altai (Northern Xinjiang and Western Mongolia) not later than 2700-2600 BCE. The architecture of the burial constructions, specifically megalithic chambers constructed from huge stone slabs with collective burials surrounded by multiple stone and soil cairns overlapping one another, the form and ornamentation of vessels, the stylistics of stone statue-menhirs with protruding facial contours and circular protruding eyes, paintings on slabs inside burial chambers (rhombs and chevrons inscribed into each other, parallel multi-triangle festoons, sloping nets, nets with cells filled with roundish spots, meander-shaped and volute-shaped curves), and other religious objects and images have strong analogies to cultural materials of the Middle-Late Neolithic of Western Europe. Only burial and ritual complexes of this entity were excavated (no settlements were identified). Finds included tin bronze and lead artifacts and domesticated animal bones. The Chemurchek people had close contact with peoples of the late Afanasievo culture who spread into Xinjiang, and later influenced the Elunino culture of Steppe Altai, the Karakol culture of Mountain Altai, and the Okunevo culture of Middle Yenisei basin.

**Early Bronze Age of Ulgii:** This group is represented by five sites dated to the latter half of the 3rd millennium BCE excavated near the northwest edge of Mongolia in the high-altitude Altai mountains (Bayan-Ulgii aimag). Burials were placed in the middle of rectangular fences with stone stelae with ritual purposes. These fences with stone stelae were also a characteristic attribute of Chemurchek burial places in the neighboring Chinese Altai. Pottery vessels, bone tools, bone arrowheads, small stone balls with analogs in Early Bronze Age cultures of Eastern Kazakhstan, Russian Altai and Khakassia have been unearthed in these graves^28^.

**Munkhkhairkhan culture:** This culture dates to ~1800-1400 BCE and ranges across the territories of Western and Northern Mongolia, Tuva, and the southern part of the Krasnoyarsk region in Russia. It is presently characterized only by funerary and ritual complexes (no settlements have been identified). Crouched burials were arranged in small pits filled by stones inside flat round or rectangular stone platforms. Ordinary barrows are 3-5 m in diameter; however, in special cases, platforms were built measuring 30-40 m in diameter. Ritual structures include stone circles with stelae having astronomical significance. No pottery has been found. Burial goods include tin bronze knives and awls, tin bronze two-trumpet shaped rings, bone spoons, bone arrowheads, bone, and shell and stone ornaments. Knives and rings have analogs both in Western Siberia and in Chinese territories occupied by the late Qijia, Lower Xiajiadian, Siba, and Erlitou cultures. As such, it has been suggested that cultural innovation from Seima-Turbino and Andronovo complexes was introduced into the Central Plain via the Munkh-Khairkhan culture. The origin of the Munkh-Khairkhan culture may be in the forest zone of Eastern Siberia in the upper Lena river basin near the Cis-Baikalian region. Bone spoons, arrowheads, and mother-of-pearl discs attributed to Cis-Baikalian Neolithic and Early Bronze Age cultures have similarities with Munkh-Khairkhan, as well as funerary customs such as the filling of burial pits with stones^29,30^.

**Ulaanzukh type:** The Ulaanzukh type is part of the “figured graves” that spread over Southeast Mongolia ~1450-1250 BCE. This group was characterized by burials under square platforms edged by masonry walls. Other groups of such graves have concave or convex sides (considered part of the Tevsh culture)^28^ and are spread primarily in South Mongolia, but also partly in Central Mongolia as well as in the north of Inner Mongolia. All these burials were characterized by skeletons in prone positions with the head to the east. Burials are arranged in pits dug into the ground and filled with soil. Artifactual finds in Ulaanzukh type graves include pottery tripods, cornelian and limestone beads, gold ornaments, and stone polished tools. The prone position of buried bodies and findings of metal ornaments similar to those of northern Chinese Late Bronze Age cultures suggest contact with northern Chinese minorities during the late Shang period. It has been proposed that all graves with masonry-walled platforms and skeletons buried in the prone position be considered part of the Ulaanzukh-Tvesh culture^3^^1^.

**Late Bronze Age of Center and West Mongolia and “Mongun-Taiga type”:** Explorations and excavations in Center and West Mongolia and in Tuva conducted since the 19^th^ century have revealed a large number of stone barrows of different shapes. The majority of these barrows include human burials but lack burial goods. The skeletons are arranged in stretched or in slightly crouched positions on their side or on their back in narrow earth pits or in stone cists on the ancient surface, covered by stones or organic materials^31,32^. Cairns of these barrows have circular or quadrangular shapes and may be surrounded by circular or trapezoidal fences with stelae or small stone heaps in the corners (so-called Khereksurs in Mongolian). Near to these kurgans, ritual burials of horse skins skulls were arranged within stone rings and under stone pavements. All of these barrows can be dated to the Late Bronze Age (~ 1400-1000 BCE). Mongolian and foreigner scholars have attempted to isolate particular types of these burial structures and designate them as belonging to specific “cultures” (e.g., “culture of Khereksurs and Deer Stones”, “Sagsai culture”); however, there is not strong evidence for these cultural designations. Russian archaeologist Alexander Grach who carried out excavations in Tuva in the 1950-1970s years proposes the use of the name “Mongun-Taiga type” as a designation for burials with the body in a stretched position on the side with the head to the west. Excavations and investigations in the Mongolian Altai by A. Kovalev and D. Erdenebaatar show that such burials were not surrounded by stone fences and were not accompanied by horse sacrifices. Radiocarbon dates of such barrows places them into a narrow span ranging between 1400 and 1200 BCE. Therefore, we considered the LBA Mongun-Taiga group independently, a classification that found support in our genetic analysis as the three individuals we analyzed from the Mongun-Taiga context were genetically distinctive from the main group of LBA Center-West individuals (Figure 3). Explorations of this region suggest that Late Bronze Age people from Central-West Mongolia and Tuva practiced sheep breeding and did not have permanent settlements. However, based on the number of burial mounds, it is likely that the population of this area of Mongolia and Tuva was very large. Anthropological typing identifies these people as ‘Caucasian’ with ‘Mongoloid’ admixture increasing from west to east, a suggestion that is qualitatively supported from a genetic point of view as ancestry related to Sintashta and Andronovo people is more evident in the west than in the east. Russian anthropologist I. Gohman identified similarities between these populations and Afanasievo people of the Russian Altai and Minusinsk Basin.

**Slab Grave culture:** The people of this culture spread across East and Central Mongolia (up to the Gobi region) beginning in the 10^th^ to 8^th^ centuries BCE, and lived in this region at least until the 5th century BCE. It is likely that the center of this culture was in Transbaikalia, where rich burials of the so-called Dvortsovo type and different types of Slab graves have been discovered. Anthropological typing of the individuals of this culture suggests they are ‘Mongoloid.’ The economy of this culture was based on the breeding of sheep, goats, cows, and horses. Graves of this culture are characterized by rectangular stone fences oriented west to east, inside of which burials were positioned stretched out on their back with their heads toward the east. In the corners of the stone fences, vertical stone slabs were often erected. The space inside the fence was littered with stones and soil to make a flat platform. Ceramic vessels were broken and fragments were placed in the corners of the platforms. Burial goods included bronze weapons, horse harnesses, and ornaments and tools of so-called Scythian types as well of types used by peoples of Northeast China (such as the Upper Xiajiadian culture and Yuhuangmiao culture, dating to the first half of the 1^st^ millennium BCE). It is possible that there was both demographic and cultural connections between the populations of Transbaikalia, East Mongolia, East Inner Mongolia, and North Hebei during this period.

**Sagly culture:** This is one of the cultures of the so-called Scythian-Siberian type^33^. This culture spread in Tuva during the 6^th^–3^rd^ centuries BCE. One cemetery of this culture (Ulaangom cemetery) was excavated in Mongolia near the Tuvinian border, and Mongolian scholars designated these graves as belonging to the “Chandman culture.” At present, only one cemetery in Mongolia is still attributed to this culture, while more than 50 cemeteries of this culture were excavated from the beginning of the 20^th^ century in Tuva. Burials of this culture were arranged in large timber chambers (approximately 4x4 m to 5x5 m and 3-4 m deep). Four to five people were buried in each chamber, likely one after the other. Burial goods included bronze and iron weapons and horse harnesses of Scythian style; bronze, gold and bone ornaments in “Animal style”; and other items attributed to the neighboring Pazyryk culture in the Altai Mountains. Ceramic vessels were determined to have similarities to Pazyryk vessels as well. People of this culture are anthropologically typed as mixed between ‘Caucasoid’ and ‘Mongolid’ with the ‘Mongolid’ component increasing in presence in the later part of this culture. It is possible that an East Asian component represents matrimonial connections with Mongoloid peoples of North and North-West China during the Zhanguo period. Like the Pazyryk culture, the Sagly culture had strong connections with the regions of Ordos and Western Tianshan. Genetically, we found that Sagly culture individuals and Pazyryk culture individuals were similar in their deep ancestry proportions, further supporting their connection (Figure 3).

**Xiongnu (Hunnu) burials and Xianbei.** The first evidence of Xiongnu nomadic cattle-breeding tribes in the Chinese historical sources appeared in 3rd century BCE. In this time, the Xiongnu were attested to live to the North of the bend of Huanghe river (Hetao) of modern Shanxi and Hebei provinces^34^. Archaeologically, the first evidence of the Xiongnu (Hunnu) dates to the 2nd century BCE, when this cultural tradition was associated with a vast empire, and rapid material culture evolution. Xiongnu style graves in this period spread over Mongolia and over the Buryatia and Tuva regions in Russia. In Xiongnu style burials, physical anthropological evidence shows that the “Mongoloid” component decreases from the West to East suggesting a mixed ancestry. The origin of Xiongnu has been hypothesized to have been in Eastern Mongolia and Manchuria. Ordinary graves of Xiongnu are arranged as deep rectangular pits with wooden coffins with buried people lying on the back with the head to the north. In the northern part of the pit, heads of goats and other livestock animals were often laid. Elite graves of the Xiongnu were arranged as imitation of elite Han empire tombs in huge pits with a ramp from the south. Traces of shamanistic rituals and human sacrifices are evident in elite burials. In the north of Mongolia and in Buryatia, Xiongnu settlements are associated with a high level of craftsmanship. Following the 2nd century BCE, people practicing Xiongnu culture began to use powerful weapons, most notably a long compound bow with arrows tipped with iron pints; they also used Chinese iron long swords and iron horse harnesses. According to Chinese narrative sources, after Chinese offensives and the Xianbei expansion in the 2nd century CE, north Xiongnu leaders migrated out of Mongolia and local people accepted Xianbei customs. The earliest Xianbei cemeteries are thought to be from eastern Inner Mongolia. The 3-4 century CE tombs associated with this material culture in Mongolia in fact demonstrated a broad diversity of burial customs and goods so we can call them Xianbei only conditionally.

**Mongol graves.** Starting from the 10th century, a characteristic grave type associated with Mongols can be distinguished. These graves looks like narrow rectangular pits with burials on the back and head to the north, with a lamb shoulder blade near the head. Later, in the period of Mongol empire, buried people were laid in a special room dug into the side of burial pit. After Genghis Khan united the disparate Mongol tribes, the anthropological characteristics became somewhat different. The origin of the Mongols has been hypothesized to be associated with eastern Mongolia but Mongol ancestors settled in Central and Western Mongolia before the Turkic and Uigur expansions. Mongol scholars consider the Xiongnu as ancestors of the Mongols.

**Supplementary Information section 2: Overview of genetic substructure**

There are more than 200 languages in East Asia that are distributed into seven families: Sino-Tibetan, Austroasiatic, Austronesian, Tai-Kadai, Hmong-Mien, Indo-European and Altaic (including Mongolic, Turkic, and Tungusic)^1^. However, genetic structure across human populations in East Asia remains enigmatic due to limited sampling compared to other world regions, particularly in the Tibetan Plateau and in southwest China^2^. We report a fine-scale survey of the genetic structure of East Eurasians based on genome-wide variation from East Asians, Southeast Asians, and Siberians, with a specific focus on East Asia. We use a variety of methods for studying genome-wide variation data to identify qualitative patterns in the dataset. In many instances, we find that genetic clusters correlate strongly to language groupings. However, there are also exceptions where language group does not correspond to genetic patterns, and we highlight some of these patterns.

**Genetic differentiation patterns (*F_st_*)**

We computed *F_st_* between all pairs of the present-day populations using smartpca (version: 13050^3^, part of the EIGENSOFT package^4,5^). We found that the inbreeding corrected and uncorrected *F_st_* were nearly identical (within ~0.001), and in this study, we always analyzed uncorrected *F_st_*. Han Chinese are relatively homogeneous with the smallest pairwise *F_st_* ranging from 0 to 0.006. The largest pairwise *F_st_* values within East Asia are found between Tungusic-speakers (e.g. Nivh) and Austronesian-speakers (e.g. Atayal), where genetic differentiation reaches trans-continental levels of ~0.1 (Figure S1).

We observe that neighbor-joining *F_st_* trees are strongly correlated to geography or language groupings. Qualitatively, we observe five main clusters, which correlate strongly to Altaic-speakers, Tibeto-Burman-speakers, Han Chinese, Southeast Asians (largely Tai-Kadai, Austronesian and Austroasiatic speakers), and Oceanians.

We also observe sub-clusters within these main groupings, including a sub-cluster of populations centered on the Amur River Basin (Nivh and the Tungusic-speaking Ulchi, Nanai, and Negidal), northeastern Siberians (Chukchi, Itelmen, Koryak and Eskimo), Ainu-Japanese, Nepalese groups, and Papuan-Australian. We caution that any patterns in neighbor-joining *F_st_* trees should be viewed with caution. For example, Papuans and Australians are known highly divergent lineages, but they cluster with other Oceanians. It is likely that their derived position in the tree reflects admixture from Papuan-related (Melanesian) populations into nearby groups.

There are also some notable cases in which the position in the tree does not correlate perfectly to linguistic clusters or geography:

*Populations with genetic affinities with West Eurasians*. Three Turkic-speaking populations in northwest China—Uygur, Kazakh-China, and Kyrgyz-China—cluster with French and Russian rather with other Turkic groups in China. The Nepalese Bahun also cluster with West Eurasians.

*Austroasiatic speakers*. Some Vietnamese including the Kinh cluster with mainland Tai-Kadai speakers rather than with other Austroasiatic speakers. Nicobarese and Cambodians form a separate cluster among Island Southeast Asians and Oceanians together with Thai and Borneo.

*The Lahu*. This Lolo-Burmese speaking population is genetically close to Tai-Kadai speakers.

*The Gannan Tibetans*. This northern Tibetan population clusters with Turkic and Mongolic populations.

*Hmong-Mien speakers*. These populations tend to be genetically intermediate between Tai-Kadai and Han.

**Principal Component Analysis**

We used smartpca (version: 13050), part of the EIGENSOFT package^4,5^, to carry out Principal Component Analysis (PCA). We performed PCA on present-day populations from East Asia, Siberia, Europe, and Southeast Asia and then projected the ancient samples using the lsqproject: YES option, which accounts for samples with substantial missing data. We did not perform any outlier removal iterations (numoutlieriter: 0). We set all other options to the default. We assessed statistical significance with a Tracy-Widom test using the twstats program of EIGENSOFT. All the first six principal components that we discuss and plot in what follows were highly statistically significant (P<10^-12^).

In the first two components, there are two obvious clines from East Asians to West Eurasians, and from East Asians to Native Americans. Nepalese groups and populations in northwest China and Siberia (including Turkic speakers, Mongolic speakers and Tungusic speakers) are shifted towards the West Eurasians, while Northeast Asians are intermediate between East Asians and Oceanians. We observe that populations speaking the same language in East Asia tend to cluster, albeit with exceptions such as Turkic speakers, who are relatively scattered in the plot which is plausibly due to complex admixture with surrounding populations^6^ (Figure 2A).

The ancient individuals from Devil’s Gate, Boisman, and Yankovsky project close to present-day Tungusic speakers in the Amur River Basin, documenting a continuous presence of this ancestry profile in the Amur River Basin stretching back at least eight thousand years. It shows that the ancient Heishui_Mohe individual I1209 and present-day people in northwest China and Siberia deviate in the direction of West Eurasians. Nepalese populations share a similar statistical signal. Note that the other Heishui_Mohe sample I3358 plots together with Tibetan groups and populations in northern China. In any case, because of the inconsistency of the evidence for West Eurasian genetic affinity in the two Heishui_Mohe individuals, we do not make any claim in this study for West Eurasian admixture in this ancient population but discuss the two samples individually in some analyses.

The Mongolian ancient individuals project on the cline from East Asians to West Eurasians. The indiviudals with minimum proportion of West Eurasian-related admixture (< 10%), such as Mongolia_N_East, Mongolia_N_North, Mongolia_LBA_2_Ulaanzukh, Mongolia_EIA_1_SlabGrave, and Mongolia_EIA_8 project close to the Amur River Basin populations. The individuals samples with about 20% West Eurasian-related ancestry from cultures such as Mongolia_LBA_4_CenterWest, Mongolia_EBA_1_Ulgii, Mongolia_MBA_1_Munkhkhairkhan, and Mongolia_LBA_6_Khovsgol project closely with published Neolithic and Early Bronze Age individuals around Lake Baikal. The samples with about 40%-80% of West Eurasian related ancestry from Mongolia_EBA_2_Chemurchek, Mongolia_EIA_6_Pazyryk, Mongolia_LBA_5_CenterWest, Mongolia_EIA_4_Sagly, Mongolia_MBA_2_Munkhkhairkhan project at an intermediate position in the East-West cline. The individuals from Mongolia_Chalcolithic_1_Afanasievo cluster together with West Eurasians.

PC2 reveals a “south-north cline” within East Asia defined by the Southeast Asian Cluster (Tai-Kadai speakers in southern China and Austronesian speakers in Taiwan Island) at one extreme and Eskimo-Inupiat speakers in northern Siberia and Native Americans on the other. Ancient Taiwan samples from Hanben and Gongguan sites project closely together within the Southeast Asian Cluster. The ancient Devil’s Gate, Boisman and the ancient Yankovsky project close to present-day populations in the Amur River Basin, at an intermediate position in the south-north cline. The Neolithic Wuzhuangguoliang samples from northern China project with northern Han Chinese and other populations in northern China. Tibeto-Burman speaking populations on the Tibetan Plateau show complex patterns. Tibetans in Tibet plot tightly between the Amur River cluster and Han Chinese, but slightly off of the south-north cline. Northern Tibetans deviate in the West Eurasian direction. In the Tibeto-Burman Corridor, Tibetans, Lolo-Burmese speakers (Yi, Naxi, and Lahu), and Qiang are scattered along the south-north cline.

We also projected ancient samples onto the PCA inferred using only East Asian populations after removing Uygur, Kazakh_China, Kyrgyz_China, Salar, Dongxiang, Bonan, and Yugur in northwest China from the dataset as they deviate towards West Eurasians in the East Asian-Siberian PCA (Figure 2A). The first two PCs correlate to the geographic map of East Asia and linguistic categories. There is a clear south-north cline from indigious populations in Taiwan and Nicobar Islands, to populations in the Amur River Basin and Tibetan Plateau. Han Chinese appear intermediate between the south and the north. We also observed genetic substructure in Han Chinese, with northern Han Chinese closer to ancient and present-day populations in the Amur River Basin, Wuzhuangguoliang and Tibetan Plateau, and southern Han Chinese deviating toward populations in Mainland Southeast Asia and southern China. Populations of the same region tend to group together, forming Taiwan Island, southern China, Han Chinese, Tibetan Plateau, and Amur Basin clusters. The ancient Devil’s Gate, Boisman and Yankovsky samples project within the Amur Basin cluster. There is substructure within Southeast Asian groups. Mainland Southeast Asians (Vietnamese, Cambodian, and Kinh) plot in the southern China cluster, while island individuals (Nicobarese) are outliers. Miao and She show closer affinity with Han Chinese. Populations in the Tibeto-Burman Corridor are intermediate between Han Chinese and Tibetans in Tibet. Our ancient Jomon and Sakhalin_HG samples project on the direction driven by Japanese and Amur River Basin populations, showing affinity with those populations. Mongolia ancient samples project into two separate clusters at an intermediate position between the Amur River Basin cluster and Tibetan Plateau cluster.

**ADMIXTURE analysis**

We carried out model-based clustering analysis using ADMIXTURE 1.23^7^, combining the present-day worldwide populations with our newly reported ancient individuals. We used PLINK v1.90^8^ to thin the dataset of 597,573 autosomal SNPS to remove SNPs in strong linkage disequilibrium, employing a window of 200 SNPs advanced by 25 SNPs and an *r^2^* threshold of 0.4 (with the flag: --indep-pairwise 200 25 0.4). A total of 317,286 SNPs remained for analysis after this procedure. We ran ADMIXTURE with default 5-fold cross-validation (--cv=5), varying the number of ancestral populations between K=2 and K=15 in 100 bootstraps with different random seeds. We used unsupervised ADMIXTURE, in which allele frequencies for non-admixed ancestral populations are unknown and are computed during the analysis. We used point estimation and terminated the block relaxation algorithm when the objective function delta < 0.0001. We chose the best run according to the highest log likelihood.

We used cross-validation to identify an “optimal” number of clusters. We observed the lowest CV errors for K=10. However, we report the full results for K=2-15, as we are interested in the genetic structure revealed by ADMIXTURE in successive models with increasing number of “ancestral populations” (increasing K) (Figure S2). In what follows we describe some qualitative patterns evident in this analysis.

K=2: African and West Eurasian populations separate from East Eurasians, Native Americans and Oceanians.

K=3: An African component separate from West Eurasian component. The ancient Mongolia samples, populations in northwest China, northern Tibetans in Qinghai and Gansu, and populations in Siberia and Nepal are assigned some ancestry from the West Eurasian component.

K=4: Native Americans form their own component, which is widely distributed in Siberian and East Asian populations, but nearly absent in populations of southern China and Taiwan Island. The Amur River cluster derives approximately half of its ancestry from the Native American component.

K=5: A component maximized in Oceanians appears, which is also distributed in ancient Southeast Asians and Jomon.

K=6: The Native American component separates into two parts, with one component maximized in southern Native Americans (Karitiana and Surui) and one maximized in Siberian populations such as Nganasan. Most East Asian populations have some of the Siberian component.

K=7: A component maximized in EHG appears, which also reaches high proportions in Steppe EMBA and MLBA populations and present-day Europeans and South Asians.

K=8: A component maximized in Early Neolithic Central Asians appears, and is represented to a lesser extent in ancient Steppe populations, present-day South Asians, Central Asians and Middle Easterner.

K=9: Tibetans on the Tibetan Plateau obtain their own component, which is also widely distributed in Han Chinese, Japanese, Koreans, and some other populations in Southeast Asia, southern China, northwest China and southern Siberia. Far Eastern Siberians, such as Itelmen, Koryak, Chukchi, and Eskimo, separate from other Siberian populations.

K=10: A component maximized in ancient Lapita and Tonga samples appears, which also reaches high proportions in ancient samples from Southeast Asia and Taiwan. Far Eastern Siberians, such as Itelmen, Koryak, Chukchi, and Eskimo, lose their own component.

K=11: Far Eastern Siberians, such as Itelmen, Koryak, Chukchi, and Eskimo, are assigned their own component again, which separate them from other Siberian populations.

K=12: A new component that is maximized in Neolithic and Early Bronze Age individuals around Lake Baikal appears, which is also distributed in ancient Steppe populations including our newly reported Mongolia ancient ones.

K=13: Ancient Jomon samples are assigned their own component, which also contributes substantially to the hunter-gatherer from Sakhalin island in Russia.

K=14: A new component that is maximized in Russia_Okunevo_BA appears, which is also distributed in ancient Steppe populations and Central Asians. Present-day Japanese show Jomon related component.

K=15: Surui obtains its own separate component.

**Genetic substructure in Tibetans**

As discussed in Principal Component Analysis and ADMIXTURE analysis, northern Tibetans in Gangcha, Gannan, and Xunhua have more West Eurasian related admixture than southern Tibetans in Tibet and the Tibeto-Burman Corridor, a phenomenon that we also highlight in Table S3 and Table S4. Northern Tibetans also share less genetic drift with other Tibetan populations in outgroup *f_3_*-statistics, which we hypothesize reflects their West Eurasian ancestry component (Figure S3). Tibetans in the Tibeto-Burman Corridor and Lolo-Burmese populations (Yi, Naxi, and Lahu) share more alleles with Southeast Asian Cluster in southern China and Taiwan than do Tibetans in Tibet itself (Figure S3). Based on these observations, we classify Tibetan populations into three groups based on outgroup *f_3_*- and *f_4_*-statistics: northern Tibetan, Tibet Tibetan, and Corridor Tibetan.

**Summary**

Qualitatively, East Eurasian genetic structure is strongly correlated with geography and linguistic categories, with important exceptions. We defined four genetic clusters within East Asia, correlating strongly to Amur River Basin, Han Chinese, Southeast Asian Cluster (populations in southern China, Taiwan Island, and mainland Southeast Asia), and Tibeto-Burman on the Tibetan Plateau. We observe two genetic subclusters within Mongolia: one falls closer to ancient individuals from the Amur Basin Cluster (hence termed ‘East’), and the second clusters towards ancient individuals of Afanasievo culture (henceforth called ‘West’), while a few individuals take intermediate positions between the two.

**Supplementary Information section 3: Admixture graph modeling**

In this section, we use *f_4_*-statistics and *qpGraph (version 6750)*,^1^ as implemented in ADMIXTOOLS,^2^ to investigate models of possible phylogenetic relationships for the representative East Asian populations. *qpGraph* assesses the fit of Admixture Graph models to data by comparing fitted and estimated *f_2_*-, *f_3_*-, and *f_4_*-statistics. We first use the 1240K dataset to explore the phylogeny and then restrict the analysis to transversion sites only to replicate the results. We use ancient Mongolia_Neolithic, Boisman, Chokhopani, Taiwan_Hanben, and Japan_Jomon to represent the distinct genetic clusters within East Asia. We use the following parameters in running the *qpGraph* program:

*outpop: NULL*

*blgsize: 0.05*

*lsqmode: NO*

*diag: 0.0001*

*hires: YES*

*initmix: 1000*

*precision: 0.0001*

*zthresh: 0*

*terse: NO*

*useallsnps: NO*

In what follows, we only accepting models as “fits” if all possible *f_2_-*, *f_3_-*, and *f_4_-* statistics have a |Z|-score of less than 3 between the observed and expected values, as determined by the Block Jackknife as implanted in *qpGraph*. We also inspect the likelihood scores output by *qpGraph* to further differentiated between models, and do not accept models that have internal branch lengths of zero as these may indicate error in the graph topology.

We constrained the models used MSMC^3^, following the procedures in Mallick et al^4^ to infer cross-coalescence rates and population sizes among Ami/Atayal, Tibetan, and Ulchi to represent ancient Taiwan_Hanben, Chokhopani, and Mongolia_Neolithic/Boisman respectively to provide insight into the relative timing of the population splits within East Asia. We used Mixe as an outgroup of present-day East Asians for comparison. The result suggests that the split between Tibetan and Ulchi is more recent on average than the splits of Tibetan and Ami/Atayal (Figure SP1), although it is important to realize that these splits represent averages of multiple lineage separations contributing to the groups being compared. Thus, it is possible that a subset of the lineages contributing jointly to Tibetan and Ami/Atayal may be more closely related than some of the lineages contributing jointly to Tibetan and Ulchi. We nevertheless used the relative split times to constrain the *qpGraph* models, prioritizing models that imply closer relatedness of Tibetans and Ulchi.

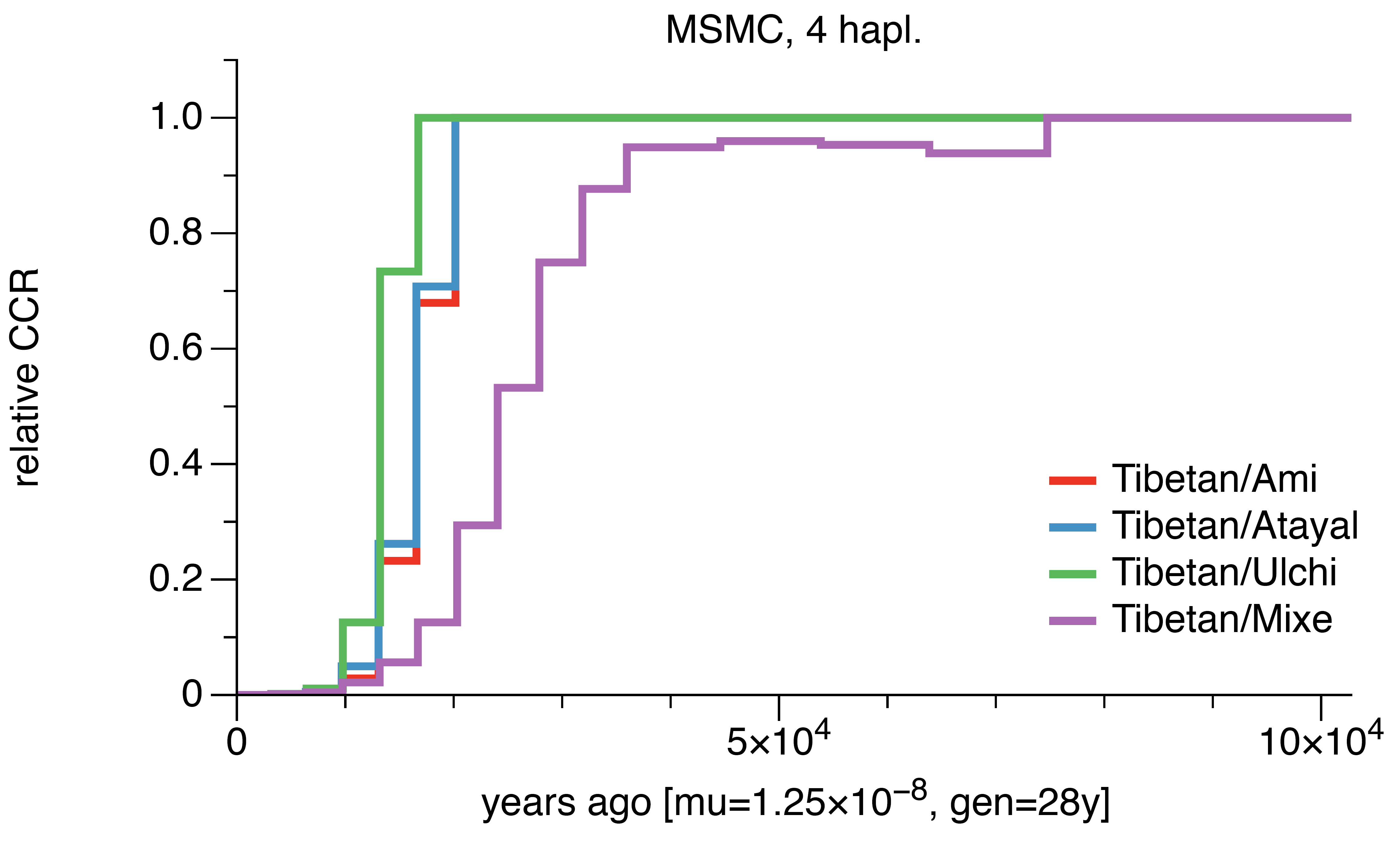

**Figure SP1: Cross-coalescence rates for selected population pairs. We ran MSMC for four pairs of populations: Tibetan-Ami, Tibetan-Atayal, Tibetan-Ulchi and Tibetan-Mixe. We used two individuals from each population in this analysis. The genome data for those samples are extracted from the Simons Genome Diversity Project^4^. The times are calculated based on the mutation rate and generation time specified on the x-axis.**

**1240K dataset:**

We first used the 1240K dataset. We started with a skeleton phylogenetic tree consisting of Denisovan, Mbuti, Loschbour, Tianyuan, and Onge with one admixture from Denisovan to Tianyuan (Figure SP2). The tree was an adequate fit with the likelihood score of 0.213 and the worst Z-score of *f*-statistic (Mbuti, Loschbour; Onge, Tianyuan) = 0.379.

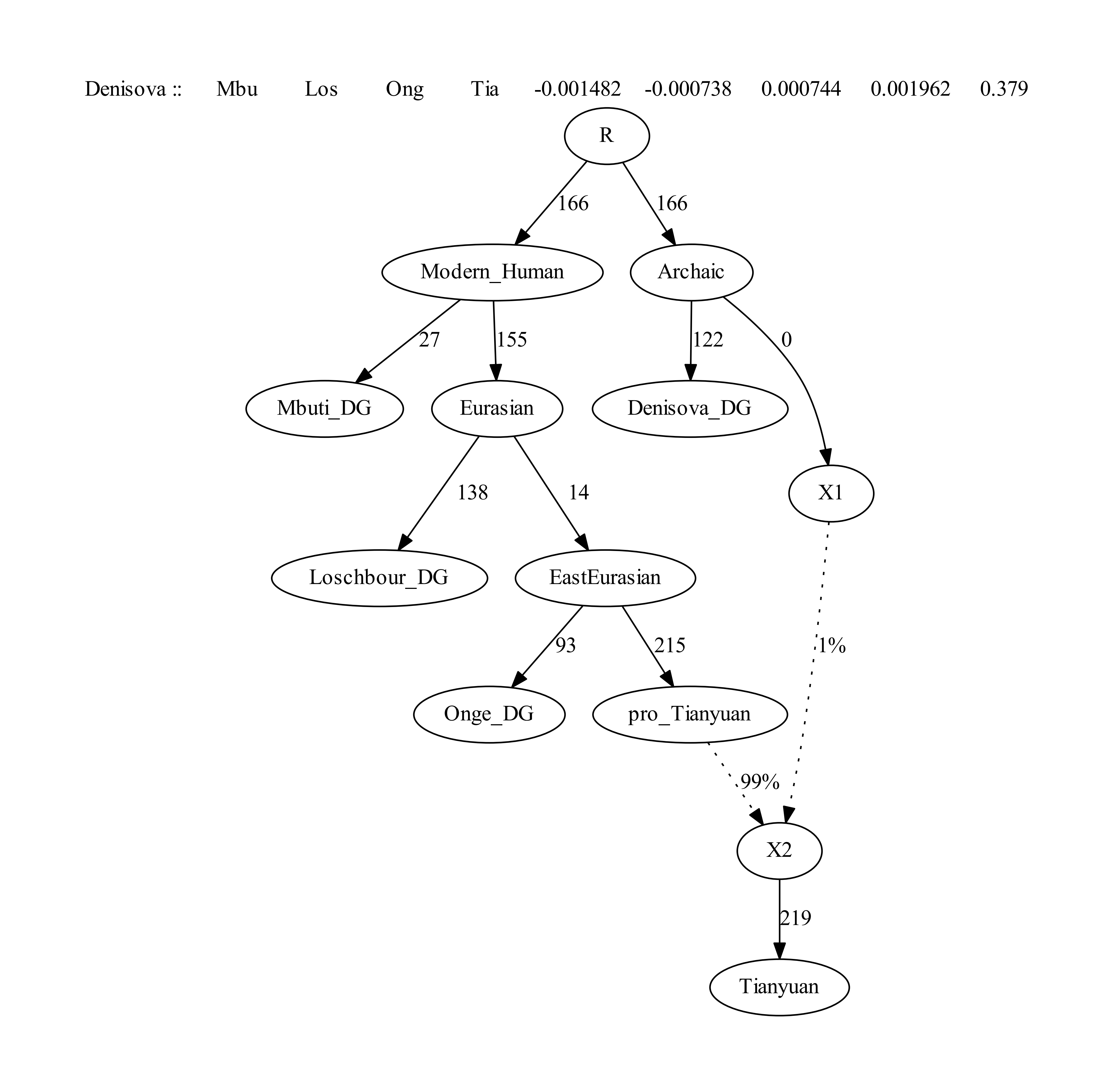

**Figure SP2: Skeleton Admixture Graph with one admixture using 652283 SNPs, which is a fit to the genetic data in the sense that there are no *f*-statistics more than |Z|>3 different between model and expectation. The drifts along edges in this figure and the following ones are multiplied by 1000.**

We then added Mongolia_Neolithic onto all the possible edges of the graph and found a fitted model with no additional admixture event of assigning Mongolia_Neolithic as a sister branch of Tianyuan (Figure SP3).

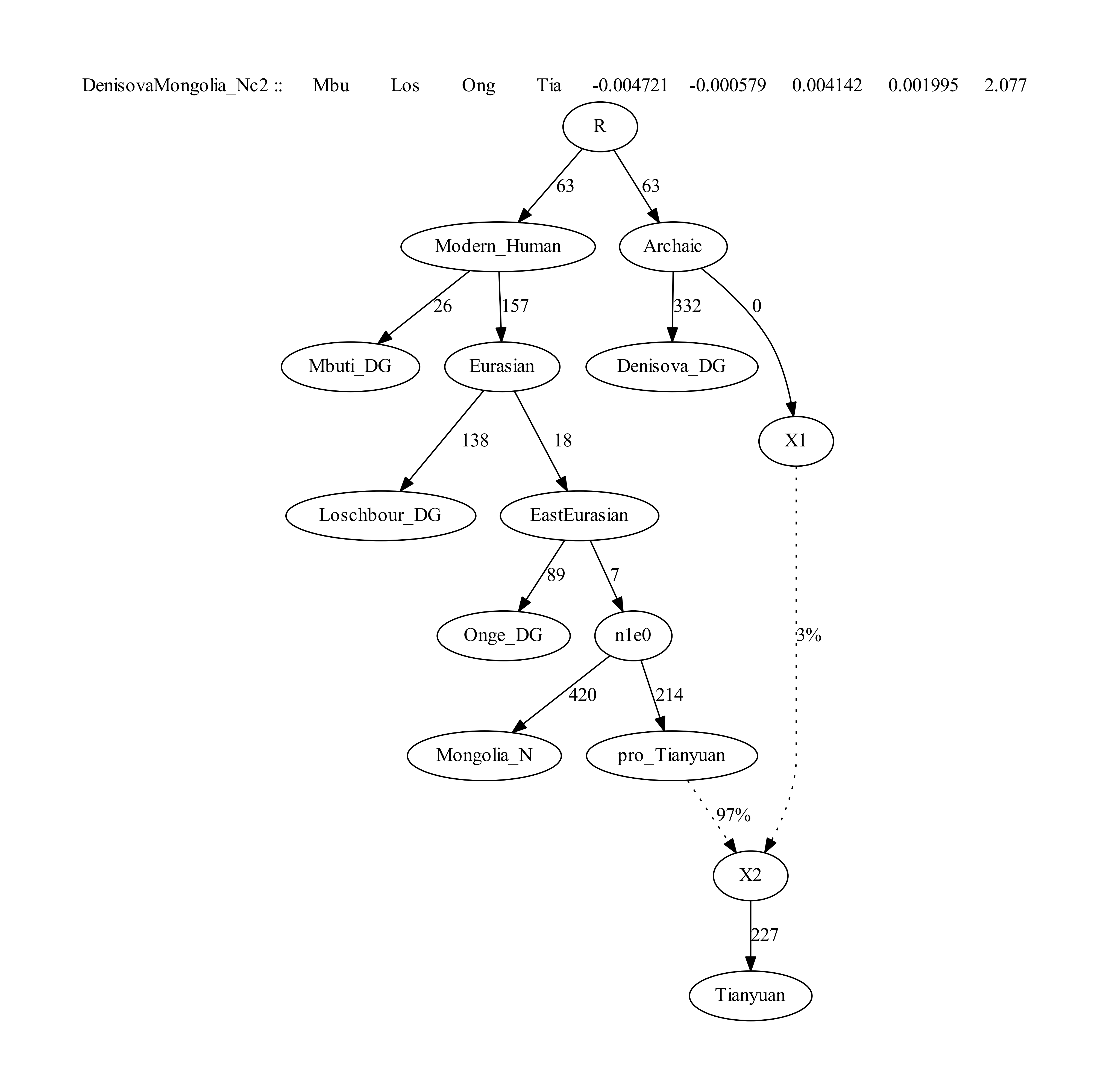

**Figure SP3: The best fitting tree of adding Mongolia_Neolithic without admixture was an adequate fit as in the sense that there are no *f*-statistics more than |Z|>3 different between model and expectation. The likelihood score is 9.206 and the worst Z-score of *f*-statistic (Mbuti, Loschbour; Onge, Tianyuan) = 2.077. The number of SNPs used is 561411.**

We then added Chokhopani onto all the possible edges of the graph. We were not able to fit Chokhopani into the graph without invoking an additional admixture event. We obtained only one fitted graph for modeling Chokhopani as admixed as shown below in the Figure SP4, suggesting Chokhopani deriving 86% of the ancestry from Mongolia_Neolithic related lineage and 14% from the Onge related lineage.

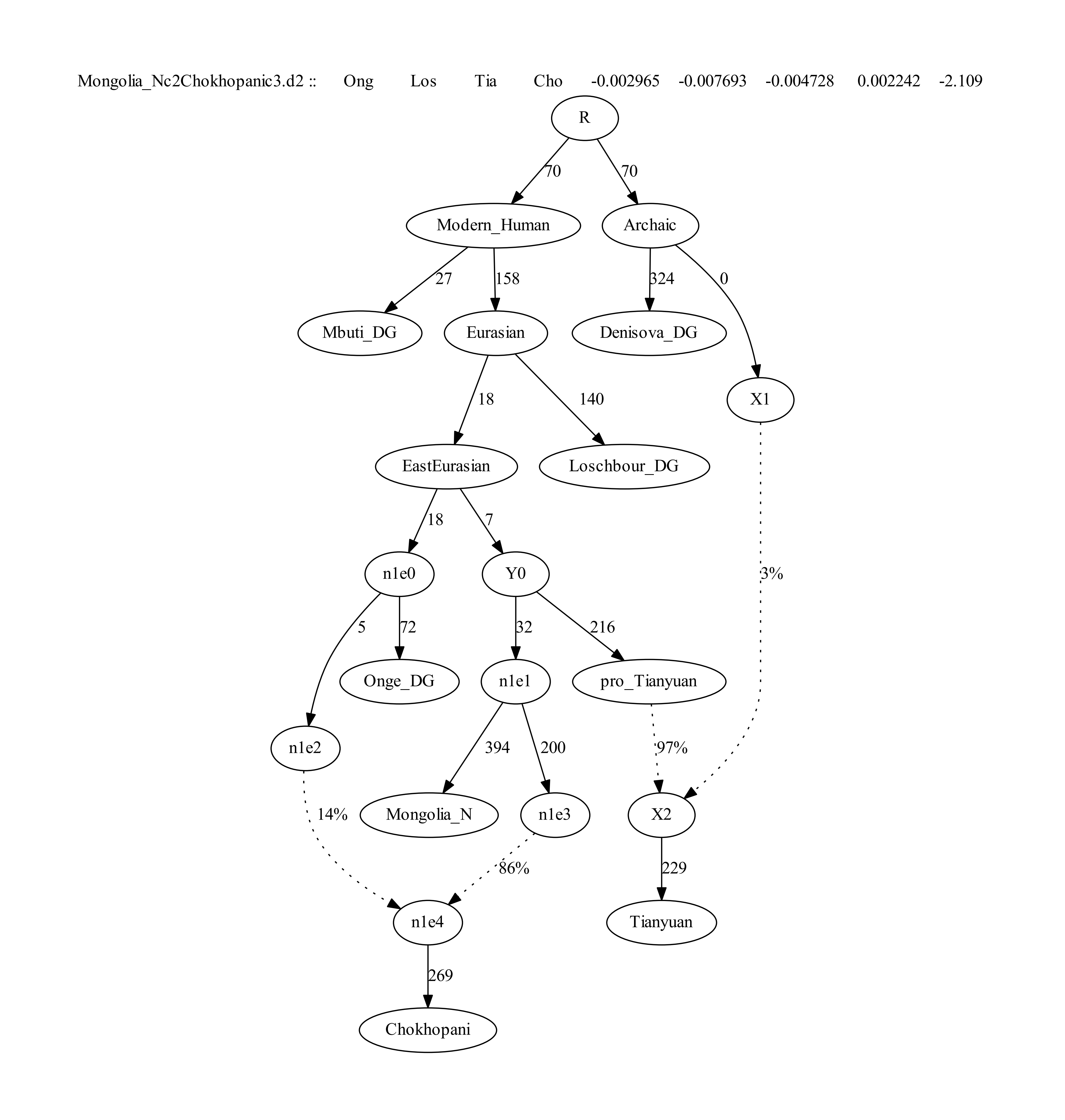

**Figure SP4: The model in which Chokhopani is admixed using 565749 SNPs, which is a fit to the data in the sense that there are no *f*-statistics more than |Z|>3 different between model and expectation. The likelihood score is 12.16 and the worst Z-score is for *f*-statistic (Onge, Loschbour; Tianyuan, Chokhopani) = -2.109.**

We then added Taiwan_Hanben onto all the possible edges of the graph that modeled Chokhopani sample is admixed. We obtained 61 fitted graphs for modeling Taiwan_Hanben samples as admixed as shown below in Table SP1. Forty-six of the models in Table SP1 involve Taiwan_Hanben and Chokhopani or Taiwan_Hanben and Mongolia_N sharing the majority of their ancestry, which is not consistent with the MSMC results that Ulchi (related to Mongolia_N/Boisman) and Tibetans share ancestors more recently than do Tibetan and Ami/Atayal. We were able to fit Taiwan_Hanben into the graph without invoking an additional admixture event, but with the worst Z-score of *f*-statistic (Onge, Loschbour; Tianyuan, Taiwan_Hanben) = -2.907 for the model “Taiwan_Hanbene1”. The large Z-score could be interpreted as suggesting unmodeled affinity between Onge and Taiwan_Hanben. We obtain three fitted graphs consistent with the MSMC results with lower maximum Z-scores by modeling Taiwan_Hanben as having Onge related admixture: Taiwan_Hanbene1.e4, Taiwan_Hanbene0.e1 and Taiwan_Hanbene1.e2. We show the unadmixed model “Taiwan_Hanbene1” and the best possible model “Taiwan_Hanbene1.e4” in Figure SP5.

**Table SP1. The fitted models for modeling Taiwan_Hanben samples onto all the possible edges of the graph in Figure SP4.**

| Model | likelihood  score | worst  Z-score | Note |
| --- | --- | --- | --- |
| Taiwan_Hanbend1.e6 | 12.733 | -2.190 | not consistent with MSMC |
| Taiwan_Hanbenc4.e6 | 13.034 | -2.106 | not consistent with MSMC |
| Taiwan_Hanbene1.e6 | 13.275 | -2.085 | not consistent with MSMC |
| Taiwan_Hanbene1.e4 | 13.771 | -2.224 |  |
| Taiwan_Hanbend0.e6 | 13.776 | -2.230 | not consistent with MSMC |
| Taiwan_Hanbene0.e1 | 14.438 | -2.329 |  |
| Taiwan_Hanbene1.e2 | 14.47 | -2.441 |  |
| Taiwan_Hanbene0.e5 | 14.893 | -2.150 | not consistent with MSMC |
| Taiwan_Hanbene0.e6 | 15.35 | -2.360 | not consistent with MSMC |
| Taiwan_Hanbenc0.e6 | 15.399 | -2.361 | not consistent with MSMC |
| Taiwan_Hanbene0.e3 | 15.516 | -2.406 | not consistent with MSMC |
| Taiwan_Hanbend0.e5 | 15.997 | -2.906 | not consistent with MSMC |
| Taiwan_Hanbend1.e5 | 16.111 | -2.907 | not consistent with MSMC |
| Taiwan_Hanbene1.e5 | 16.146 | -2.907 | not consistent with MSMC |
| Taiwan_Hanbenb0.e1 | 16.796 | -2.910 |  |
| Taiwan_Hanbene4.e5 | 17.004 | -2.765 | not consistent with MSMC |
| Taiwan_Hanbene2.e5 | 17.017 | -2.765 | not consistent with MSMC |
| Taiwan_Hanbenc1.e6 | 17.097 | -2.177 | not consistent with MSMC |
| Taiwan_Hanbend1.e1 | 17.133 | -2.907 | internal zero-length branch |
| Taiwan_Hanbend1.e3 | 17.139 | -2.907 | not consistent with MSMC |
| Taiwan_Hanbene1.e3 | 17.143 | -2.907 | not consistent with MSMC |
| Taiwan_Hanbene4.e6 | 17.152 | -2.751 | not consistent with MSMC |
| Taiwan_Hanbene2.e6 | 17.154 | -2.769 | not consistent with MSMC |
| Taiwan_Hanbene3.e4 | 17.162 | -2.760 | not consistent with MSMC |
| Taiwan_Hanbenb3.e6 | 17.185 | -2.160 | not consistent with MSMC |
| Taiwan_Hanbend0.e3 | 17.242 | -2.908 | not consistent with MSMC |
| Taiwan_Hanbene1 | 17.242 | -2.907 |  |
| Taiwan_Hanbend0.e1 | 17.243 | -2.906 | internal zero-length branch |
| Taiwan_Hanbenb1.e1 | 17.266 | -2.906 | internal zero-length branch |
| Taiwan_Hanbenc1.e1 | 17.268 | -2.907 |  |
| Taiwan_Hanbenb3.e1 | 17.295 | -2.909 |  |
| Taiwan_Hanbenb2.e1 | 17.316 | -2.906 |  |
| Taiwan_Hanbenc0.e1 | 17.333 | -2.907 |  |
| Taiwan_Hanbena0.e1 | 17.363 | -2.907 |  |
| Taiwan_Hanbena1.e1 | 17.372 | -2.905 |  |
| Taiwan_Hanbenc0.e5 | 17.931 | -2.912 | not consistent with MSMC |
| Taiwan_Hanbenc0.e3 | 18.136 | -2.910 | not consistent with MSMC |
| Taiwan_Hanbenc4.e1 | 18.395 | -2.917 |  |
| Taiwan_Hanbenb2.e6 | 18.503 | -2.446 | not consistent with MSMC |
| Taiwan_Hanbena0.e6 | 18.675 | 2.455 | not consistent with MSMC |
| Taiwan_Hanbenb0.e6 | 18.705 | -2.503 | not consistent with MSMC |
| Taiwan_Hanbena1.e6 | 18.804 | 2.472 | not consistent with MSMC |
| Taiwan_Hanbenb1.e6 | 18.945 | 2.475 | not consistent with MSMC |
| Taiwan_Hanbenc4.e5 | 19.307 | -2.927 | not consistent with MSMC |
| Taiwan_Hanbenc4.e3 | 19.31 | -2.928 | not consistent with MSMC |
| Taiwan_Hanbene6 | 19.564 | 2.525 | not consistent with MSMC |
| Taiwan_Hanbene3.e6 | 19.643 | 2.534 | not consistent with MSMC |
| Taiwan_Hanbene5.e6 | 19.722 | 2.522 | not consistent with MSMC |
| Taiwan_Hanbenb0.e5 | 24.374 | -2.961 | not consistent with MSMC |
| Taiwan_Hanbenb0.e3 | 24.383 | -2.967 | not consistent with MSMC |
| Taiwan_Hanbenb1.e5 | 24.728 | -2.975 | not consistent with MSMC |
| Taiwan_Hanbenb1.e3 | 24.731 | -2.975 | not consistent with MSMC |
| Taiwan_Hanbenb2.e5 | 24.768 | -2.976 | not consistent with MSMC |
| Taiwan_Hanbenb2.e3 | 24.77 | -2.974 | not consistent with MSMC |
| Taiwan_Hanbena0.e5 | 24.778 | -2.977 | not consistent with MSMC |
| Taiwan_Hanbena0.e3 | 24.781 | -2.976 | not consistent with MSMC |
| Taiwan_Hanbena1.e5 | 24.793 | -2.974 | not consistent with MSMC |
| Taiwan_Hanbena1.e3 | 24.795 | -2.977 | not consistent with MSMC |
| Taiwan_Hanbene3.e5 | 24.854 | -2.939 | not consistent with MSMC |
| Taiwan_Hanbene5 | 25.046 | -2.947 | not consistent with MSMC |
| Taiwan_Hanbene3 | 25.049 | -2.945 | not consistent with MSMC |

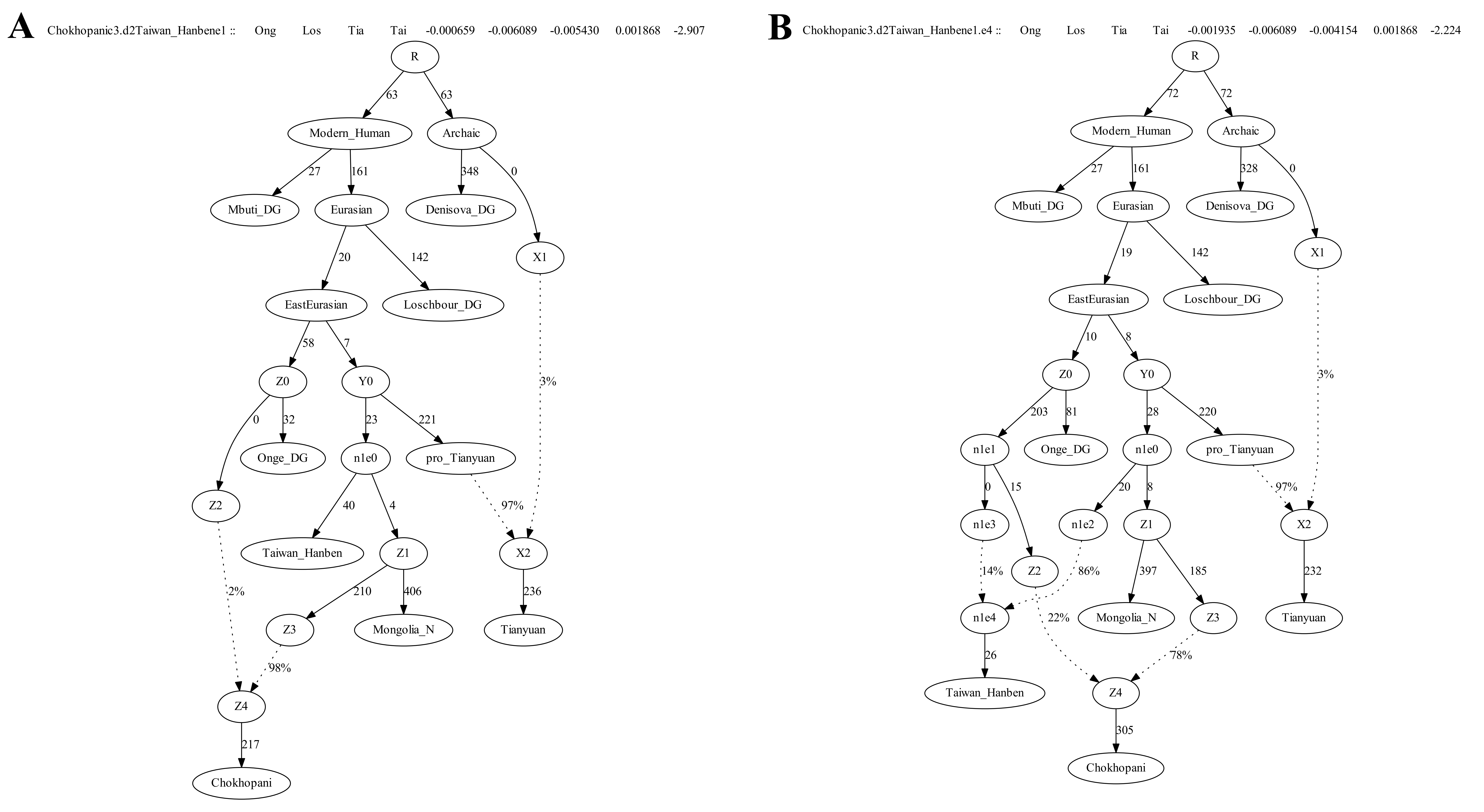

**Figure SP5: The unadmixed model “Taiwan_Hanbene1” and the model “Taiwan_Hanbene1.e4” with the lowest likelihood score and least mismatching *f*-statistics (the worst Z-score of *f*-statistic (Onge, Loschbour; Tianyuan, Taiwan_Hanben) = -2.224) and also consistent with the MSMC results.** **The number of SNPs used is 598829.**

We then added Japan_Jomon onto all the possible edges of the graph“Taiwan_Hanbene1.e4” but failed to find fitted models with no additional admixture events. We obtained eight fitted graphs that model Japan_Jomon as admixed as shown in Table SP2. The model with the lowest likelihood score and Z-score suggests that Jomon is an admixture deriving 56% of the ancestry from a lineage on the Mongolia_Neolithic side that also contributed to Taiwan_Hanben and 44% from a basal lineage on the Onge side but branched off earlier than Onge (Figure SP6). The model “Japan_Jomond0.f6” suggests Jomon derive 56% of the ancestry from Taiwan_Hanben related group and the remaining 44% from a basal lineage that branched off earlier than the separation of Tianyuan with other East Asians. The model “Japan_Jomonf1.f4” suggests Jomon derive 55% of the ancestry from Taiwan_Hanben related lineage on the Mongolia_N side and the remaining 45% from a lineage on the Onge side (Figure SP6).

**Table SP2. The fitted models for modeling Japan_Jomon samples as admixed.**

| Model | likelihood score | worst Z-score | Note |
| --- | --- | --- | --- |
| Japan_Jomone0.f4 | 24.358 | 2.800 |  |
| Japan_Jomond0.f6 | 25.503 | 2.800 |  |
| Japan_Jomonc0.f4 | 25.518 | 2.800 | internal zero-length branch |
| Japan_Jomond0.f4 | 25.524 | 2.800 | internal zero-length branch |
| Japan_Jomonf1.f4 | 25.685 | -2.902 |  |
| Japan_Jomone0.f6 | 26.046 | 2.800 | internal zero-length branch |
| Japan_Jomonc0.f6 | 26.098 | 2.800 | internal zero-length branch |
| Japan_Jomone2.f4 | 27.133 | -2.935 | internal zero-length branch |

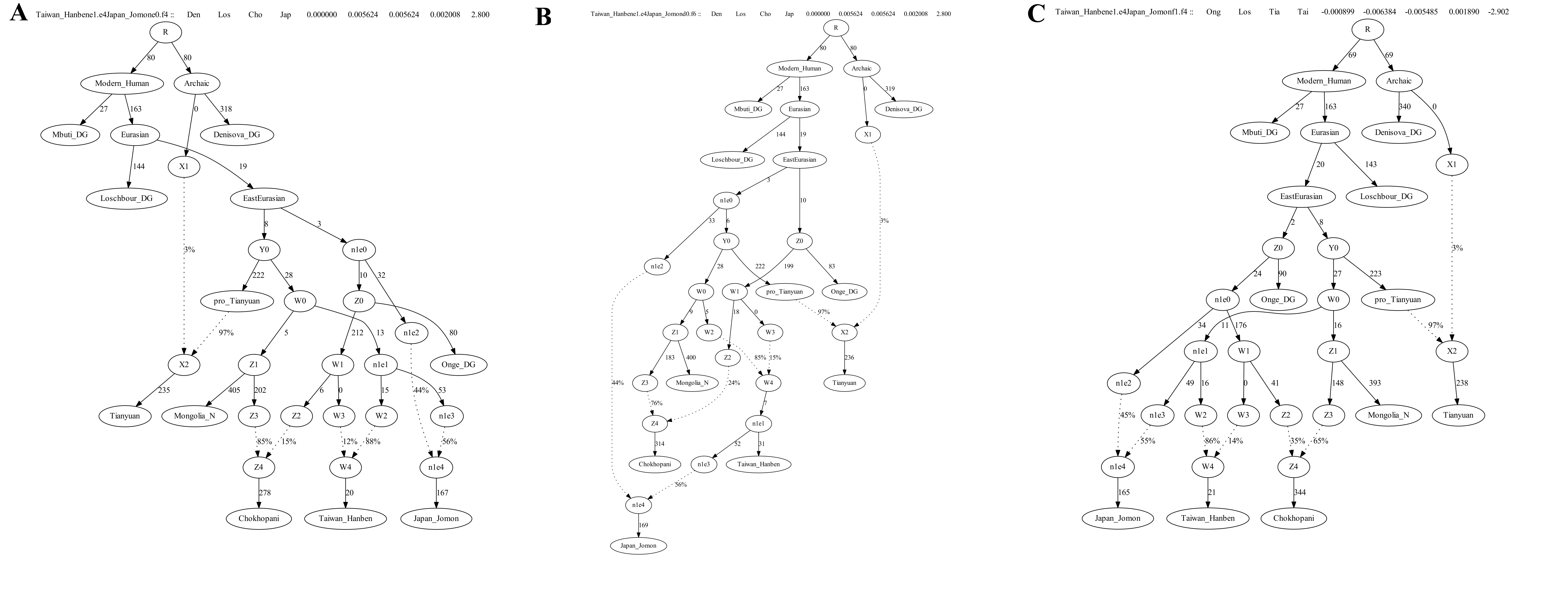

**Figure SP6. (A) The model “Japan_Jomone0.f4” with the lowest likelihood score and Z-score by modeling Japan_Jomon is admixed. The worst Z-score of the *f*-statistic (Denisovan, Loschbour; Chokhopani, Japan_Jomon) = 2.800.** **(B) The model “Japan_Jomond0.f6”. (C) The model “Japan_Jomonf1.f4”. The number of SNPs used is 580671.**

We added Boisman onto all the possible edges of the model “Japan_Jomone0.f4” and obtained 7 fitted models as shown below in the Table SP3. There are 5 models with internal zero-length branches. We show the model “Boisman_MNe3.g5” with lowest likelihood score and Z-score in Figure SP7.

**Table SP3. The fitted models for modeling Boisman_MN samples as admixed.**

| Model | likelihood score | worst Z-score | Note |
| --- | --- | --- | --- |
| Boisman_MNe3.g5 | 32.451 | 2.751 |  |
| Boisman_MNf2.g5 | 40.888 | 2.883 | internal zero-length branch |
| Boisman_MNe5.g5 | 40.892 | 2.883 | internal zero-length branch |
| Boisman_MNf0.g5 | 44.438 | 2.879 | internal zero-length branch |
| Boisman_MNg1.g5 | 44.459 | 2.873 | internal zero-length branch |
| Boisman_MNg1.g6 | 45.125 | 2.895 |  |
| Boisman_MNg1.g4 | 46.055 | 2.910 | internal zero-length branch |

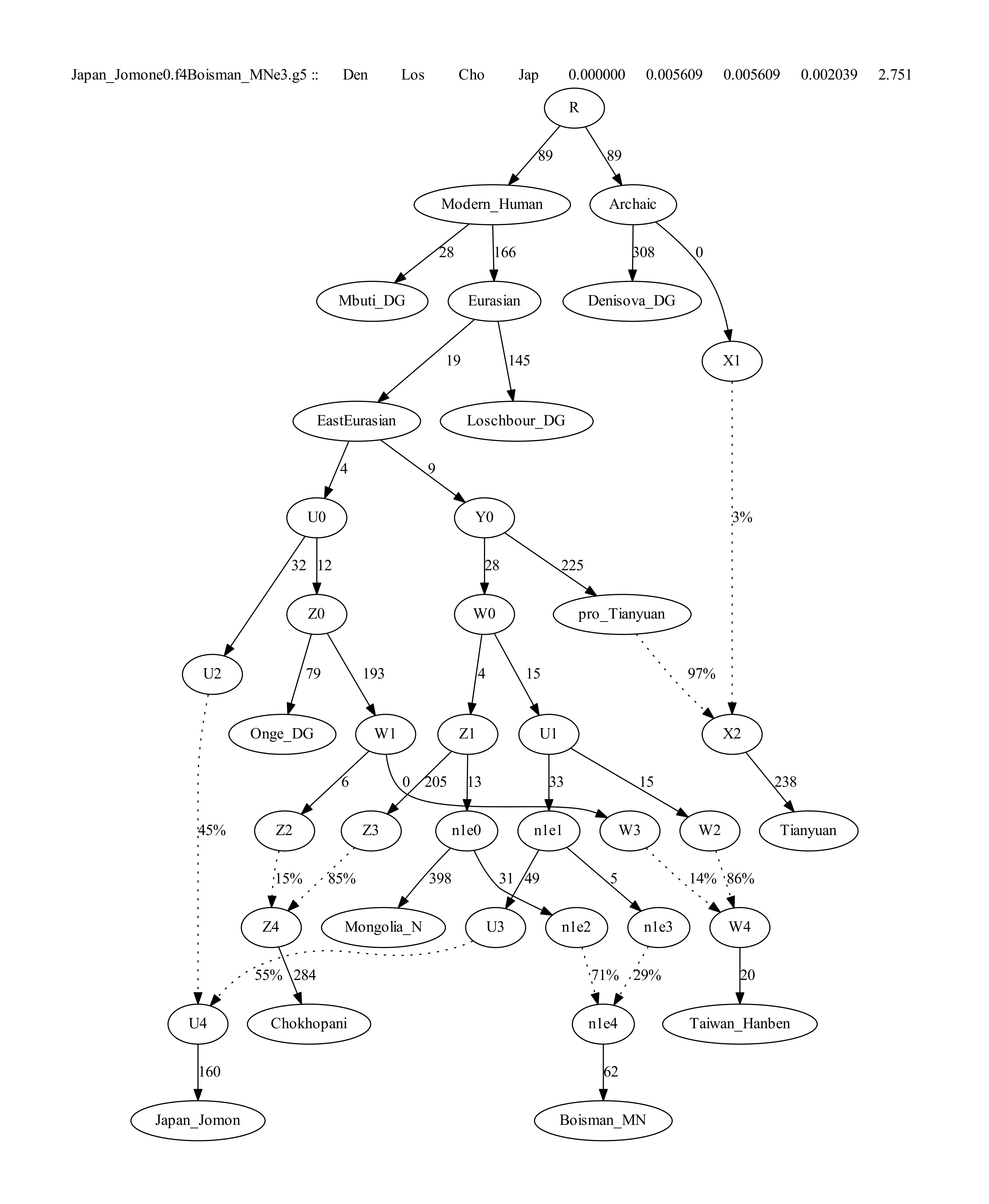

**Figure SP7: The fitted model with lowest likelihood score and Z-score without internal zero-length branches by adding Boisman_MN onto all the possible edges of the model “Japan_Jomone0.f4”. The likelihood score and the worst Z-score are shown in Table SP3. The number of SNPs used is 577995.**

We then added Wuzhuangguoliang onto all the possible edges of the model “Boisman_MNe3.g5”, but failed to find fitted models with no additional admixture events or with only one additional admixture event. We show the four graphs with the top four lowest likelihood-scores (<40) and Z-scores in Figure SP8.

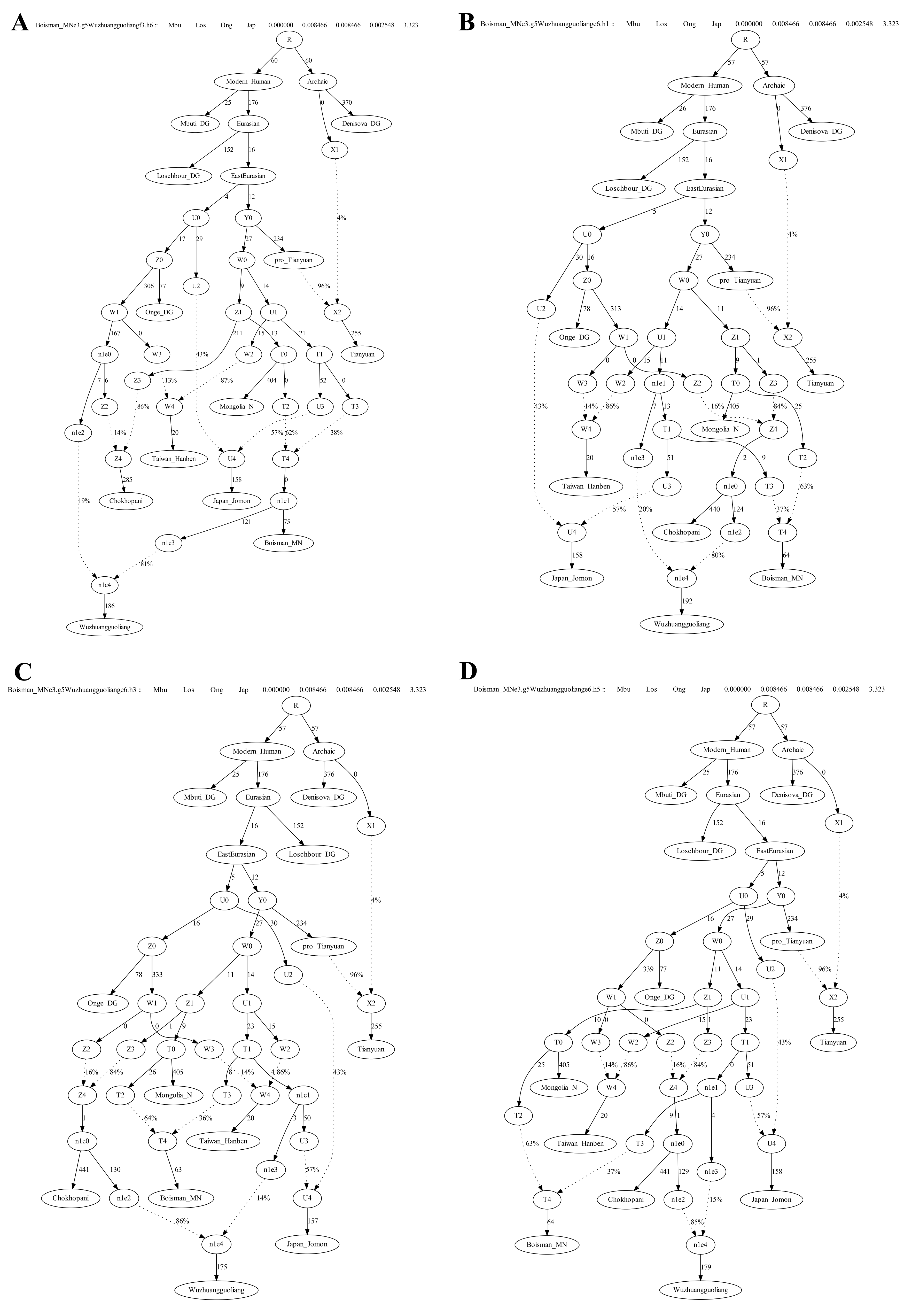

**Figure SP8. The four graphs with the top four lowest likelihood-scores (<40) and Z-scores by adding Wuzhuangguoliang onto all the possible edges of the model “Boisman_MNe3.g5”. The number of SNPs used is 98316. A: model Wuzhuangguoliangf3.h6 with a likelihood-score of 38.06 and Z-score of 3.323; B: Wuzhuangguoliange6.h1 with a likelihood-score of 39.233 and Z-score of 3.323; C: Wuzhuangguoliange6.h3 with a likelihood-score of 39.428 and Z-score of 3.323; D: model Wuzhuangguoliange6.h5 with a likelihood-score of 39.466 and Z-score of 3.323.**

We then added Tibetan onto all the possible edges of the model “Boisman_MNe3.g5” and got five fitted models with |Z-score|<3 as shown in Table SP4. The model “Tibetan.DGc1.e6” with the lowest Z-score and likelihood-score was shown in Figure SP9, which suggest Tibetan derive 95% ancestry from Chokhopani related lineage and the remaining 5% from groups related to West Eurasians.

**Table SP4. The fitted models for modeling Tibetan samples as admixed.**

| Model | likelihood score | worst Z score | Note |
| --- | --- | --- | --- |
| Tibetan.DGc1.e6 | 43.002 | 2.752 |  |
| Tibetan.DGb3.e6 | 44.156 | 2.752 | internal zero-length branch |
| Tibetan.DGc0.e6 | 44.197 | 2.752 | internal zero-length branch |
| Tibetan.DGe6.g4 | 44.211 | -2.879 |  |
| Tibetan.DGe6.g6 | 46.644 | -2.852 |  |

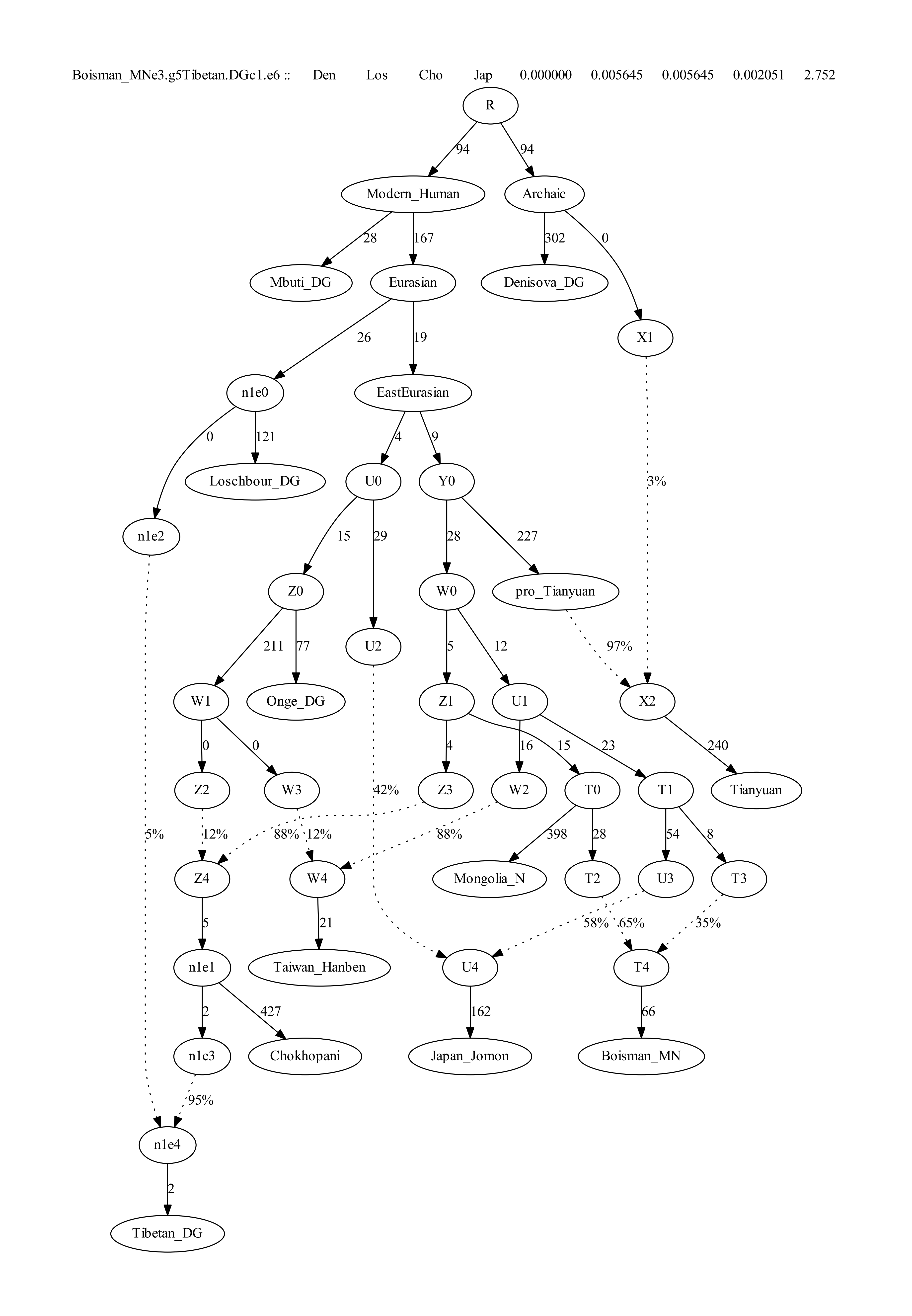

**Figure SP9.the model with the lowest Z-score and likelihood-score by adding Tibetan onto all the possible edges of the model “Boisman_MNe3.g5”.**

We then added present-day Ami, Han and Japanese onto all the possible edges of model Boisman_MNe3.g5.

We got three fitted models for Ami, all of which suggest Ami share 95%-98% of the ancestry with Taiwan_Hanben samples (Figure SP10).

We got two fitted models for Han Chinese as shown in Figure SP11: one model “Han.DGe6.g3” suggests Han Chinese derive 63% of the ancestry from Chokhopani related group and the remining 37% from a lineage that also contributed to Taiwan_Hanben, the other model suggests Han Chinese derive 77% of the ancestry from Boisman related group and the remining 23% from a lineage that also contributed to Taiwan_Hanben.

We failed to find fitted models for Japanese with worst |Z|-score less than 3, but find three models with |Z|-scores less than 4 as shown in Figure SP12. All the three models suggest Japanese derive about 68%-69% of the ancestry from Chokhopani related groups and the remining from Jomon related lineage.

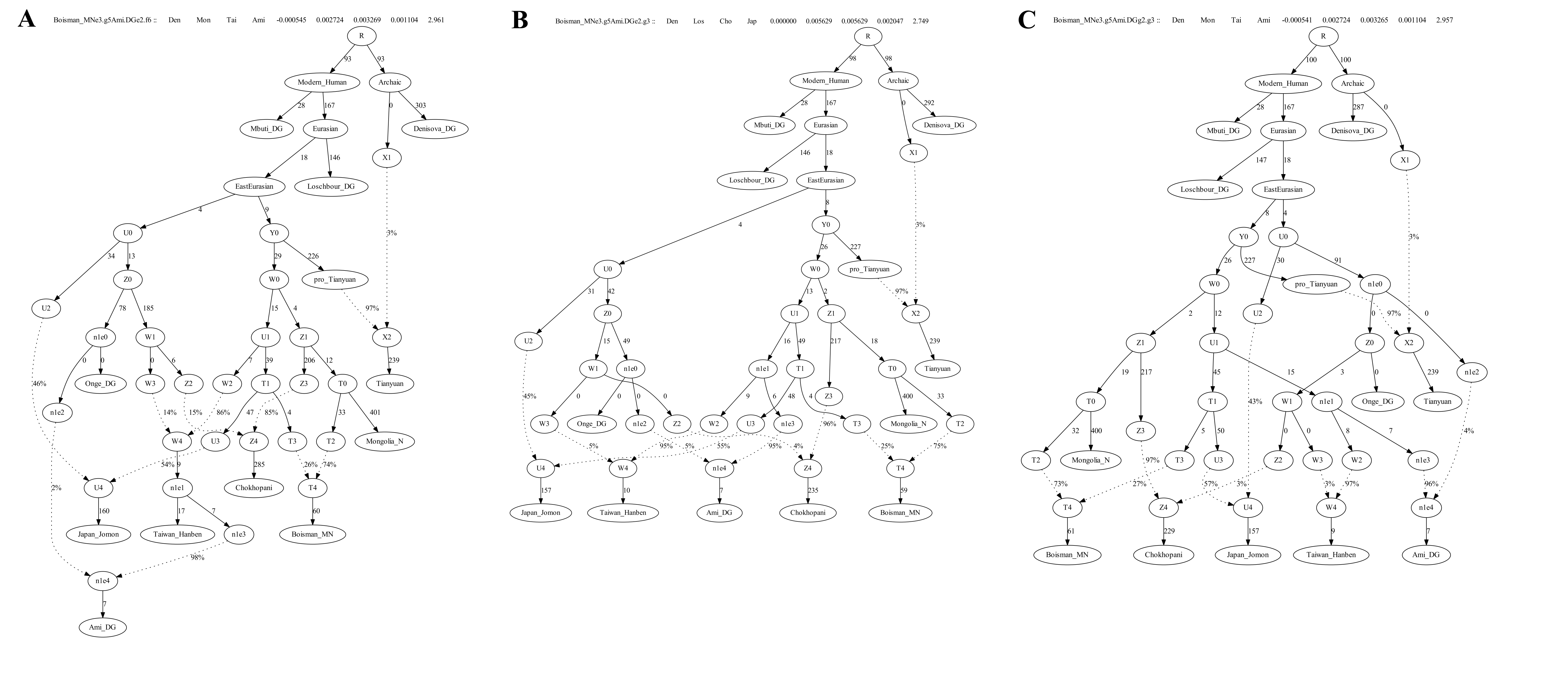

**Figure SP10. the models with the lowest Z-score and likelihood-score by adding Ami onto all the possible edges of the model “Boisman_MNe3.g5”.**

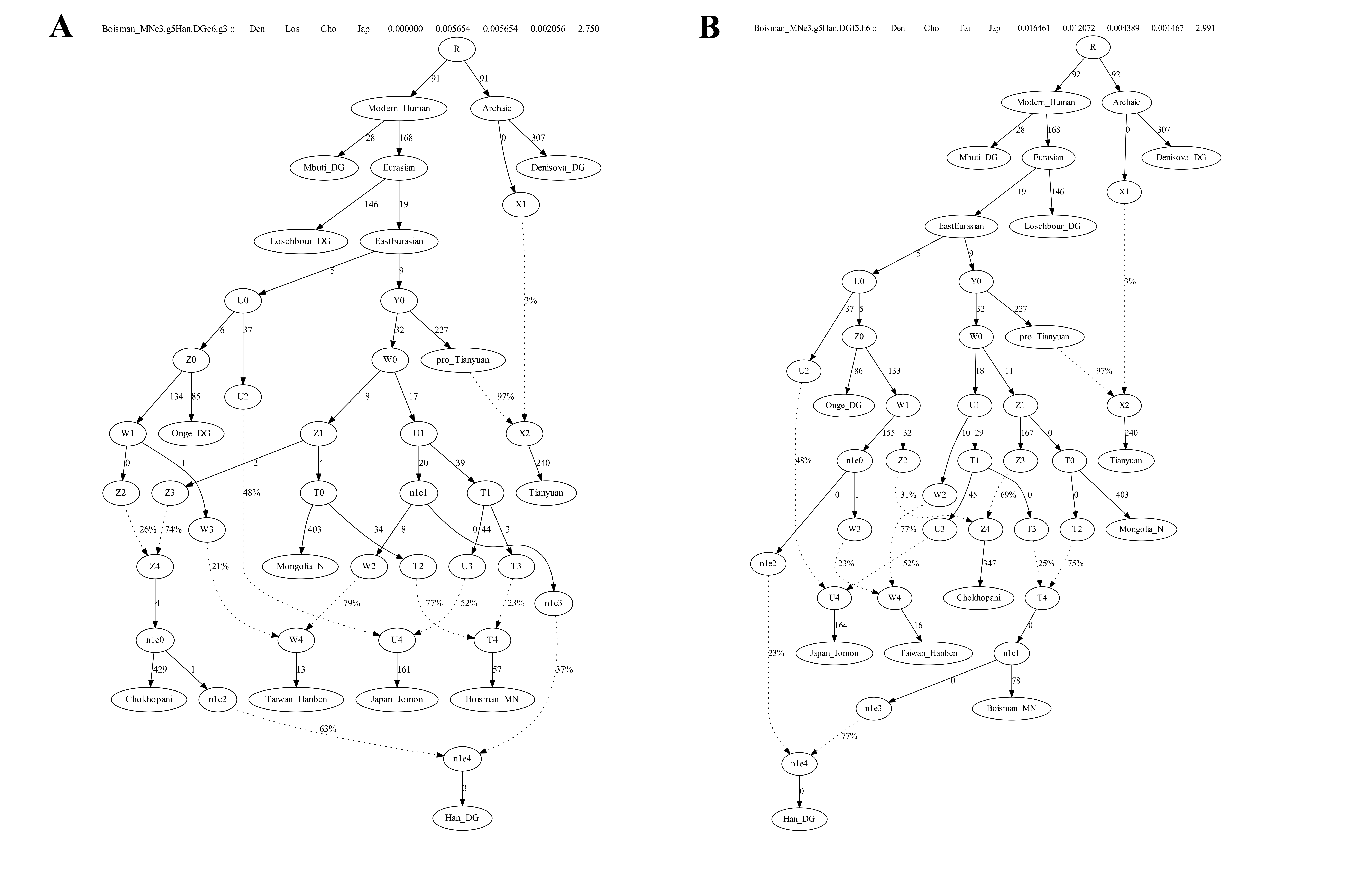

**Figure SP11. the models with the lowest Z-score and likelihood-score by adding Han Chinese onto all the possible edges of the model “Boisman_MNe3.g5”.**

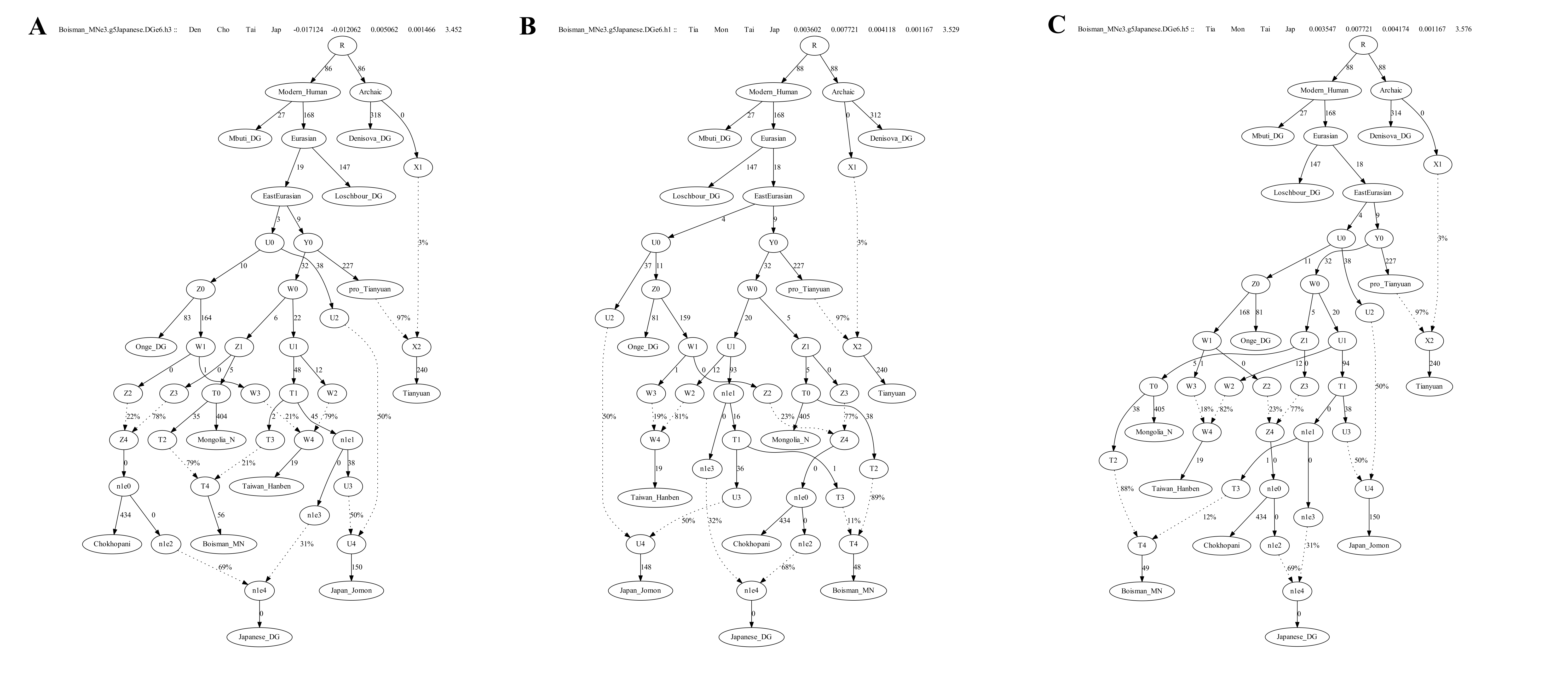

**Figure SP12. the models with the lowest Z-score and likelihood-score by adding Japanese onto all the possible edges of the model “Boisman_MNe3.g5”.**

**Transversions only**

We next restrict the analysis to transversion sites only. We started with the same skeleton phylogenetic tree consisting of Denisovan, Mbuti, Loschbour, Tianyuan, and Onge with one admixture from Denisovan to Tianyuan as shown in Figure SP2. The tree generated using transversions was an adequate fit with the likelihood score of 0.637 and the worst Z-score of *f*-statistic (Mbuti, Denisovan; Onge, Loschbour) = 0.596 (Figure SP13).

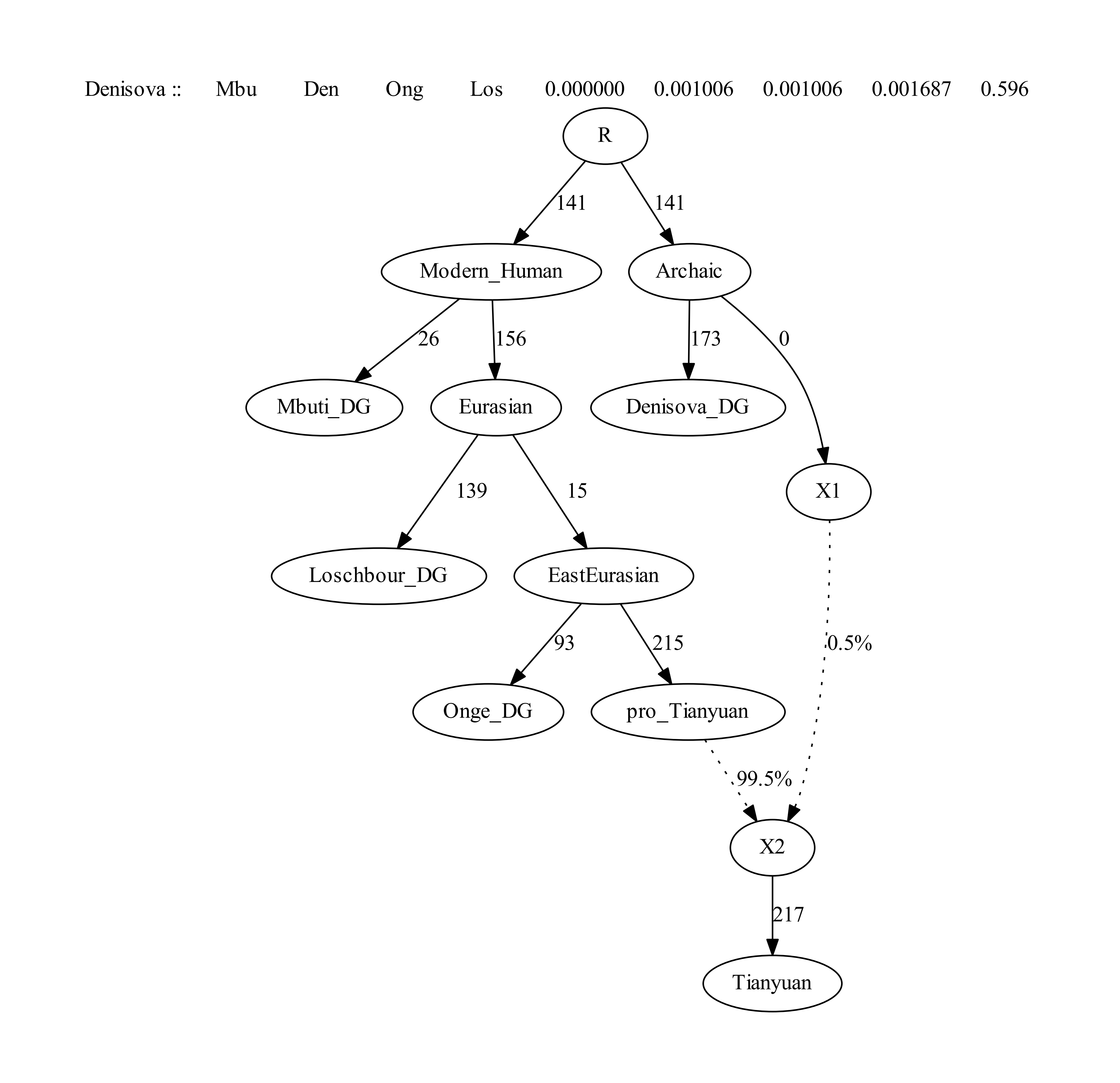

**Figure SP13: Skeleton Admixture Graph with one admixture using 124903 transversion sites, which is a fit to the genetic data in the sense that there are no *f*-statistics more than |Z|>3 different between model and expectation. The drifts along edges in this figure and the following ones are multiplied by 1000.**

We then added Mongolia_Neolithic onto all the possible edges of the graph and found a fitted model with no additional admixture event of assigning Mongolia_Neolithic as a sister branch of Tianyuan (Figure SP14).

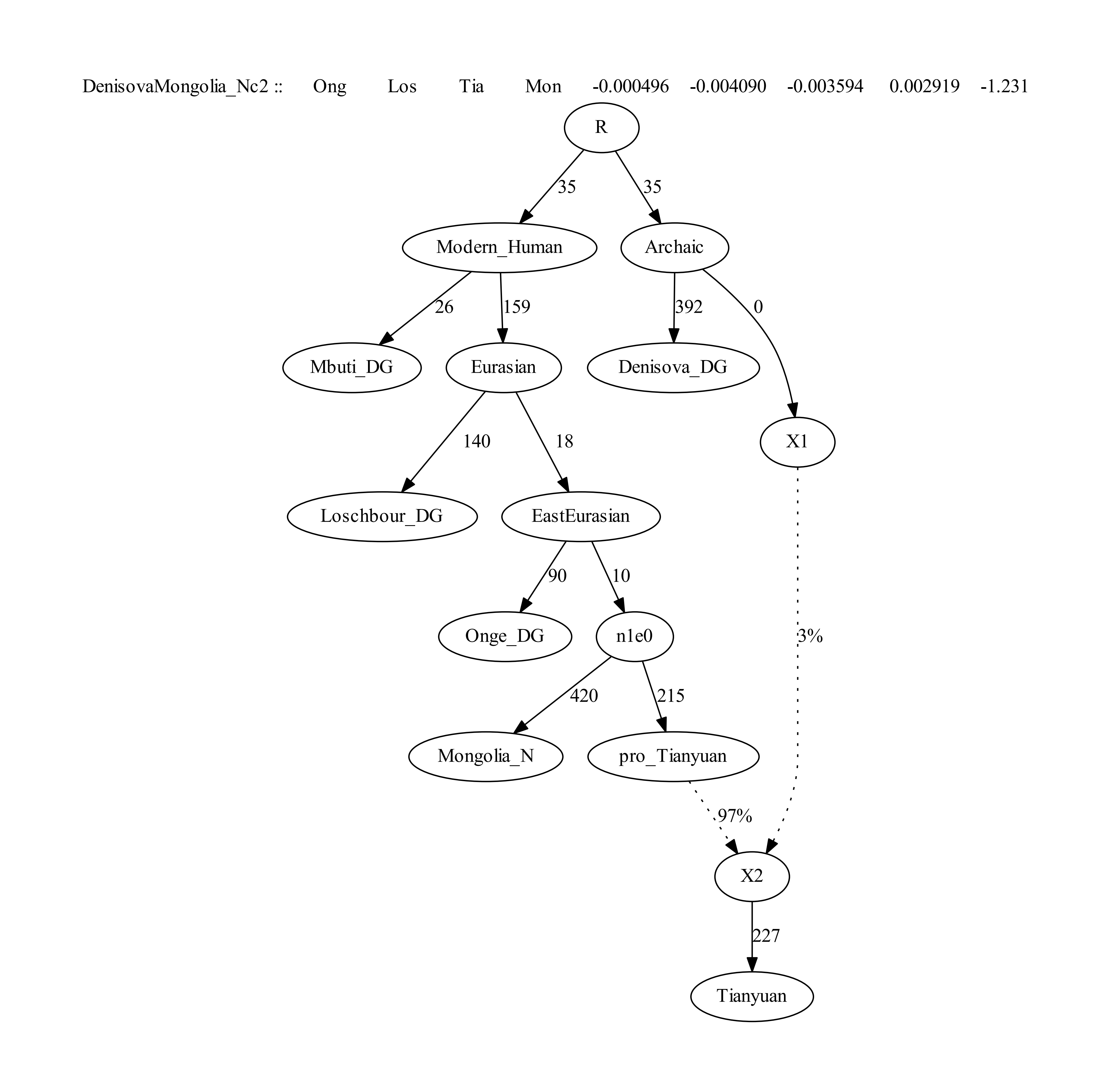

**Figure SP14: The best fitting tree of adding Mongolia_Neolithic without admixture was an adequate fit as in the sense that there are no *f*-statistics more than |Z|>3 different between model and expectation. The likelihood score is 4.523 and the worst Z-score of *f*-statistic (Onge, Loschbour; Tianyuan, Mongolia_N) = -1.231. The number of transversion sites used is 104964.**

We then added Chokhopani onto all the possible edges of the graph in Figure SP14. We were able to fit Chokhopani into the graph as a sister lineage of Mongolia_N without invoking an additional admixture event, but with the worst Z-score of f-statistic (Onge, Loschbour; Tianyuan, Chokhopani) = -2.810 for the model “Chokhopanid2”. The large Z-score could be interpreted as suggesting unmodeled affinity between Onge and Chokhopani. We can also fit Chokhopani into the graph as admixed as shown in Table SP5. The model “Chokhopanic3.d2” with lowest likelihood score and Z-score in Figure SP15 suggests that Chokhopani is an admixture deriving 23% and 77% from Onge and Mongolia_Neolithic related lineages, respectively.

**Table SP5. The fitted models for modeling Chokhopani sample as admixed.**

| Model | likelihood score | worst Z score | Note |
| --- | --- | --- | --- |
| Chokhopanic3.d2 | 8.596 | -1.717 |  |
| Chokhopanic0.d2 | 11.779 | -2.811 | internal zero-length branch |
| Chokhopanid0.d2 | 11.787 | -2.811 |  |
| Chokhopanib2.d2 | 12.813 | -2.838 |  |
| Chokhopania1.d2 | 13.004 | -2.840 |  |
| Chokhopania0.d2 | 13.045 | -2.838 | internal zero-length branch |
| Chokhopanib3.d2 | 13.134 | -2.848 |  |
| Chokhopanib1.d2 | 13.382 | -2.819 | internal zero-length branch |
| Chokhopanic4.d2 | 13.387 | -2.819 | internal zero-length branch |
| Chokhopanid1.d2 | 13.408 | -2.811 | internal zero-length branch |
| Chokhopanid2 | 13.482 | -2.810 |  |
| Chokhopanic1.d2 | 13.487 | -2.830 | internal zero-length branch |
| Chokhopanib0.d2 | 13.872 | -2.849 | internal zero-length branch |

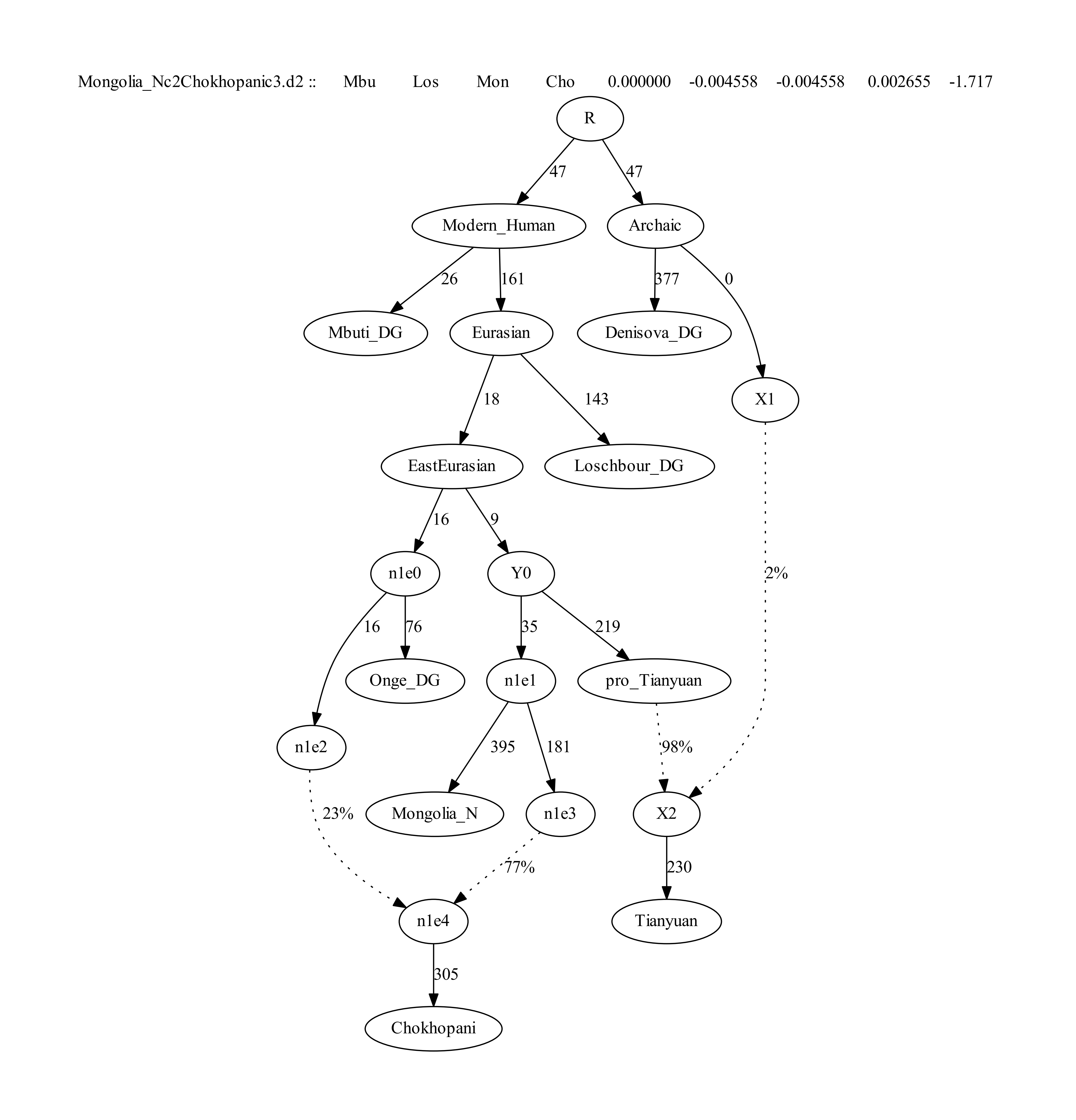

**Figure SP15: The model “Chokhopanic3.d2” in which Chokhopani is admixed using 105573 transversion sites, which is a fit to the data in the sense that there are no *f*-statistics more than |Z|>3 different between model and expectation. The likelihood score is 8.596 and the worst Z-score is for *f*-statistic (Mbuti, Loschbour; Mongolia_N, Chokhopani) = -1.717.**

We then added Taiwan_Hanben onto all the possible edges of the graph “Chokhopanic3.d2” that modeled Chokhopani sample is admixed. We obtained 4 fitted graphs for modeling Taiwan_Hanben as unadmixed and 62 fitted graphs for modeling Taiwan_Hanben samples as admixed as shown below in Table SP6. Fifty-one of the models in Table SP6 involve Taiwan_Hanben and Chokhopani or Taiwan_Hanben and Mongolia_N sharing the majority of their ancestry, which is not consistent with the MSMC results that Ulchi (related to Mongolia_N/Boisman) and Tibetans share ancestors more recently than do Tibetan and Ami/Atayal. Therefore, the only workable unadmixed model is “Taiwan_Hanbene1” with the worst Z-score of -2.250, which is lower than that of using all 1240K sites. The reason of a lower Z-score we suspect is due to the lower number of available SNPs used in the transversion-only analysis. Consistent with the analysis of using all sites, we observed the lowest likelihood score and Z-score in model “Taiwan_Hanbene1.e4” of suggesting Taiwan_Hanben deriving 14% of the ancestry from Onge related lineage. Here we show the unadmixed model “Taiwan_Hanbene1” and the best possible model “Taiwan_Hanbene1.e4” in Figure SP16.

**Table SP6. The fitted models for modeling Taiwan_Hanben samples onto all the possible edges of the graph in Figure SP14.**

| Model | likelihood score | | worst Z score | Note |
| --- | --- | --- | --- | --- |
| Taiwan_Hanbene1.e6 | | 10.684 | 2.032 | not consistent with MSMC |
| Taiwan_Hanbene1.e4 | | 10.782 | 2.032 |  |
| Taiwan_Hanbene0.e5 | | 11.440 | 2.032 | not consistent with MSMC |
| Taiwan_Hanbend1.e6 | | 11.617 | -2.169 | not consistent with MSMC |
| Taiwan_Hanbene0.e1 | | 11.623 | 2.032 |  |
| Taiwan_Hanbene1.e2 | | 11.624 | 2.032 | internal zero-length branch |
| Taiwan_Hanbenc4.e6 | | 11.681 | -2.168 | not consistent with MSMC |
| Taiwan_Hanbene3.e6 | | 11.717 | 2.032 | not consistent with MSMC, internal zero-length branch |
| Taiwan_Hanbene0.e3 | | 11.764 | 2.032 | not consistent with MSMC, internal zero-length branch |
| Taiwan_Hanbene5.e6 | | 11.836 | 2.032 | not consistent with MSMC, internal zero-length branch |
| Taiwan_Hanbene4.e5 | | 11.924 | 2.032 | not consistent with MSMC, internal zero-length branch |
| Taiwan_Hanbenc1.e6 | | 11.938 | -2.042 | not consistent with MSMC |
| Taiwan_Hanbene2.e5 | | 11.945 | 2.032 | not consistent with MSMC, internal zero-length branch |
| Taiwan_Hanbene2.e3 | | 11.985 | 2.032 | not consistent with MSMC, internal zero-length branch |
| Taiwan_Hanbend0.e5 | | 12.414 | -2.237 | not consistent with MSMC |
| Taiwan_Hanbenb0.e6 | | 12.579 | -2.174 | not consistent with MSMC, internal zero-length branch |
| Taiwan_Hanbene3.e4 | | 12.628 | -2.098 | not consistent with MSMC, internal zero-length branch |
| Taiwan_Hanbend0.e6 | | 12.808 | -2.178 | not consistent with MSMC, internal zero-length branch |
| Taiwan_Hanbend1.e5 | | 12.866 | -2.252 | not consistent with MSMC, internal zero-length branch |
| Taiwan_Hanbene1.e5 | | 12.873 | -2.253 | not consistent with MSMC |
| Taiwan_Hanbene6 | | 12.951 | -2.209 | not consistent with MSMC |
| Taiwan_Hanbenb3.e6 | | 12.967 | -2.150 | not consistent with MSMC |
| Taiwan_Hanbenb0.e1 | | 12.985 | -2.208 |  |
| Taiwan_Hanbenb1.e6 | | 12.985 | -2.206 | not consistent with MSMC, internal zero-length branch |
| Taiwan_Hanbene4.e6 | | 12.985 | -2.207 | not consistent with MSMC, internal zero-length branch |
| Taiwan_Hanbene0.e6 | | 12.997 | -2.205 | not consistent with MSMC, internal zero-length branch |
| Taiwan_Hanbenc0.e6 | | 13.003 | -2.160 | not consistent with MSMC, internal zero-length branch |
| Taiwan_Hanbene2.e6 | | 13.004 | -2.213 | not consistent with MSMC |
| Taiwan_Hanbenc0.e5 | | 13.014 | -2.217 | not consistent with MSMC, internal zero-length branch |
| Taiwan_Hanbenb2.e6 | | 13.038 | -2.206 | not consistent with MSMC |
| Taiwan_Hanbena1.e6 | | 13.084 | -2.212 | not consistent with MSMC |
| Taiwan_Hanbena0.e6 | | 13.139 | -2.186 | not consistent with MSMC |
| Taiwan_Hanbene1.e3 | | 13.256 | -2.244 | not consistent with MSMC, internal zero-length branch |
| Taiwan_Hanbenc1.e1 | | 13.285 | -2.236 |  |
| Taiwan_Hanbend1.e1 | | 13.334 | -2.246 | internal zero-length branch |
| Taiwan_Hanbene1 | | 13.337 | -2.250 |  |
| Taiwan_Hanbenb1.e1 | | 13.376 | -2.243 | internal zero-length branch |
| Taiwan_Hanbenc0.e1 | | 13.380 | -2.246 | internal zero-length branch |
| Taiwan_Hanbend0.e1 | | 13.383 | -2.251 | internal zero-length branch |
| Taiwan_Hanbend1.e3 | | 13.385 | -2.249 | not consistent with MSMC, internal zero-length branch |
| Taiwan_Hanbenc4.e1 | | 13.401 | -2.268 |  |
| Taiwan_Hanbenb2.e1 | | 13.428 | -2.263 |  |
| Taiwan_Hanbena1.e1 | | 13.431 | -2.263 |  |
| Taiwan_Hanbena0.e1 | | 13.433 | -2.277 |  |
| Taiwan_Hanbenb3.e1 | | 13.433 | -2.169 |  |
| Taiwan_Hanbend0.e3 | | 13.438 | -2.254 | not consistent with MSMC, internal zero-length branch |
| Taiwan_Hanbenc0.e3 | | 13.449 | -2.244 | not consistent with MSMC, internal zero-length branch |
| Taiwan_Hanbenb0.e3 | | 13.939 | -2.235 | not consistent with MSMC, internal zero-length branch |
| Taiwan_Hanbenb0.e5 | | 13.942 | -2.233 | not consistent with MSMC, internal zero-length branch |
| Taiwan_Hanbenc1.e3 | | 14.086 | -2.25 | not consistent with MSMC, internal zero-length branch |
| Taiwan_Hanbenc1.e5 | | 14.088 | -2.242 | not consistent with MSMC, internal zero-length branch |
| Taiwan_Hanbene3.e5 | | 14.176 | -2.263 | not consistent with MSMC, internal zero-length branch |
| Taiwan_Hanbene3 | | 14.363 | -2.268 | not consistent with MSMC, internal zero-length branch |
| Taiwan_Hanbene5 | | 14.365 | -2.276 | not consistent with MSMC, internal zero-length branch |
| Taiwan_Hanbenb1.e3 | | 14.396 | -2.29 | not consistent with MSMC, internal zero-length branch |
| Taiwan_Hanbenb1.e5 | | 14.397 | -2.274 | not consistent with MSMC, internal zero-length branch |
| Taiwan_Hanbenb3.e5 | | 14.426 | -2.292 | not consistent with MSMC, internal zero-length branch |
| Taiwan_Hanbenb3.e3 | | 14.432 | -2.29 | not consistent with MSMC, internal zero-length branch |
| Taiwan_Hanbenb2.e3 | | 14.446 | -2.28 | not consistent with MSMC, internal zero-length branch |
| Taiwan_Hanbenb2.e5 | | 14.448 | -2.277 | not consistent with MSMC, internal zero-length branch |
| Taiwan_Hanbena0.e3 | | 14.458 | -2.300 | not consistent with MSMC, internal zero-length branch |
| Taiwan_Hanbena1.e3 | | 14.458 | -2.287 | not consistent with MSMC, internal zero-length branch |
| Taiwan_Hanbena0.e5 | | 14.461 | -2.298 | not consistent with MSMC, internal zero-length branch |
| Taiwan_Hanbena1.e5 | | 14.461 | -2.295 | not consistent with MSMC, internal zero-length branch |
| Taiwan_Hanbenc4.e3 | | 14.899 | -2.247 | not consistent with MSMC, internal zero-length branch |
| Taiwan_Hanbenc4.e5 | | 14.902 | -2.247 | not consistent with MSMC, internal zero-length branch |

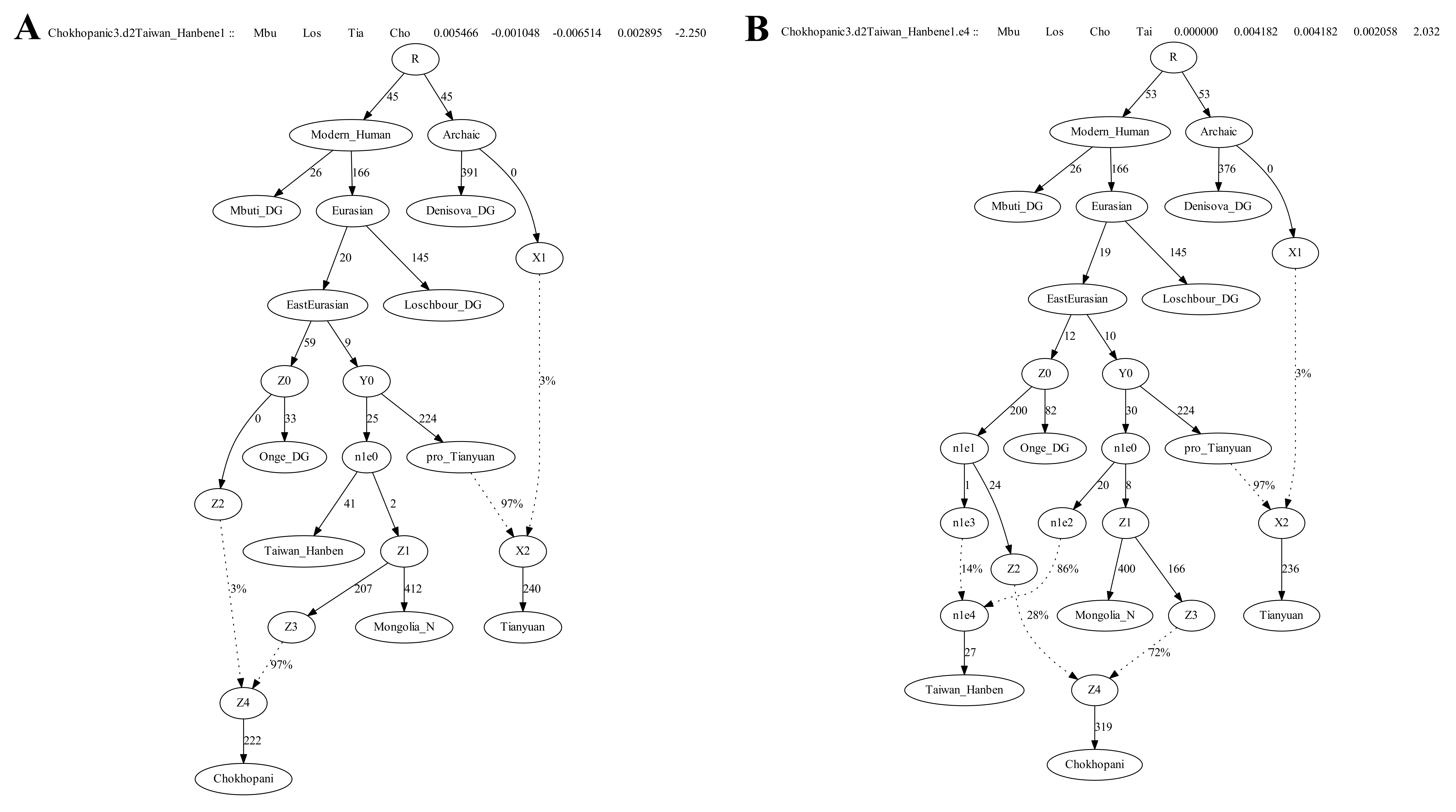

**Figure SP16: The unadmixed model “Taiwan_Hanbene1” (A) and the model “Taiwan_Hanbene1.e4” (B) with the lowest likelihood score and least mismatching *f*-statistics (the worst Z-score of *f*-statistic (Mbuti, Loschbour; Chokhopani, Taiwan_Hanben) = 2.032) and also consistent with the MSMC results.** **The number of transversion sites used is 109411.**

We then added Japan_Jomon onto all the possible edges of the graph“Taiwan_Hanbene1.e4” but failed to find fitted models with no additional admixture events. We obtained 33 fitted graphs that model Japan_Jomon as admixed as shown in Table SP7. The model “Japan_Jomone0.f4” with the lowest likelihood score and Z-score without internal zero-length branches suggests that Jomon is an admixture deriving 56% and 44% from Mongolia_Neolithic and Onge related lineages, respectively. We also show other four fitted models without internal zero-length branches in Figure SP17.

**Table SP7. The fitted models for modeling Japan_Jomon samples as admixed.**

| Model | likelihood score | worst Z score | Note |
| --- | --- | --- | --- |
| Japan_Jomone0.f4 | 19.987 | -2.849 |  |
| Japan_Jomond0.f6 | 20.450 | -2.849 |  |
| Japan_Jomone0.f6 | 20.484 | -2.849 |  |
| Japan_Jomonc0.f6 | 20.634 | -2.849 | internal zero-length branch |
| Japan_Jomonf1.f4 | 20.965 | -2.849 |  |
| Japan_Jomonc0.f4 | 21.052 | -2.849 | internal zero-length branch |
| Japan_Jomond0.f4 | 21.060 | -2.849 | internal zero-length branch |
| Japan_Jomone2.f4 | 21.584 | -2.849 | internal zero-length branch |
| Japan_Jomone3.f5 | 22.638 | -2.849 | internal zero-length branch |
| Japan_Jomonf1.f6 | 22.820 | -2.849 | internal zero-length branch |
| Japan_Jomone2.f6 | 22.835 | -2.849 | internal zero-length branch |
| Japan_Jomone5.f5 | 23.538 | -2.849 | internal zero-length branch |
| Japan_Jomonf2.f5 | 23.555 | -2.849 | internal zero-length branch |
| Japan_Jomone6.f5 | 23.557 | -2.849 | internal zero-length branch |
| Japan_Jomonf3.f4 | 23.562 | -2.849 | internal zero-length branch |
| Japan_Jomonf3.f6 | 23.562 | -2.849 | internal zero-length branch |
| Japan_Jomonf4.f5 | 23.562 | -2.849 | internal zero-length branch |
| Japan_Jomonf5.f6 | 23.562 | -2.849 | internal zero-length branch |
| Japan_Jomond1.f6 | 23.846 | -2.849 | internal zero-length branch |
| Japan_Jomonf0.f6 | 23.884 | -2.849 | internal zero-length branch |
| Japan_Jomone3.f1 | 24.017 | -2.849 | internal zero-length branch |
| Japan_Jomone3.f3 | 24.019 | -2.849 | internal zero-length branch |
| Japan_Jomonf0.f4 | 24.442 | -2.849 | internal zero-length branch |
| Japan_Jomonf0.f5 | 24.695 | -2.849 |  |
| Japan_Jomone5.f1 | 24.980 | -2.849 | internal zero-length branch |
| Japan_Jomone5.f3 | 24.982 | -2.849 | internal zero-length branch |
| Japan_Jomone6.f3 | 24.992 | -2.849 | internal zero-length branch |
| Japan_Jomone6.f1 | 24.993 | -2.849 | internal zero-length branch |
| Japan_Jomonf1.f2 | 24.994 | -2.849 | internal zero-length branch |
| Japan_Jomonf2.f3 | 24.998 | -2.849 | internal zero-length branch |
| Japan_Jomonf0.f1 | 25.062 | -2.849 | internal zero-length branch |
| Japan_Jomonf0.f3 | 25.064 | -2.849 | internal zero-length branch |
| Japan_Jomond1.f4 | 25.429 | -2.849 | internal zero-length branch |

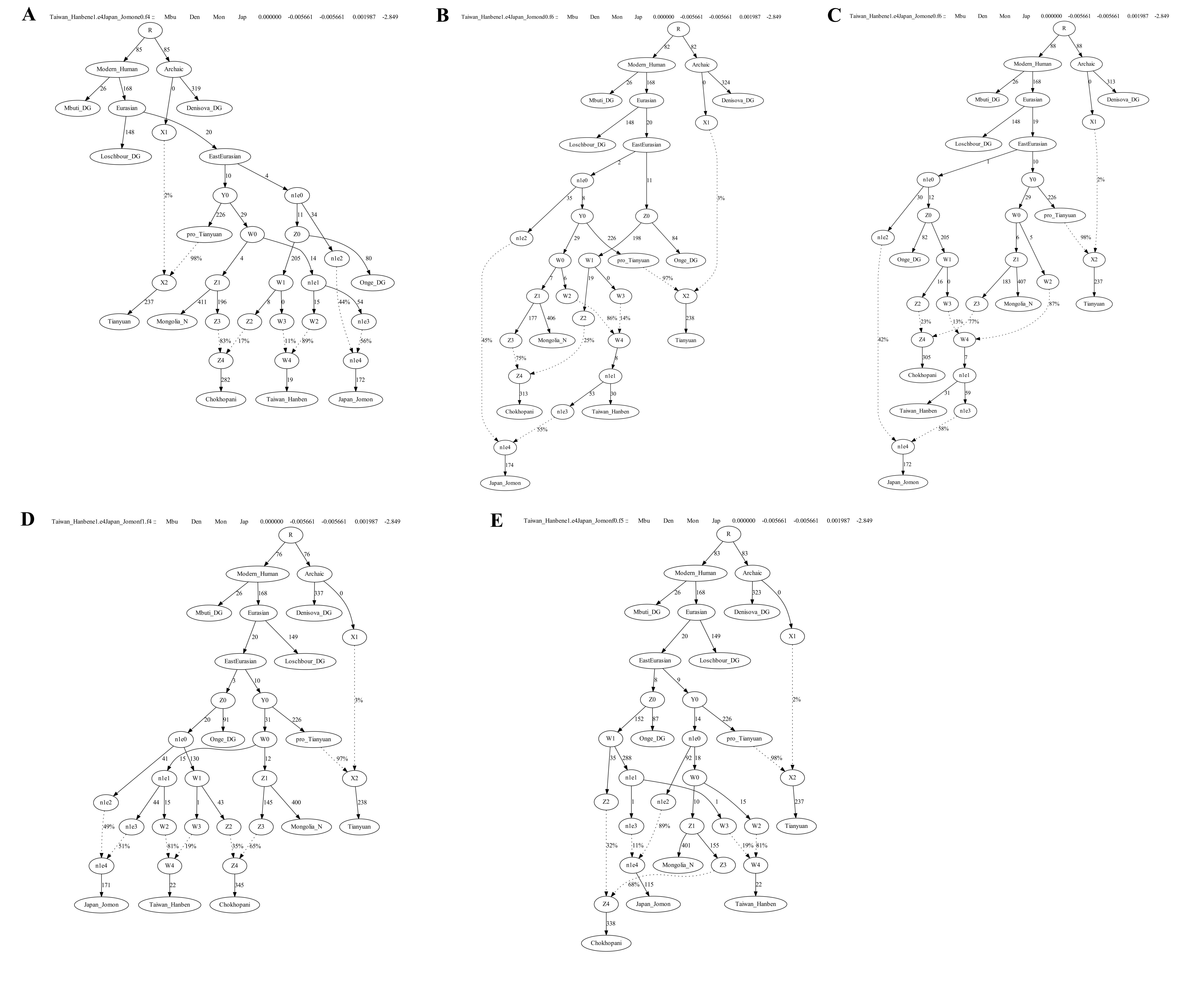
 **Figure SP17. (A) The model “Japan_Jomone0.f4” with the lowest likelihood score and Z-score by modeling Japan_Jomon is admixed;** **(B) The model “Japan_Jomond0.f6”; (C) The model “Japan_Jomone0.f6”; (D) The model “Japan_Jomonf1.f4”; (E) the model “Japan_Jomonf0.f5”. The number of transversion sites used is 105271.**

We then added Boisman onto all the possible edges of the model “Japan_Jomone0.f4” and obtained 7 fitted models as shown below in the Table SP8. There are 6 models with internal zero-length branches. We show the model “Boisman_MNe3.g5” with lowest likelihood score and Z-score without internal zero-length branches in Figure SP18.

**Table SP8. The fitted models for modeling Boisman_MN samples as admixed.**

| Model | likelihood score | worst Z score | Note |
| --- | --- | --- | --- |
| Boisman_MNe3.g5 | 23.286 | 2.848 |  |
| Boisman_MNf2.g5 | 27.511 | 2.848 | internal zero-length branch |
| Boisman_MNe5.g5 | 27.52 | 2.848 | internal zero-length branch |
| Boisman_MNf0.g5 | 30.217 | 2.848 | internal zero-length branch |
| Boisman_MNg1.g5 | 30.22 | 2.848 | internal zero-length branch |
| Boisman_MNg1.g6 | 30.829 | 2.848 | internal zero-length branch |
| Boisman_MNg1.g4 | 31.84 | 2.848 | internal zero-length branch |

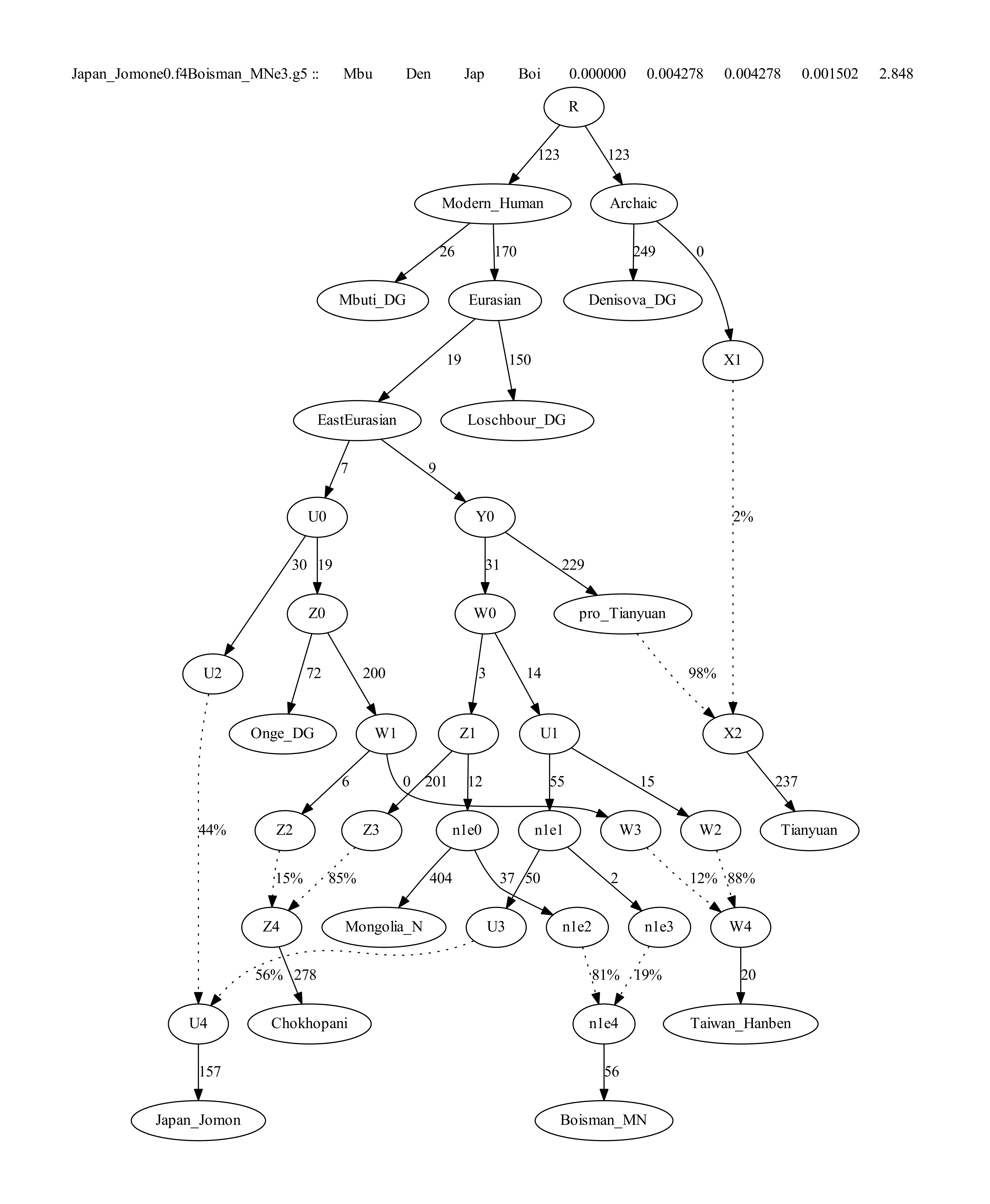

**Figure SP18: The fitted model “Boisman_MNe3.g5” with lowest likelihood score and Z-score without internal zero-length branches by adding Boisman_MN onto all the possible edges of the model “Japan_Jomone0.f4”. The likelihood score and the worst Z-score are shown in Table SP8. The number of transversion sites used is 104631.**

We have not included Wuzhuangguoliang because of the limited available number of overlapping transversions. We observed consistent results by using all sites and transversion-only datasets in exploring the phylogeny and admixture events in East Asian populations. Due to the paucity of ancient genomic data from Upper Paleolithic East Asians, there are limited constraints at present on the deep branching patterns of East Asian ancestral populations, and it is certain that this admixture graph is an oversimplification and that additional features of deep population relationships will be revealed through future work. We therefore propose this model as a null hypothesis for further testing that can help to advance future work when more ancient DNA from Pleistocene East Asians become available.
